# KineticMSI, an R-based framework for relative quantification of spatial isotopic incorporation in mass spectrometry imaging experiments

**DOI:** 10.1101/2022.08.31.505954

**Authors:** Farheen Farzana, Federico Martinez-Seidel, Anthony J. Hannan, Danny Hatters, Berin A Boughton

**Affiliations:** Florey Institute of Neuroscience & Mental Health, University of Melbourne, Melbourne, Australia; Department of Biochemistry and Pharmacology, Bio21 Molecular Science and Biotechnology Institute, The University of Melbourne, Victoria 3010 Australia; Molecular Physiology Department, Max Planck Institute of Molecular Plant Physiology, Potsdam, Germany; School of Biosciences, University of Melbourne, Melbourne, Australia; Australian National Phenome Centre, Murdoch University, Perth, Australia

## Abstract

Kinetic mass spectrometry imaging (kMSI) integrates imaging-MS with stable isotope labelling to elucidate metabolic fluxes in a spatiotemporal manner. kMSI studies are hampered by high volumes of complex data and a lack of computational workflows for data analysis that additionally address replicated experiments. To meet these challenges, we developed KineticMSI, an open-source R-based tool for processing and analyzing kMSI datasets. KineticMSI includes statistical tools to quantify tracer incorporation across replicated treatment groups spatially in tissues. It allows users to make data-driven decisions by elucidating affected pathways associated with changes in metabolic turnover. We demonstrate a validation of our method by identifying metabolic changes in the hippocampus of a transgenic Huntington’s disease (HD) mouse model as compared to wild-type mice. We discovered significant changes in metabolism of neuronal cell body lipids (phosphatidylinositol and cardiolipins) in HD mice, previously masked by conventional statistical approaches that compare mean tracer incorporation across brain regions.

## INTRODUCTION

Mass spectrometry imaging (MSI) has generated significant interest in biomedical research for its ability to spatially map the distribution and relative abundances of thousands of metabolites simultaneously within thin intact biological tissue sections in their native environment^1–4^. When used in a multimodal imaging approach, such as in combination with immunocytochemistry^5,6^, MSI allows metabolism to be examined at cell-type resolution, which can aid in understanding pathogenic mechanisms mediating the onset or progression of disease, and identifying potential therapeutic targets^7^. Typically, MSI has been used for acquiring a snapshot of an organism’s metabolism. However, when coupled to isotope labeling of tissues over time, kinetic MSI (kMSI) allows greater insight into the dynamic spatial changes in metabolism. First reported in 2013 to study phospholipid biosynthesis in a mouse tumor^8^, kMSI has since been applied in a growing number of studies in both animal and plant-based models^9–16^. kMSI generates a huge amount of data, and a lack of open-source computational tools that can automate the processing and analysis of kMSI datasets has hindered wider uptake of the method.

Currently available software for MSI users such as SCiLS Lab (Bremen, Germany), ClinPro Tools software (Bruker Daltonics GmbH, Germany), Cardinal^17^, MSiReader^18^, HIT-MAP^19^ and others are tailored for the investigation of the classical label-free MSI data from steady-state metabolomic or proteomic studies but lack features that are critical for the high-throughput analysis of stable-isotope label (SIL) data. Numerous other software pipelines such as Mass Isotopomer Distribution analysis^20^, DexSI^21^, X^13^CMS^22^, and geoRge^23^ are available for performing differential isotopic tracer labelling analysis, however these software packages have been specifically designed to support SIL data generated by traditional non-MSI approaches (i.e., gas and liquid chromatography-mass spectrometry (GC/LC-MS))^7,24,25^. While GC/LC-MS approaches are crucial for providing higher specificity and a broader metabolome coverage, these methods typically entail averaging metabolic flux across a whole tissue containing a heterogeneous population of cells, thereby compromising spatial information. Previously developed kMSI analysis pipelines do provide visualization of isotopic ratio images^10^ and spatial patterns of tracer incorporation within tissues^8,16,26^; and enable quantitative analysis of region-specific metabolism within organs^27^. However, these tools lack the statistical pipelines that allow users to conduct relative quantification of tracer incorporation between two treatment groups, such as normal versus pathophysiological circumstances, which is essential for biomedical research. In addition, it is difficult to confidently measure differential tracer incorporation between two groups, when tissue(s) display spatial heterogeneity in tracer incorporation^8,26^. The development of computational tools using freely available computational software (such as R) would aid the accessibility of data analyses pipelines. Further enhancements would be provided by tools that can streamline the entire data analysis workflow of kMSI datasets and allow users to evaluate region-specific changes in metabolic activity, which show spatially heterogenous tracer incorporation.

Here we present an open-source tool for systematically analyzing data derived from kMSI experiments, KineticMSI, which operates in R and is connected to other freely available MSI-related R packages. Key features include an automated workflow for: (1) Quality control and data pre-processing, including options to select the best tracer incorporation proxy in high isotopic quality spatial points (pixels); (2) Visualization of spatial dynamics of isotopic tracer incorporation and quick exploration of isotopic labelling patterns using unsupervised K-means clustering; (3) Coherent partitioning of replicated MSI datasets into spatial subsets comprising regions of similar tracer incorporation status and concomitant relative quantification of tracer incorporation across conditions before or after partitioning; and (4) Elucidation of significantly impacted pathways associated with the detected metabolic and proteomic changes. We applied the developed method to measure metabolic changes in a deuterium (^2^H) labelled hippocampus from a Huntington’s disease (HD) mouse model transgenic for the human huntingtin exon 1 gene fragment, versus non-transgenic wild-type (WT) littermate controls. We focused our attention on the neuron rich hippocampal subfield, Cornu Ammonis (CA1) pyramidal layer, for further analysis. The CA1 layer is vitally important for the induction of long-term potentiation (LTP) and long-term depression (LTD), mechanisms that underlie synaptic plasticity^28,29^ and hippocampal-dependent cognitive functions such as learning and memory^30^. We explored spatial heterogeneity in metabolic activity using a statistically validated unsupervised clustering approach based on ^2^H incorporation and uncovered distinct metabolic states in HD mice, where conventional statistical approaches using mean values across brain regions failed.

## RESULTS

### Experimental design to determine *in vivo* metabolic kinetics

Metabolic changes are fundamental to HD pathology. Yet it is not clear how these changes arise longitudinally as symptoms and aggregate pathology develop, and where these changes occur (i.e., which hippocampal sub-region and cell types). Here we validated our package KineticMSI by examining metabolic changes spatially within the neuron enriched CA1 hippocampal pyramidal sub-field (Fig. 1, S1) of the R6/1 mouse model of HD^31^, relative to age-matched WT mice (n=6/group). The R6/1 model involves the transgenic expression of the exon 1 gene fragment of human huntingtin containing the CAG expansion mutation, which is sufficient to cause disease-relevant pathology. To establish our dataset, we subjected WT and HD mice to isotope labeling through deuterated water following established protocols^32^, at an age corresponding to post-onset of phenotype (16 weeks) (Fig. 1). Our design was aimed at monitoring lipid synthesis by measuring the percentage of ^2^H incorporation into lipids detected by MALDI-MSI of each mouse hippocampus harvested eight days post-labelling. We selected eight days as a suitable timeframe for labelling mice as this timepoint resulted in ∼30-50% ^2^H incorporation into the metabolic targets (T_50_) across lipid classes (with at least a single substitution of ^1^H atom by ^2^H), which was sufficient to facilitate downstream statistical analysis. Around this timepoint, ^2^H concentrations in the body water have been shown to equilibrate at approximately 5% (v/v)^28^ and at this point, metabolic processes are expected to have reached a steady state. We analyzed the unlabeled lipid pools of an equal number of WT and HD mice using the classical label free-MSI approach to determine the baseline natural abundance lipid pools. In parallel, to gain a thorough understanding of the changes in ^2^H labelling in lipids found in the whole hippocampal tissue, we performed LC-MS on matched brain hemispheres (labelled and label-free) to confirm the identity of the lipid species and compare ^2^H-labelling trends achieved by kMSI.

**Fig. 1.**
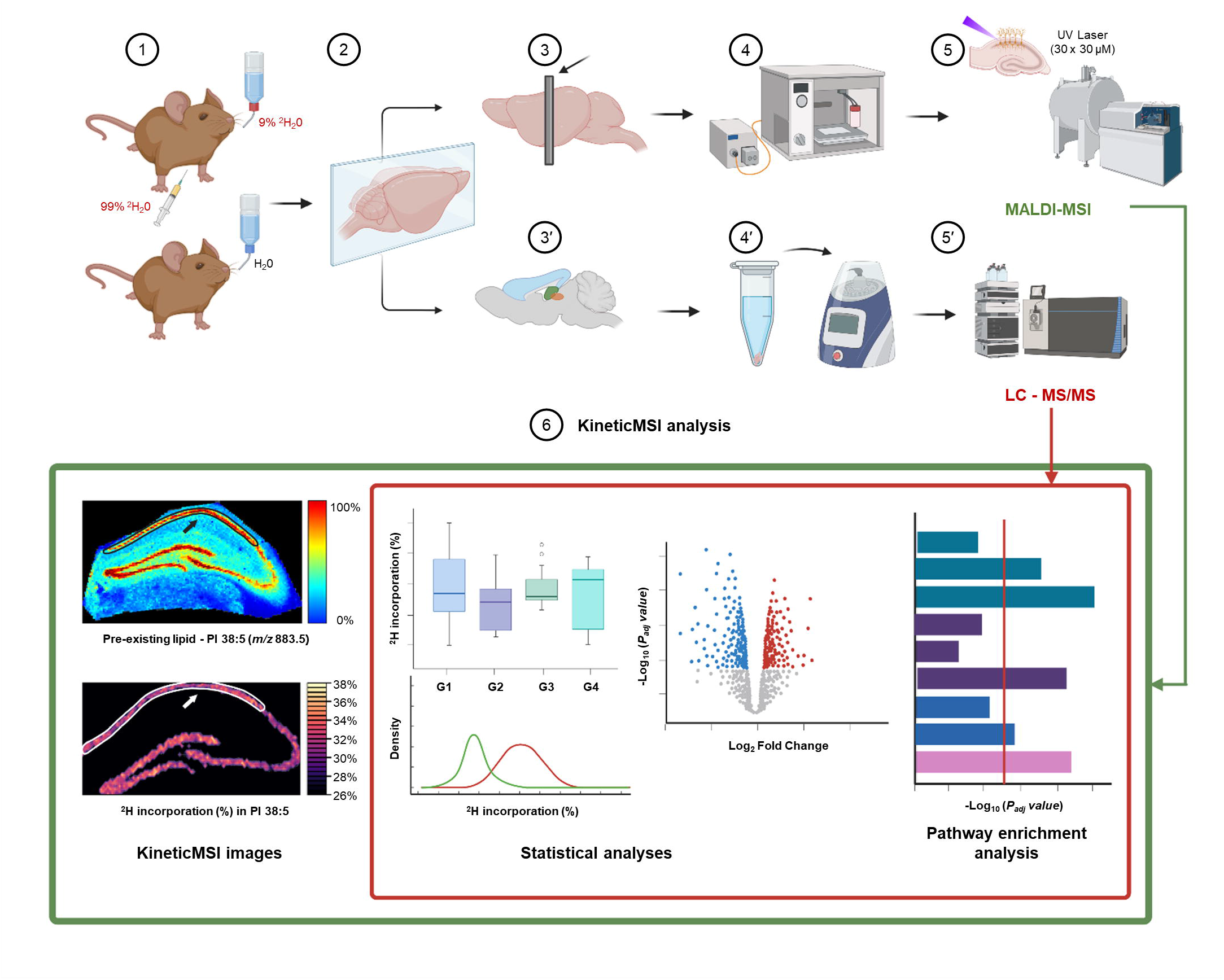
Schematic representation of the kMSI experimental workflow. (1) WT and HD mice (n = 6/group) at 16 weeks age were administered ^2^H via an intraperitoneal (IP) bolus dose and infusion of ^2^H_2_O in drinking water. Unlabeled control animals were provided free access to regular drinking water. (2) Brains were hemi-sectioned. (3) Left hemisphere was cryo-sectioned to obtain coronal hippocampal sections (20 μm). (4) Thaw-mounted sections were vacuum-desiccated followed by spray-deposition of norharmane matrix. (5) MALDI-FT-ICR-MSI, 30 × 30 µm array. (3’) Right hemisphere was dissected (hippocampus, frontal cortex, striatum) (4’) Homogenized brain regions were subjected to monophasic lipid extraction. (5’) LC-Orbitrap-MS/MS. (6) KineticMSI was used to identify lipids with differential deuterium (^2^H) incorporation in the hippocampal CA1 sub-field (shown by black and white arrowheads) of HD versus WT mice.

### KineticMSI workflow

To develop our software, several considerations were made. First, to enable users to handle the highly complex data generated by kMSI and decide the appropriate statistical approach, we designed KineticMSI to function as two modules covering different steps. The first module facilitated data quality assessment, calculation of ^2^H incorporation in a pixel-wise manner and visualization of the spatial dynamics of ^2^H incorporation within the tissue, through the reconstruction of KineticMSI images (Fig. 2a). We found that the following proxies provide the best ways to measure features of ^2^H incorporation: (1) first isotope ratio (M_1_/M_0_); (2) total isotope fraction (M_1_/(M_0_+M_1_)); (3) newly synthesized pools (corrected ∑ M_1_ + M_n_); and (4) the percent ^2^H incorporation ((corrected ∑ M_1_ + M_n_) / (corrected M_0_ + corrected ∑ M_1_ + M_n_) * 100)). Most importantly, the first module enabled selection of high-isotopic quality spatial points (i.e., pixels displaying interpretable isotopic peak profiles) and metabolic features to assess spatial differences in metabolic activity between two experimental groups.

**Fig. 2:**
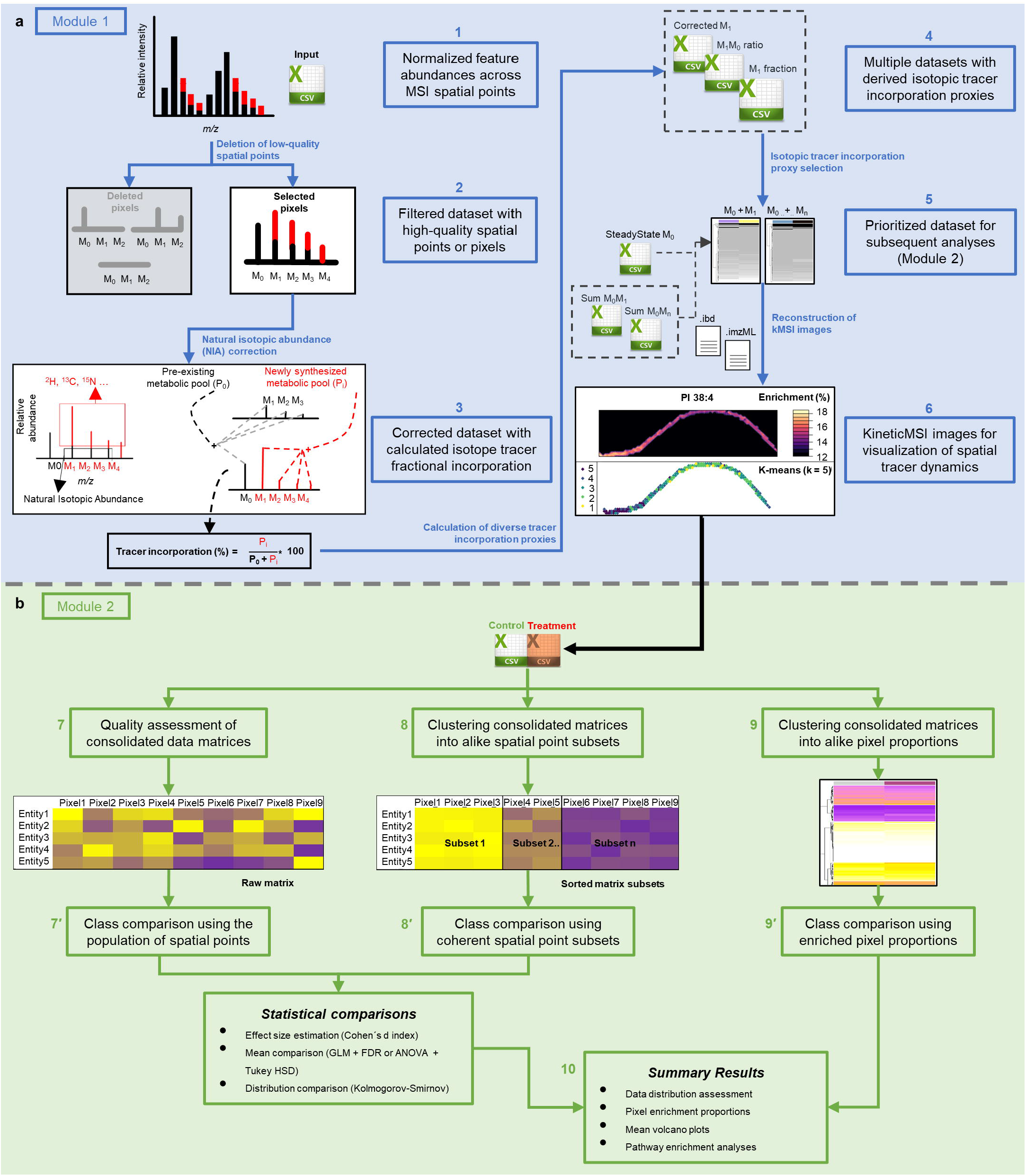
Schematic representation of KineticMSI workflow for processing and analyzing kMSI datasets. **a**, Module 1: 1, Preparation of input matrices; 2, Deletion of MSI pixels with low quality data or missing values; 3, natural isotopic abundance (NIA) correction; 4, derivation of various isotope tracer proxies; 5, definition of the most suitable isotope tracer proxies and 6, reconstruction of KineticMSI images. **b**, Module 2: 7, data quality assessment and 7’, statistical comparison using pixel population means; the second approach includes: 8, spatial segmentation into coherent pixel subsets based on tracer incorporation and 8’, statistical comparison using the pixel subset means; and the third approach includes: 9, evaluation of pixel proportions that fall under a user-defined tracer incorporation range and 9’, class comparison using enriched pixel proportions across experimental samples. 10, statistical summary of the results obtained using the above-mentioned approaches in steps 7, 8 and 9.

The second module provided statistical tools to perform relative quantification and comparative analyses of ^2^H incorporation in individual metabolic features between two samples (Fig. 2b). To enable statistical class comparison of ^2^H incorporation in targeted lipid species between WT and HD mice (n=6/group), two main approaches were used for computation of mean ^2^H incorporation (Fig. 2b): (1) ^2^H incorporation means of the complete MSI pixel population across the entire region of interest, herein termed pixel population mean, and (2) ^2^H incorporation means of coherent pixel subsets or clusters that share similar ^2^H incorporation within a region of interest, termed pixel cluster means. The clustering of MSI pixels based on ^2^H incorporation was performed using unsupervised internally validated, clustering-based approaches. The first approach may be applied to kMSI datasets that display relatively homogenous incorporation of isotopic tracer within the tissue of interest. By contrast, the second approach is suitable for kMSI datasets exhibiting intra-tissue spatial heterogeneity, and accounts for this spatial heterogeneity, prior to performing statistical comparison between treatment groups. Additionally, an extra feature allowed comparison between zones of different metabolic activity from two experimental groups, using a provision to compare pixel proportions that are below or above a pre-defined threshold of ^2^H incorporation. Finally, we performed a pathway enrichment analysis to identify significantly enriched functional categories and determine which metabolic pathways or molecular functions are associated with metabolites showing significantly altered ^2^H incorporation. For illustrative purposes, we have applied all KineticMSI tools to our exemplary dataset. To aid interpretation of the spatial data, single lipid ion images and reconstructed kMSI images were used to display differential ^2^H incorporation. Additionally, we have applied KineticMSI tools to a matching LC-MS/MS dataset from equivalent biological specimens for comparing the trends in isotope labelling obtained from kMSI datasets (Extended Data Fig. 4).

### KineticMSI application

#### Data pre-processing and spatial reconstruction of KineticMSI images

As a first step, we used KineticMSI to perform data quality control by removing MSI data pixels with missing values to ensure that they do not affect the interpretation of real spatial ^2^H incorporation dynamics in downstream calculations. Next, we corrected the data for baseline levels of natural isotopic abundance and calculated the percent ^2^H incorporation across all spatial points using the IsoCorrectoR R package^33^ (for detailed procedure, see Supplementary note 1 and Fig. S2). In the HD mouse brain dataset, we found the percentage of ^2^H incorporation i.e., the ratio of newly synthesized and total lipid pools was selected as the most suitable proxy for measuring lipid synthesis using the selection procedure outlined in Supplementary Note 2 (Fig. S3).

To visually assess the spatial dynamics of ^2^H incorporation within the tissue, we generated KineticMSI images by mapping the nominal values of ^2^H incorporation using the acquired MSI coordinates. For most lipids, we observed spatial heterogeneity evidenced by varying degrees of ^2^H incorporation across the spatial points within the tissue. The pixel-to-pixel variation in ^2^H incorporation is reflected by the dispersion of the data points in the scatterplot featuring ^2^H incorporation in PI 38:4, *m/z* 885.5 (10 – 20%) across WT and HD replicate datasets (Fig. 3a). This variation can also be visualized as color gradations in the reconstructed kMSI image of PI 38:4 in WT and HD replicated datasets (Fig. 3b). There are zones of higher ^2^H incorporation (gold) and zones of lower ^2^H incorporation (dark purple). Indeed, to gain a quick visualization of spatial patterns of metabolic synthesis, we performed K-means analysis based upon similarity in ^2^H incorporation and identified distinct clusters within the CA1 hippocampal sub-field of each MSI replicate dataset of WT and HD mice (Fig. 3c). This finding led us to utilize statistical approaches to account for the evident spatial heterogeneity in ^2^H incorporation, prior to statistical comparison of ^2^H incorporation between WT and HD mice, thus bypassing the limitation of averaging ^2^H incorporation from individual spatial points across a large tissue area, a matter that will be discussed in the next section.

**Fig. 3.**
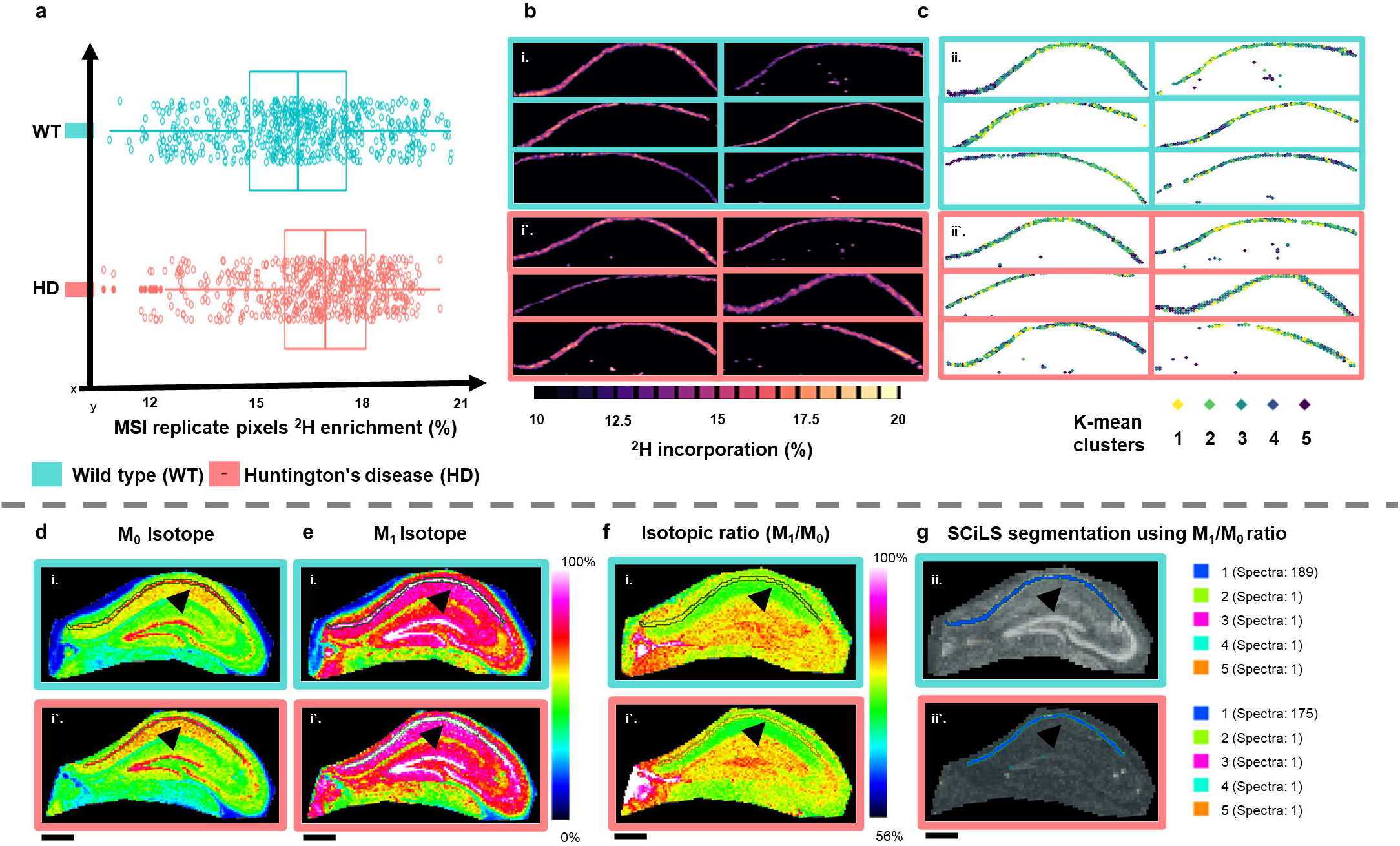
Visualization of spatial isotope labelling patterns within CA1 hippocampal sub-field. **a**, Boxplot representation of ^2^H incorporation in PI 38:4 (*m/z* 885.5) of WT and HD mice (n = 6/group). **b**, KineticMSI images depicting spatial heterogeneity of ^2^H incorporation (%) in PI 38:4 of individual WT and HD replicate datasets. Lower (closer to 0%) and higher values (closer to 100%) of ^2^H incorporation indicate decreased and increased lipid synthesis respectively. **c**, K-mean clustering (k = 5) based on ^2^H incorporation (%) in PI 38:4 through the R package ComplexHeatmap^41^ showing pixel subsets with low (yellow) to high ^2^H incorporation (purple). Intensity images of **d**, M_0_; **e**, M_1_ isotope and **f**, isotope ratio (M_1_/M_0_) for PI 38:4 of a representative WT-HD replicate pair using SCiLS Lab software. **g**, K-means clustering (k = 5) based on M_1_/M_0_ image of PI 38:4 using SCiLS Lab software, showing top five k-mean clusters and number of spectra in each cluster (right). Scale bar – 500 µm.

We benchmarked our results against SCiLS Lab (Fig. 3d-g), where we visualized the intensity image of M_0_ (Fig. 3d), M_1_ (Fig. 3e) and the isotope ratio image (i.e. M_1_ feature normalized to its M_0_ *m/z* feature, M_1_/M_0_) for PI 38:4 (Fig. 3f) for determining ^2^H incorporation in PI 38:4. As suggested by the isotope ratio (M_1_/M_0_) image for PI 38:4 (Fig. 3f), we found spatial heterogeneity in ^2^H incorporation reflected by the color gradations within the CA1 sub-field of WT and HD tissues (black arrows). In contrast to our results, segmentation of the hippocampal CA1 sub-field using K-means analysis in SCiLS Lab was unable to reveal any distinct spatial patterns based on the isotope ratio (M_1_/M_0_) for PI 38:4 in WT and HD mice (Fig. 3g).

### Differential analysis of ^2^H incorporation between WT and HD mice

To statistically compare ^2^H incorporation in targeted lipid species between WT and HD mice, we used two approaches that include: mean comparison using (1) pixel populations and (2) coherent-clustered pixel subsets that share similar ^2^H incorporation.

### Comparison using pixel population mean reveals no difference in ^2^H incorporation between WT and HD mice

To calculate ^2^H incorporation pixel population means, we first addressed the challenge of variability in the number of pixels across individual kMSI replicates by randomly sampling a matching number of pixels equal to the pixel number of the smallest dataset (For the procedure used to assess the correctness of random sampling approach, see supplementary note 3 and Extended Data Fig. 1b for details). We then evaluated data distributions of ^2^H incorporation across the selected MSI pixels of the CA1 hippocampal sub-field (See supplementary note 3 and Extended Data Fig. 1a, b and d for details) and compared pixel population means i.e., mean ^2^H incorporation in the target lipids across the entire CA1 hippocampal sub-field. This analysis revealed no significant changes in mean ^2^H incorporation between WT and HD mice (Fig. 4c, bottom left), demonstrated using the neuronal lipid PI 36:4 (*m/z* 857.5) (Generalized linear models, FDR-adjusted P value = 0.93) (Fig. 4a). However, when we compared the shapes and the extent of overlap of the distribution of ^2^H incorporation in PI 36:4, we found significantly different distributions between WT and HD mice, as evident by a rightward shift in the cumulative frequency plot of HD (red) compared to WT mice (blue) (Kolmogorov-Smirnov test, FDR-adjusted P value = 0.01 and Cohen’s d value = 0.76) (Fig. 4b). This significantly altered distribution in ^2^H incorporation was observed across the majority of neuronal cell body enriched lipids such as PI 38:4 and PI 38:5 (*m/z* 883.5) and synaptic lipids such as PA 34:1 (*m/z* 673.48) and GM1 36:1 (*m/z* 1544.8), indicative of a significantly higher ^2^H incorporation in HD mice (Fig. 4c, bottom right). This scenario suggested that averaging ^2^H incorporation across pixels displaying spatial heterogeneity in ^2^H labelling could potentially mask significant differences in ^2^H incorporation between WT and HD mice. The pathway enrichment analysis based on lipids that significantly change their distributions, identified neuronal cell body lipids (Fisher’s exact test P value = 0.001) as a significantly enriched functional category. This category is associated with lipid features displaying a trend towards higher ^2^H incorporation in HD mice (Fig. 4d). Furthermore, we observed a shift from a bimodal distribution in ^2^H incorporation in WT mice to a unimodal distribution in HD mice (Fig. 4b) that led us to hypothesize the existence of distinct clusters or sub-populations of pixels/cells with similar ^2^H incorporation within the CA1 hippocampal sub-field of the mouse hippocampus.

**Fig. 4.**
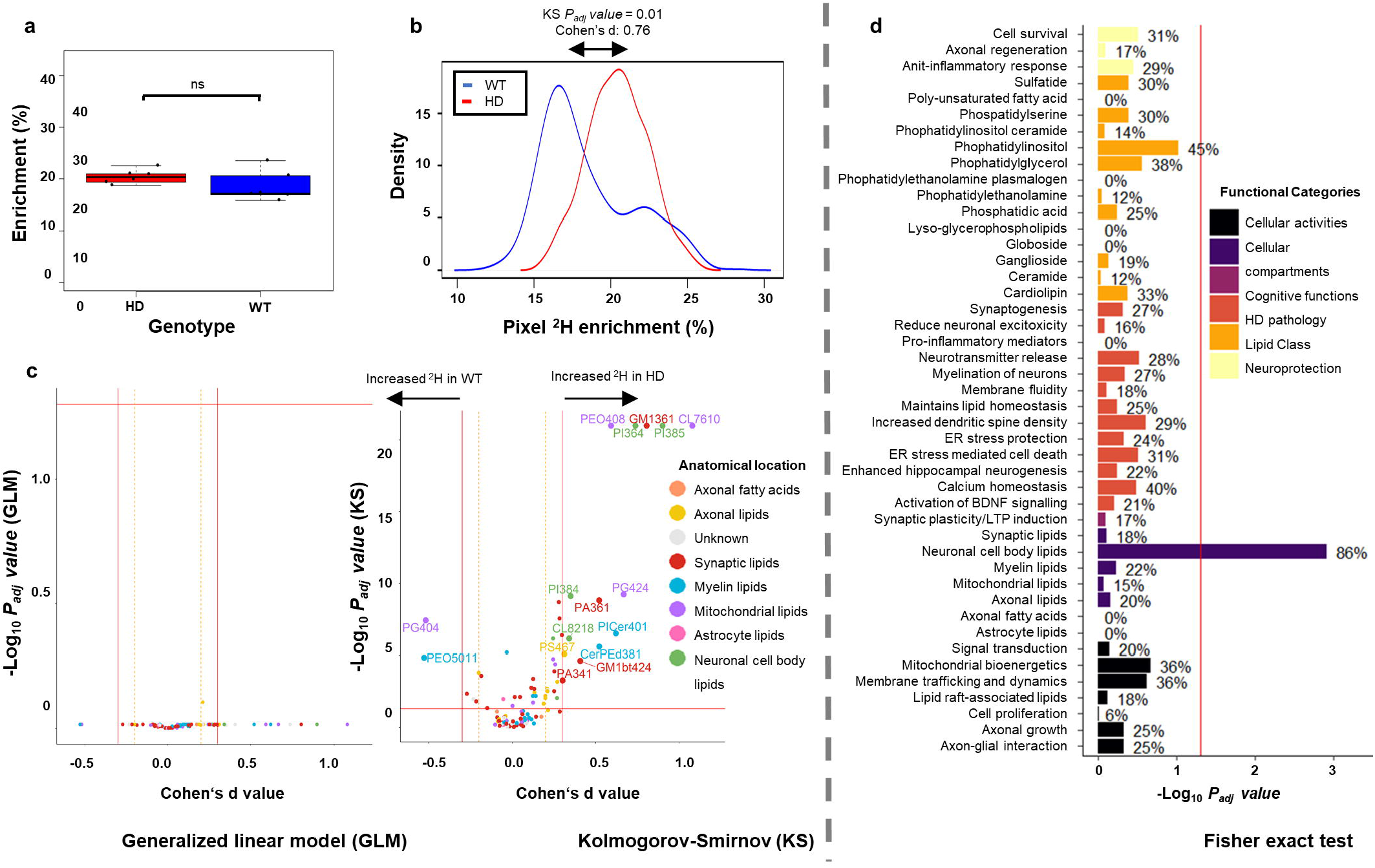
Differential analysis of ^2^H incorporation in brain lipids of WT and HD mice using pixel population means. **a**, Boxplot representation showing no significant difference between mean ^2^H incorporation in PI 36:4 (*m/z* 857.5) between WT and HD mice (n=6/group) using GLM. **b**, Kolmogorov-Smirnov (KS) test and effect size estimation (Cohen’s d values) of ^2^H incorporation in PI 36:4 showing significantly different ^2^H incorporation of WT (blue) and HD mice (red). **c**, Volcano plot – bottom left (Cohen’s d values (X-axis) versus -log_10_ FDR-adjusted P value from GLM (Y-axis)) showing no significant difference in mean ^2^H incorporation in lipid features WT versus HD mice. Volcano plot – bottom right (Cohen’s d values (X-axis) versus -log_10_ FDR-adjusted P value (Y-axis) using KS test) showing significantly different distributions of ^2^H incorporation in lipid features of WT and HD mice. Cohen’s d values ≥ 0.3 indicate differentiation of WT and HD mice. Each dot represents individual lipid feature coloured by known anatomical location in mouse hippocampus. Grey dots represent features that have no defined category in the custom-built library provided for pathway enrichment analysis (Table S3). Significantly changed features are highlighted and labelled. **d**, Bar plot representation showing the results of pathway enrichment analysis (-log_10_ FDR-adjusted P value from the Fisher exact test (X-axis) versus functional categories associated with the detected lipid features in our study (Y-axis)). Functional categories are colored by lipid class, cellular activity, compartment, cognitive function, known HD pathology and neuroprotection to enhance interpretation of results (BDNF – Brain-derived neurotrophic factor; LTP – Long-term potentiation). Values denotes the proportion of lipids with significantly altered ^2^H incorporation in each functional category. Red line denotes significance P < 0.05.

### Clustering analysis reveals significant differences in ^2^H incorporation between WT and HD mice

To test for the presence of pixel subsets sharing similar ^2^H incorporation within the hippocampal CA1 sub-field, we performed tissue segmentation based upon similarity in ^2^H incorporation in an unsupervised manner, independently within the WT and HD groups (n=6/group). This revealed the presence of two coherent pixel clusters of ^2^H incorporation (AU-P value ≥ 0.95) in the hippocampal CA1 sub-field, corresponding to the ‘low’ and ‘high’ ^2^H enrichment zones (Extended Data Fig. 2a). The partitioning into two coherent clusters of MSI pixels based on the degree of ^2^H incorporation was observed in the majority of lipid features, as confirmed by the density histogram summarizing the number of significant clusters found across the evaluated lipid features (Extended Data Fig. 2b). The cluster dendrogram of the lipid PI 38:4 (*m/z* 885.5) shows two significant pixel cluster subsets (AU-P value = 0.95) obtained by sub-setting the kMSI data structure based on the extent of ^2^H incorporation (Extended Data Fig. 2c).

In addition, we visualized spatial patterns of metabolic synthesis by mapping the obtained significant clusters subsets onto the original MSI images and found areas of preferentially high versus low ^2^H incorporation that potentially reflect the presence of “metabolic hotspots” versus “metabolically inactive” areas within the CA1 hippocampal sub-field of WT and HD mice.

These metabolic patterns were particularly evident in the neuron-enriched lipid PI 38:4, which showed spatial constraints in ^2^H incorporation with the high ^2^H enrichment zones (yellow) localized at the edges of the hippocampal CA1 field in the WT mice. On the other hand, the HD mice showed a more dispersed distribution of ‘high’ and ‘low’ ^2^H incorporated pixels, suggesting a potential loss of spatial coordination in lipid synthesis in HD mice (Extended Data Fig. 2d). These results confirm the intra-tissue spatial heterogeneity in ^2^H incorporation as observed in PI 38:4 (Fig. 3a) and implies the presence of cellular sub-populations with their own distinct ^2^H incorporation dynamics into the target lipid pools within the CA1 hippocampal sub-fields of WT and HD mice.

Next, using the coherent pixel clusters obtained from the above clustering analysis, we compared mean ^2^H incorporation from each cluster pair of individual lipid features between WT and HD mice. This analysis identified lipids showing significant differences in ^2^H incorporation in majority of the neuronal cell body enriched lipids such as PI 36:4 (*m/z* 857.5), PI 38:5 (*m/z* 883.5) and PG 44:12 (*m/z* 865.5) and synaptic lipids such as CerP 23:3 (*m/z* 788.5) and PA 36:4 (*m/z* 695.5) in HD mice, relative to WT controls (Fig. S4a). Using the lipid PI 36:4 as an example, Fig. 5a highlights the multiple comparisons performed between the cluster pairs of WT and HD mice. We found a significant increase in ^2^H incorporation in PI 36:4 in HD mice, when the comparison was performed using both pixel cluster means using One-way ANOVA followed by post-hoc Tukey HSD test (Fig. 5b) and distribution of ^2^H incorporation across MSI pixels between WT and HD mice (Kolmogorov-Smirnov test, FDR-adjusted P value = 0.02 and Cohen’s d value = 2.26) (Fig. 5c). Therefore, addressing spatial heterogeneity in metabolic activity, prior to statistical comparison of ^2^H incorporation between WT and HD mice revealed distinct clustering patterns and significant changes in ^2^H incorporation in lipids of HD mice, compared to WT controls (Fig. S4a) that were otherwise masked by comparing overall population means of ^2^H incorporation across the entire CA1 hippocampal sub-field (Fig. 4c, bottom left).

**Fig. 5.**
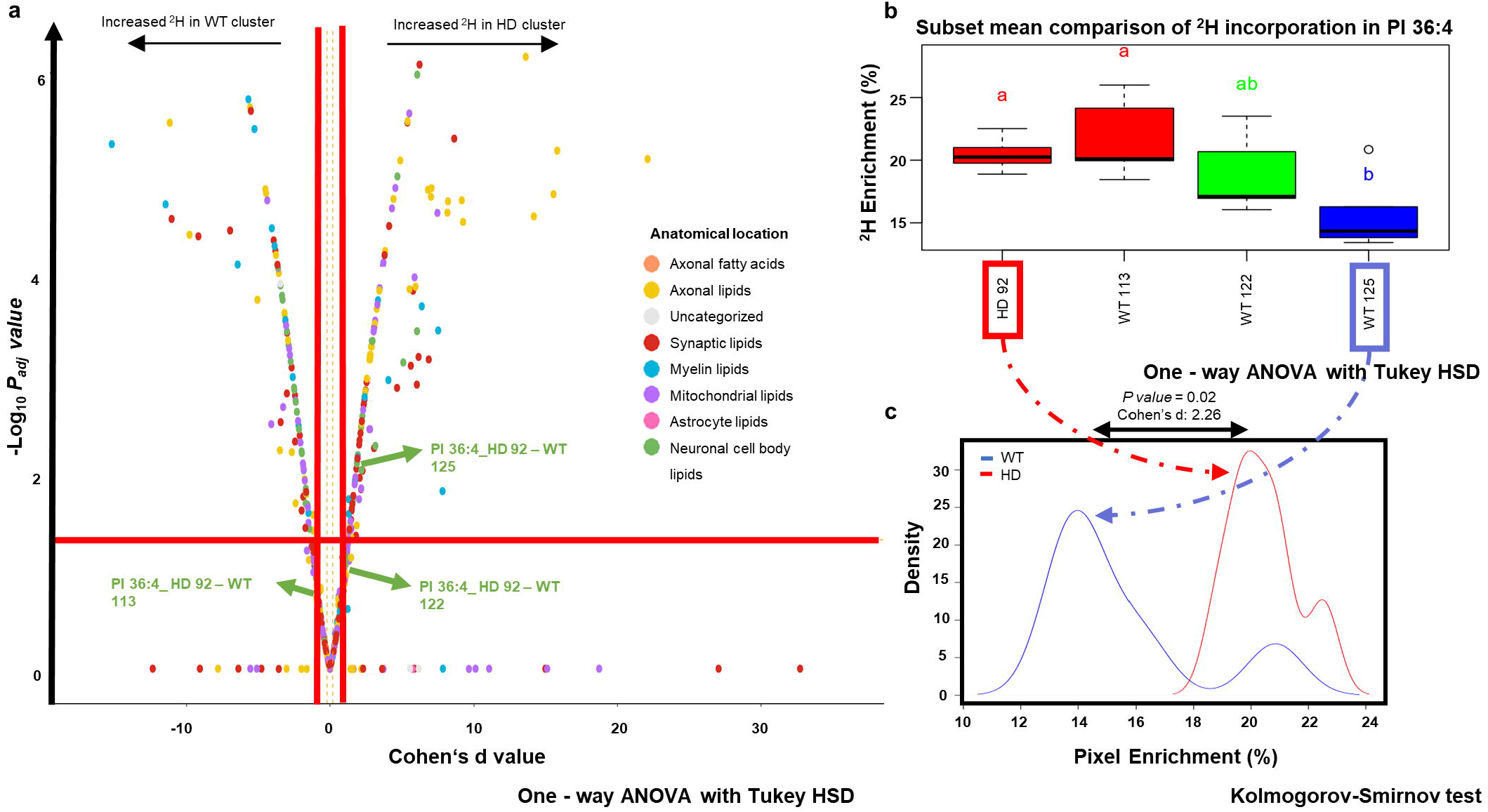
Differential ^2^H incorporation in lipids of WT and HD mice using pixel cluster means. **a**, Volcano plot (Cohen’s d values (X-axis) versus -log_10_ FDR-adjusted P value from GLM (Y-axis)) showing mean ^2^H incorporation in PI 36:4 (*m/z* 857.5) WT versus HD mice (n = 6/group). Cohen’s d values ≥ 0.3 indicate differentiation of WT and HD mice. Each dot represents the individual cluster pixel subset comparison from PI 36:4 coloured by known anatomical location in mouse hippocampus. Grey dots represent features that have no defined category in the custom-built library provided for pathway enrichment analysis (Table S3). Significant cluster pair showing differential ^2^H incorporation highlighted in bold **b**, Boxplot representation showing multiple pairwise comparison of pixel subset means ^2^H incorporation in PI 36:4 between WT and HD mice (n=6/group) using One-way ANOVA followed by Tukey HSD test. **c**, Kolmogorov-Smirnov test and effect size estimation (Cohen’s d values) of ^2^H incorporation in PI 36:4 showing significantly different pixel subset pair (HD_92 versus WT_125) of WT (blue) and HD mice (red).

### Exploration of ‘metabolic hotspots’ reveals distinct metabolic states in HD mice

Next, we explored metabolically active zones by comparing proportion of pixels with ‘high metabolic activity’ i.e., pixels exhibiting at least half of the total ^2^H incorporation detected (i.e., >15%) between WT and HD mice. The outcome of this analysis represented by a graphical heatmap (Extended Data Fig. 3) showed significantly higher proportion of ^2^H enriched (≥ 15%) in the majority of lipid features of HD mice, relative to WT controls. This shift towards higher metabolic activity state in HD mice is in agreement with the significant increase in ^2^H incorporation observed in HD mice, relative to WT controls using the above-mentioned statistical approaches. Hence, our results provide a strong basis for exploring pixel subsets sharing similar tracer incorporation between two groups of interest, prior to statistical comparison of tracer incorporation across the entire tissue to avoid misinterpretation of the statistical results.

## DISCUSSION

Here we describe KineticMSI, an open-source automated R-based pipeline to process and analyze the enormous amount of data generated isotope-labelled MSI data. We applied KineticMSI to show how different lipid metabolites changed their apparent synthesis rates in the hippocampus of HD mouse model, relative to control mice. KineticMSI package has the potential to work with any isotope labelled MSI data such as ^2^H, ^13^C, ^15^N, ^18^O and ^34^S. Our workflow incorporates a range of statistical tools to conduct relative quantification of isotopic tracer incorporation into common biomolecules (proteins, metabolites, lipids) and compare tracer incorporation between different treatment groups containing multiple kMSI replicate sets that display intra-tissue spatial heterogeneity. This work is significant as it is the first to address the challenges posed by the need for replication in kMSI studies. Furthermore, KineticMSI allows users to take data-driven decisions by providing a tool for the elucidation of significantly perturbed pathways, thus providing an in-depth assessment of the detected metabolic turnover changes and avenue to gain mechanistic insights into (disease) biology in a wide range of systems.

While previous methods described for analyzing ^13^C or ^2^H-MSI experiments potentially fulfill the need to examine distinct isotopic labelling patterns spatially *in vivo* ^8,16,27^, the originality of our method relies on its features that facilitate statistical comparison of tracer incorporation between experimental groups (containing multiple replicate datasets) displaying regional spatial heterogeneity in tracer incorporation. This feature not only allows users to investigate region and sub-region-specific changes in metabolic activity across different biological systems but also paves the way toward understanding metabolic synthesis shifts under pathological conditions. Moreover, while tissue segmentation maps provided by the available MSI-data analysis software such as SCiLS Lab (Bremen, Germany), Cardinal^17^ and others are based on spatial distribution of metabolites, KineticMSI complements these existing methods by providing additional features to perform segmentation based on a user-defined isotope tracer incorporation proxy and downstream statistical analysis. While SCiLS Lab allows for the visualization of isotope ratio (M_1_/M_0_) images, it does not compute the percent ^2^H incorporation which takes into the account all the detected labelled isotopic peaks (M_1_, M_2_, …M_n,_) (see supplemental note 2 and Fig S2 for further explanation on the reason behind why isotope ratio (M_1_/M_0_) is not ideal for calculation of ^2^H incorporation). Moreover, SCiLS Lab permits visualization of isotope ratio (M_1_/M_0_) images on an individual basis for each metabolic feature using single ion normalization (M_0_) and hence does not permit high-throughput analysis of kMSI datasets. Indeed, the inability to reproduce the segmentation results using SCiLS Lab (Fig. 3g) and Cardinal^17^ (Fig. S5a) was likely due to the limitations related to the pre-processing of kMSI datasets by these methods that include: (1) Lack of a feature to filter out pixels with missing values from either the monoisotopic peak (M_0_), the isotopologue peaks (M_1_, M_2_, …M_n,_), or both; (2) Omission of the correction for natural isotopic abundance for calculating ^2^H incorporation; and (3) Inability to compute the percent ^2^H incorporation.

Furthermore, the bootstrapped HCA approach - pvclust^34^ implemented in our workflow is superior to conventional K-means clustering algorithms as it computes a statistical measure i.e., Approximately unbiased probability values (AU-P values) for each cluster and only returns the most robust and significantly valid clusters that satisfy the significance threshold i.e., the AU-P value. This feature permits simultaneous visualization and identification of isotopic labelling patterns of several target biomolecules such as proteins, metabolites and lipids within the tissue, thus allowing researchers to capture *in vivo* kinetics of metabolic or protein synthesis in a high throughput manner.

In this work, we validated our method by discerning metabolic changes in samples obtained from the hippocampus of a Huntington’s disease (HD) neurodegenerative disease mouse model, compared to wild-type (WT) mice. We first generated KineticMSI images to visualize ^2^H incorporation dynamics in a pixel-wise manner and found spatial heterogeneity in ^2^H incorporation in the examined lipids across the MSI pixels of CA1 hippocampal sub-field of both WT and HD mice. The pixel-to-pixel variability in ^2^H labelling is not due to the differences in concentration or ionization efficiency of the features examined, and the percent ^2^H incorporation is independent of the absolute abundances of monoisotopic peak (M_0_) and the labelled isotopologue peaks (M_1_, M_2_, …M_n,_), but rather depends on their ratios. This suggests the presence of metabolically heterogeneous cellular sub-populations within the CA1 hippocampal sub-field. This is not surprising, given the complex cellular heterogeneity of the brain, characterized by the presence of multiple neuronal and non-neuronal cell-types (including sub-types) with diverse functional and metabolic characteristics^35,36^. In agreement to our study, spatial heterogeneity has also been reported in phospholipid synthesis within the mouse tumor tissue by previous kMSI-based studies^8^.

Although the presence of metabolically heterogeneous cellular sub-populations within a tissue of interest adds a new level of complexity for data interpretation, in the current work, we present clustering approaches that extract distinct labelling patterns to account for regional heterogeneity in tracer incorporation, prior to statistical comparison of tracer incorporation between two conditions. Indeed, by addressing spatial heterogeneity in ^2^H incorporation in the examined lipids of WT and HD mice, we uncovered distinct metabolic states with significantly higher ^2^H incorporation in the lipids of HD mice, relative to WT controls, that failed to be revealed by comparing the overall mean ^2^H incorporation across the entire hippocampal sub-field. These findings highlight the importance of acknowledging the presence of intra-tissue spatial heterogeneity in isotopic tracer incorporation in relatively homogenous regions which, when unaccounted for, can potentially obscure significant differences in tracer incorporation between two treatment groups, which could mislead statistical analysis and lead to incorrect interpretation of biology. Indeed, the LC-MS study confirmed a significantly higher ^2^H incorporation in majority of neuronal lipids in the hippocampal tissue of HD mice, relative to WT controls (Extended Data Fig. 4a), thus reproducing the trend observed in the kMSI study. The higher number of significantly impacted functional categories reported by LC-MS represent the global changes measured across the entire hippocampal region and are in contrast to the changes observed in the CA1 hippocampal sub-field measured using kMSI, thus confirming the loss of sub-field specific changes in metabolic activity measured using MSI (Extended Data Fig. 4c).

The modular design and multi-step analysis in the KineticMSI workflow provides maximum flexibility to the users to optimize strategies and parameters at different stages of the data analysis workflow to suit the needs of the system under investigation. Moreover, the entire workflow has been written using base R objects and classes, with some method dependencies to S3 and S4 packages. This allows users to avoid executing the entire workflow every time, by bypassing some of the functions if their data is already in an optimal state.

Although we have used isotopic labelled data from a single time-point for illustration purposes, the same workflow can be readily applied to analyze time-series kMSI datasets to perform metabolic flux analysis. By default, the filtering parameters cater to a partially labelled state since a short isotope labelling period of eight days was followed in the illustrated example dataset. Nevertheless, we have implemented a parameter that allows users to apply the statistical workflow to a fully labelled state. Moreover, KineticMSI package is generally applicable to isotope labelled data generated from traditional metabolomic approaches such as GC/LC-MS (For details on formatting input tables, see Methods section).

Prior to the implementation of the kMSI workflow, we recommend assessing the separability of the tissue of interest by applying existing spatial segmentation approaches such as SCiLS Lab (Bremen, Germany) (Fig. S1b) and Cardinal^17^ to segment the tissue into appropriate spatial patterns based on biomolecular compositions. This not only serves to reduce the complexity of the data but also enables statistical comparison between matched segments (i.e., similar tissue and cell-types) from two groups of interest^37^, thus accelerating the subsequent downstream analysis. However, this is not a necessity for the KineticMSI workflow. Also, we have implemented a standardized batch-effect correction algorithm i.e., ComBat correction^38^ for correcting the raw data, in this case applied to steady-state lipid pools; however, there is a provision for users to apply a normalization method of their choice and generate the input files in the correct format for further analysis.

One possible limitation of our package is that the bootstrap clustering algorithm used to segregate pixels based on tracer incorporation may result in arbitrary partitioning of the data, where pixels with highly similar tracer incorporation can be incorrectly classified into different clusters. To overcome this issue, we provide the feature to perform cross-validation of the clusters obtained from the clustering algorithm by comparing cluster means using one-way ANOVA and Tukey HSD post-hoc testing (Fig. 5B).

Taken together, our results caution against the use of pixel population mean comparisons of tracer incorporation across entire tissues. In order to make valid statistical comparisons of metabolic activity between two conditions, we recommend addressing any spatial heterogeneity in tracer incorporation prior to statistical analysis to facilitate correct downstream data interpretation. KineticMSI provides a comprehensible guide for both bench biologists and computational scientists, thus enabling a broader scientific community to take advantage of the method to analyse kMSI datasets and capture the rapid and dynamic metabolic and proteomic changes associated with healthy and pathological states. In the future, this tool can serve as a valuable resource to accelerate both fundamental and clinical research by facilitating the investigation of biomarkers for early detection of diseases in a range of medical fields, as diverse as cancer, neurodegenerative disease, cardiovascular and immune dysfunctions, parasitology, and plant biology, all of which have been associated with widespread perturbations in metabolic processes. Hence, future work focused on improving collaboration between biologists and computational scientists could pave the way for the development of user-friendly tools that will allow us to better interpret the rich biological data provided by SIL studies and advance our understanding of both normal physiology and the pathophysiology of many diseases.

## ONLINE METHODS

### Experimental workflow

The experimental design for generating kMSI dataset begins with the introduction of deuterated water (99 atom% Deuterium oxide (^2^H_2_O), Sigma-Aldrich and 0.9% (w/v) NaCl) via an intraperitoneal injection bolus of 35 µl/gm (body weight), followed by a maintenance dose of 9% (v/v) deuterated water in drinking water, in HD mice and age-matched WT controls (n = 6/group) at 16 weeks of age. Mice were euthanized 8 days post-labelling and the brain tissue was rapidly collected and hemi-sectioned (∼ 3 – 5 minutes). While one brain hemisphere (left) was flash-frozen on liquid nitrogen and used for MALDI-MSI, the other matched hemisphere (right) was dissected to obtain the frontal cortex, hippocampus, and striatum, which were homogenized, extracted, and examined in detail using Liquid Chromatography (LC)-MS/MS analysis. The left hemisphere was then cryosectioned at 20 μm thickness, followed by the deposition of Norharmane matrix (For details on animal care, sample preparation and tissue collection, see Supplementary note 1-3). Data acquisition of MALDI-MSI was carried out using a Bruker SolariX 7T XR hybrid ESI–MALDI–FT–ICR–MS platform at an estimated resolving power of 130,000 at *m/z* 400 in the negative ionization mode (see Supplementary note 4 for details). Data processing and multivariate analysis of MALDI-MSI data according to a series of tests as outlined in Supplemental Note 5 and 6. The hippocampi from the matched brain hemispheres (right) were homogenized and analyzed using LC-MS/MS operated in the negative and positive ionization mode. Species level lipid annotation for MSI were derived from LC-MS/MS molecular species level annotations (For details on lipid extraction process, acquisition, pre-processing and analysis of LC-MS data, see Supplementary).

### Preparation of input matrices for KineticMSI

Prior to applying the KineticMSI workflow, we performed data pre-processing using SCiLS Lab software (see supplementary for details). Subsequently, we exported the data matrices (.csv files) containing normalized intensities of all mass features including monoisotopic (M_0_/A_0_) and labelled isotopologue peaks (M1, M2…Mn, where n is the number of nominal mass units added to the monoisotopic mass based on the detected labelled isotopes) from SCiLS Lab. The paired *.ibd and *.imzML files were also exported to obtain the file coordinates for generating KineticMSI images. Additionally, data matrices containing batch-effect corrected signal intensities of the monoisotopic (M_0_) peaks of the matched mass features were also exported from unlabeled controls to determine the most suitable proxy for measuring ^2^H incorporation (see details in Supplementary Note 2 and Fig. S3).

### Kinetic MSI workflow

All the analysis outlined below was performed using the R package KineticMSI (https://github.com/MSeidelFed/KineticMSI)

#### Deletion of missing values

In the illustrated example i.e., a partially labelled dataset, pixels that lack either the monoisotopic peak (M_0_/A_0_ depending on fragmentation), the isotopologue peaks (M_1_, M_2_, …M_n,_), or both were filtered out. Hence, the standard KineticMSI implementation treats pixels that lack an M_0_/A_0_ signal but have a detected signal intensity in its isotope envelop as an artifact, since in a partially labelled state the M_0_/A_0_ peak is not expected to disappear due to complete mass shifts to the M_1_, M_2_, …M_n_ isotopologues (Fig. S2c). However, in a fully labelled state, complete disappearance of M_0_/A_0_ accompanied by an increase in the signal of its labelled isotopic envelope is possible (Fig. S2b). Thus, a parameter in the implemented R function was included to allow users to either (1) delete pixels that only lack M_1_…M_n_ isotopologues (applicable to fully labelled states) or (2) delete pixels that lack both, M_1_…M_n_ isotopologues and the monoisotopic peak signal (applicable to partially labelled states).

#### Calculation of isotope incorporation

Natural isotope correction and calculation of the percent ^2^H incorporation were performed in a pixel-wise manner by adapting functions from IsoCorrectoR^33^, an R-based package. The percent ^2^H incorporation was calculated as the ratio of newly synthesized (^2^H - labelled) and total lipid pool (newly synthesized + pre-existing lipid pools) i.e., ((corrected ∑ M_1_ + M_2_ + … M_n_) / (corrected M_0_ + corrected ∑ M_1_ + M_2_ + … M_n_)) * 100), where n is the number of extra atomic mass units added to the monoisotopic mass based on the detected labelled isotopes. A cross-validation and an alternative function were implemented using IsoCor^39^, a Python-based module, to confirm the equivalent percentages of background-corrected ^2^H incorporation. To select the most appropriate isotopic ^2^H proxy, batch-effect correction of the steady state pools from non-labelled controls was performed using ComBat correction^38^ as detailed in the SVA package^40^, followed by its comparison to the ^2^H-labelled metabolite steady state pools.

#### Visualization and spatial segmentation based on ^*2*^*H incorporation*

To recreate kineticMSI images, graphical reconstructions of the MSI images for each metabolite feature were built by mapping ^2^H incorporation values onto the original coordinate system obtained from the MALDI-MSI platform. To extract the file coordinates from the acquired MSI images, KineticMSI functions use a Cardinal^17^ dependency, which is an R package designed to perform statistical analysis on MSI datasets. To further explore spatial patterns of metabolic synthesis, MSI pixels were segregated based on similarity in ^2^H incorporation independently for WT and HD kMSI datasets. Segregation was done using two unsupervised clustering approaches that include: (1), K-means algorithm (with a user-defined k value = 5) through the R package ComplexHeatmap^41^; and (2), Hierarchical cluster analysis (HCA) via multiscale bootstrap resampling to return an optimized number of significant clusters that are above a user-defined significance threshold (Approximately unbiased probability (AU-P)), which is a dependency from the R package pvclust^34^. For the HD mouse brain dataset, the AU-P value and the bootstrap iteration number ‘nboot’ were set to 0.95 (95% confidence) and 1000 iterations respectively to improve robustness and confidence in the resulting clusters.

#### Statistical approaches used for the relative quantitation of ^2^H incorporation

To perform differential analysis of ^2^H incorporation between WT and HD mice using both population and cluster means, two approaches for class comparison were used that include: (1), One-way analysis of variance (ANOVA) followed by Tukey HSD (Honestly significant difference) post-hoc testing; and (2), a parametrized Generalized linear model (GLM), according to the procedure detailed in the R package RandoDiStats (https://github.com/MSeidelFed/RandodiStats_package). As an alternative to mean comparisons, the shapes of the empirical cumulative distributions of ^2^H incorporation were compared between WT and HD datasets using the two-sample Kolmogorov-Smirnov test. Complementarily to class and distribution comparison, an effect size estimation using the Effsize^42^ R package was employed to obtain Cohen’s d values that measure the extent of overlap between the distributions of WT and HD mice. Cohen’s d statistic is used to indicate the standardized difference between two means (difference between two means divided by the pooled standard deviation). Unlike ANOVA test, effect size calculations are independent of sample size, thus preventing overestimation of the significance of differences between the large number of individual spectra (pixels) collected in MSI experiments^43^. Cohen’s d absolute values of 0.1, 0.2 and 0.3 were set as thresholds corresponding to a small, medium, and large effect size respectively, based on recommendations from Gignac and Szodorai (2016)^44^. In metabolic turnover studies, small and medium effect sizes (Cohen’s values < 0.3) are indicative of perturbations in metabolic synthesis which may have major implications for lipid homeostasis and be highly associated with disease phenotypes.

Additionally, a generalized linear model was used to compare pixel proportions below or above a pre-defined magnitude threshold of ^2^H incorporation between two experimental groups. In all cases, false discovery rate (FDR) correction was performed using Benjamini Hochberg correction^45^, and FDR-adjusted P value < 0.05 were considered significant. As a final step, customizable volcano plots were built to summarize the results from the above statistical tests.

The color codes for the volcano plots were inherited from a custom-built database providing the neuronal compartment, cell-type and known neuronal functions of individual lipid features to facilitate data interpretation (Table S3).

For ^2^H-labelled LC-MS samples, statistical comparison of ^2^H incorporation was performed between WT and HD mice using One-way ANOVA test followed by a Tukey HSD post-hoc testing and P value < 0.05 were considered significant.

#### Pathway enrichment analysis

Pathway enrichment analysis was performed through a Fisher exact test, using a custom-curated pathway database. An in-house pathway database was created by categorizing the detected lipid features based on their known biological functions/processes, cell type and cellular compartment, using previously published studies (For details, refer to Table S3). An FDR P value of 0.05 was used to assess significance.

## DATA AVAILABILITY

The paired *.ibd and *.imzML files of all kMSI datasets were deposited in Metaspace at https://metaspace2020.eu/api_auth/review?prj=bd3f06aa-36d8-11ec-96db-8319877174c6&token=eZWzn30yL6FP. Additionally, the raw data matrices (.csv files) and the *.ibd and *.imzML files have also been deposited in Figshare at https://figshare.com/s/a7a8940071e04e74c0b2.

## CODE AVAILABILITY

A comprehensive and detailed step-by-step guide for installing and using the KineticMSI package can be found on GitHub (https://github.com/MSeidelFed/KineticMSI). Alternatively, it can be directly installed into any R environment using devtools::install_github (‘MSeidelFed/KineticMSI’). Additionally, the guidelines to format the input tables for adapting kinetic LC/GC-MS data for usage with the KineticMSI R package can be found on GitHub (https://github.com/MSeidelFed/KineticMSI_2_kLCMS). An installation of R (Version R-3.6.2 or higher), Microsoft Windows operating systems and a CPU with at least 16GB RAM is recommended to run the workflow.

## ACKNOWLEDGEMENTS

We thank Dr Thibault Renoir for helping make the HD and WT mice available for tissue collection. We would also like to thank the Melbourne Mass Spectrometry and Proteomics Facility of The Bio21 Molecular Science and Biotechnology Institute at The University of Melbourne for the support of mass spectrometry analysis. This work was supported by National Health and Medical Research Council (NHMRC) through the Ideas Grant ID APP1184166 awarded to D Hatters, BA Boughton. D Hatters is an NHMRC Senior Research Fellow and AJ Hannan is an NHMRC Principal Research Fellow. F Farzana and F Martinez-Seidel contributed equally to this work and have the right to list their name first in their CV. All authors contributed to the article and approved the final submitted version.

## ETHICS DECLARATIONS

The authors declare no competing interests. All animal care and experimental procedures were approved by The Florey Institute of Neuroscience and Mental Health Animal Ethics Committee and were conducted by complying with the Australian Code of Practice for the Care and Use of Animals for Scientific Purposes as outlined by the National Health and Medical Research Council of Australia (Ethics number: 19-019-FINMH).

## ADDITIONAL INFORMATION

### Extended Data Figures

**Extended Data Fig. 1.**
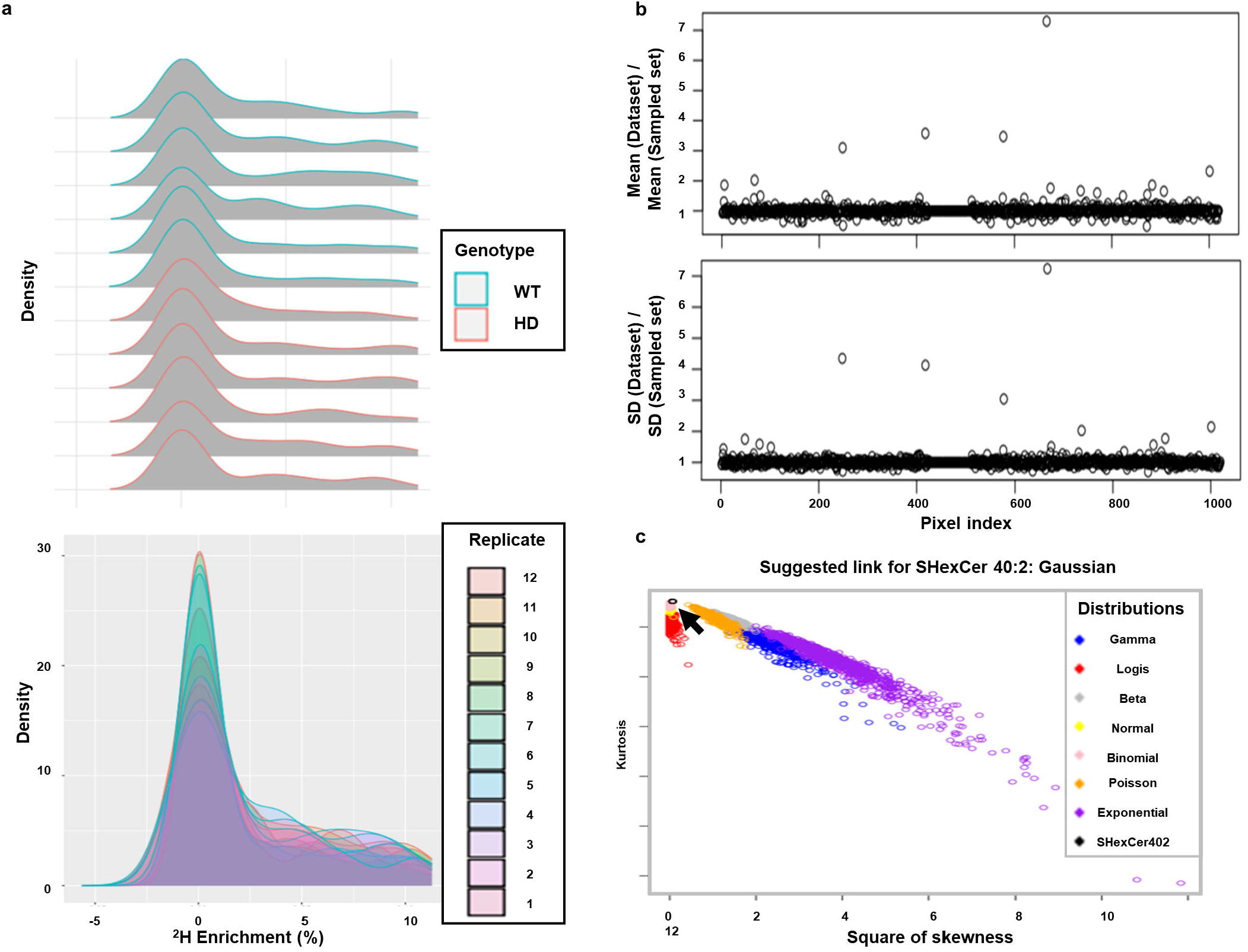
Visual evaluation of distribution of ^2^H incorporation in target lipids across MSI pixels of WT and HD kMSI datasets. **a**, Data distributions of ^2^H incorporation in SHexCer 40:2 (*m/z* 860.5) across MSI pixels of individual WT and HD replicate datasets (n=6/group). **b**, Evaluation of the correctness of random sampling of MSI pixels from WT and HD mice (index of all lipid features detected across the MSI pixels of WT and HD kMSI replicate datasets (X-axis) versus ratios of ^2^H incorporation mean and standard deviation from randomly sampled sub-subsets and the entire datasets (Y-axis). **c**, Evaluation of the distribution of ^2^H incorporation in SHexCer 40:2 using the R package RandodiStats. Based on the suggested link i.e., Gaussian distribution (shown by black arrow), a parametric test (GLM) was applied for mean comparison of ^2^H incorporation in SHexCer 40:2 between WT and HD mice.

**Extended Data Fig. 2.**
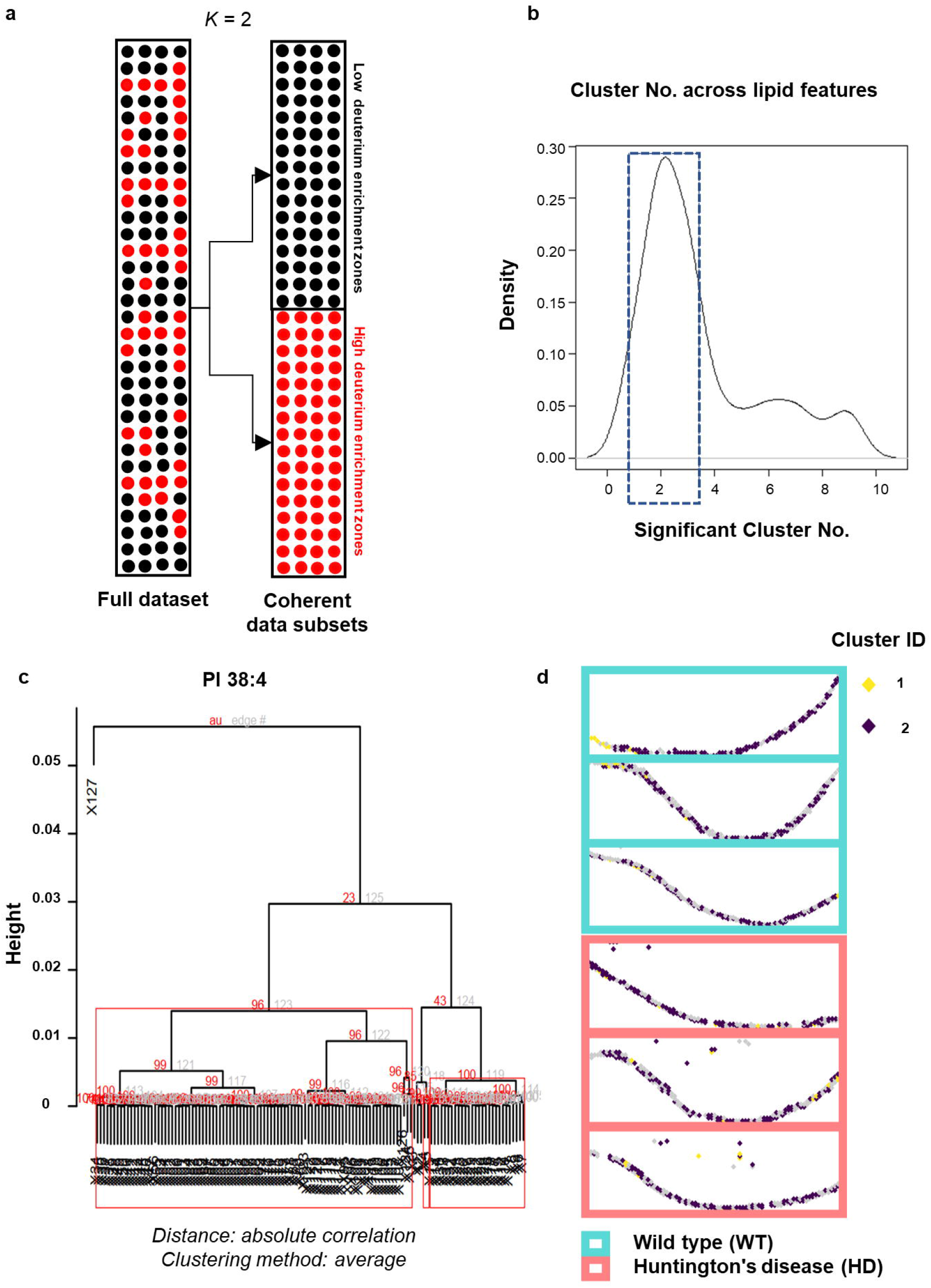
Sub-setting kMSI datasets into coherent pixel subsets based on ^2^H incorporation. **a**, Schematic diagram showing sorted matrices from all biological replicates (n = 6/group) (left) and subset matrices obtained through hierarchical clustering (k=2) corresponding to ‘low’ (black) and ‘high’ ^2^H (red) enrichment zones (right). **b**, Density histogram showing significant number of coherent partitions found across the evaluated features. Blue box highlights lipid features that returned two partitions based on ^2^H incorporation. **c**, Cluster dendrogram for PI 38:4 (*m/z* 885.5) showing optimum number of significant clusters (K=2) returned by bootstrapped hierarchical clustering algorithm. **d**, Reconstructed kMSI images displaying spatial distribution of significant cluster subsets obtained based on ^2^H incorporation from WT and HD mice. Clusters corresponding to ‘low’ and ‘high’ zones of ^2^H incorporation are highlighted in yellow (Cluster ID - 1) and purple (Cluster ID - 2). Grey pixels represent pixels that were excluded from analysis during random sampling process.

**Extended Data Fig. 3.**
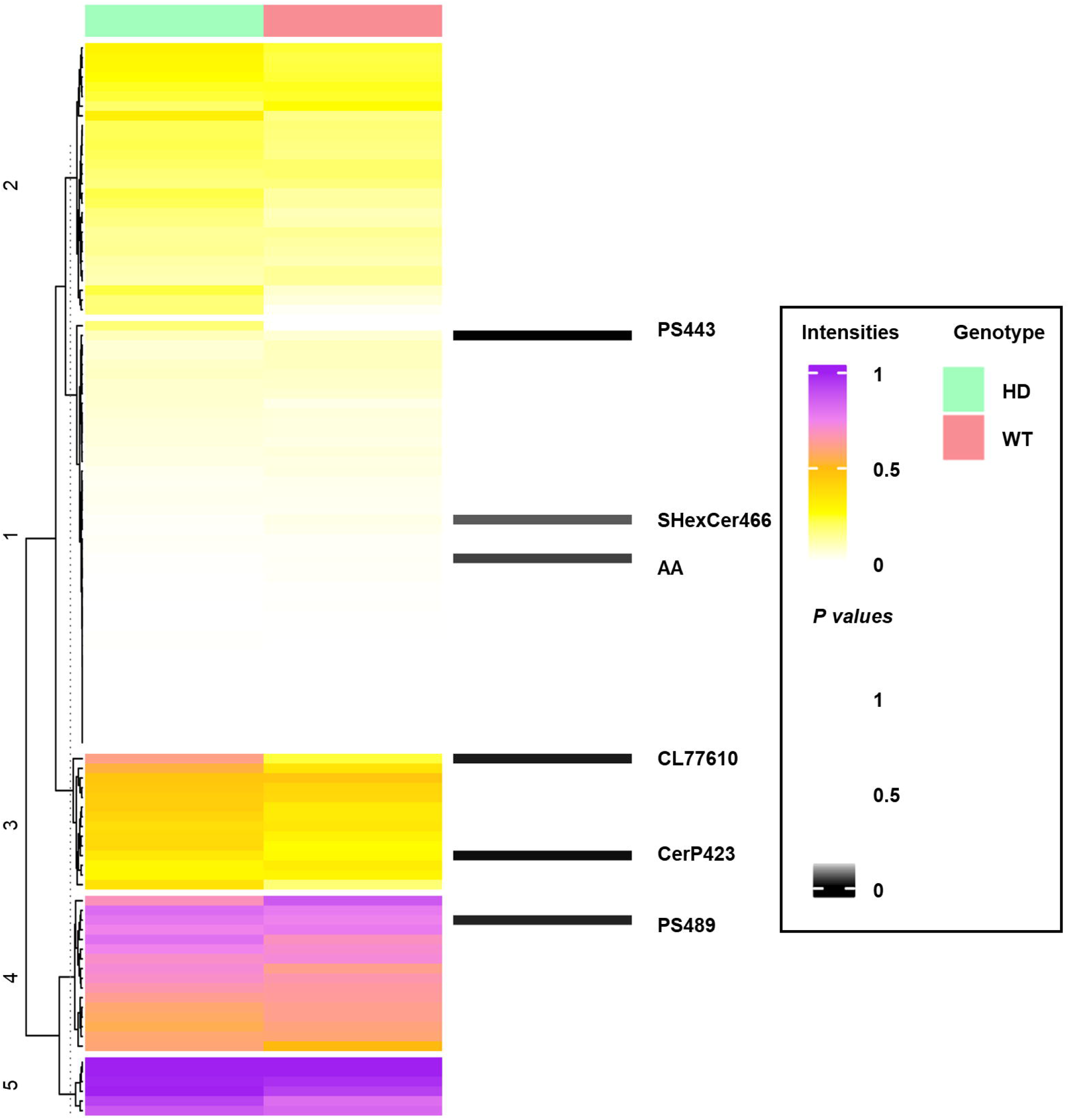
Heatmap representation showing the comparison of ^2^H enriched pixel proportions between WT and HD mice. The example shown here uses pixel proportions displaying high metabolic activity (> 15% ^2^H incorporation). K-means analysis was used to perform clustering of consolidated data matrices into clusters (k = 3) displaying similarity in ^2^H enriched pixel proportions. FDR adjusted P values obtained by performing class comparison of ^2^H enriched pixel proportions between WT and HD mice (n = 6/group) using GLM have been provided on the right. The significance level has been set to P < 0.05 (black bars). Only the significantly changed lipid features have been labelled in the heatmap.

**Extended Data Fig. 4.**
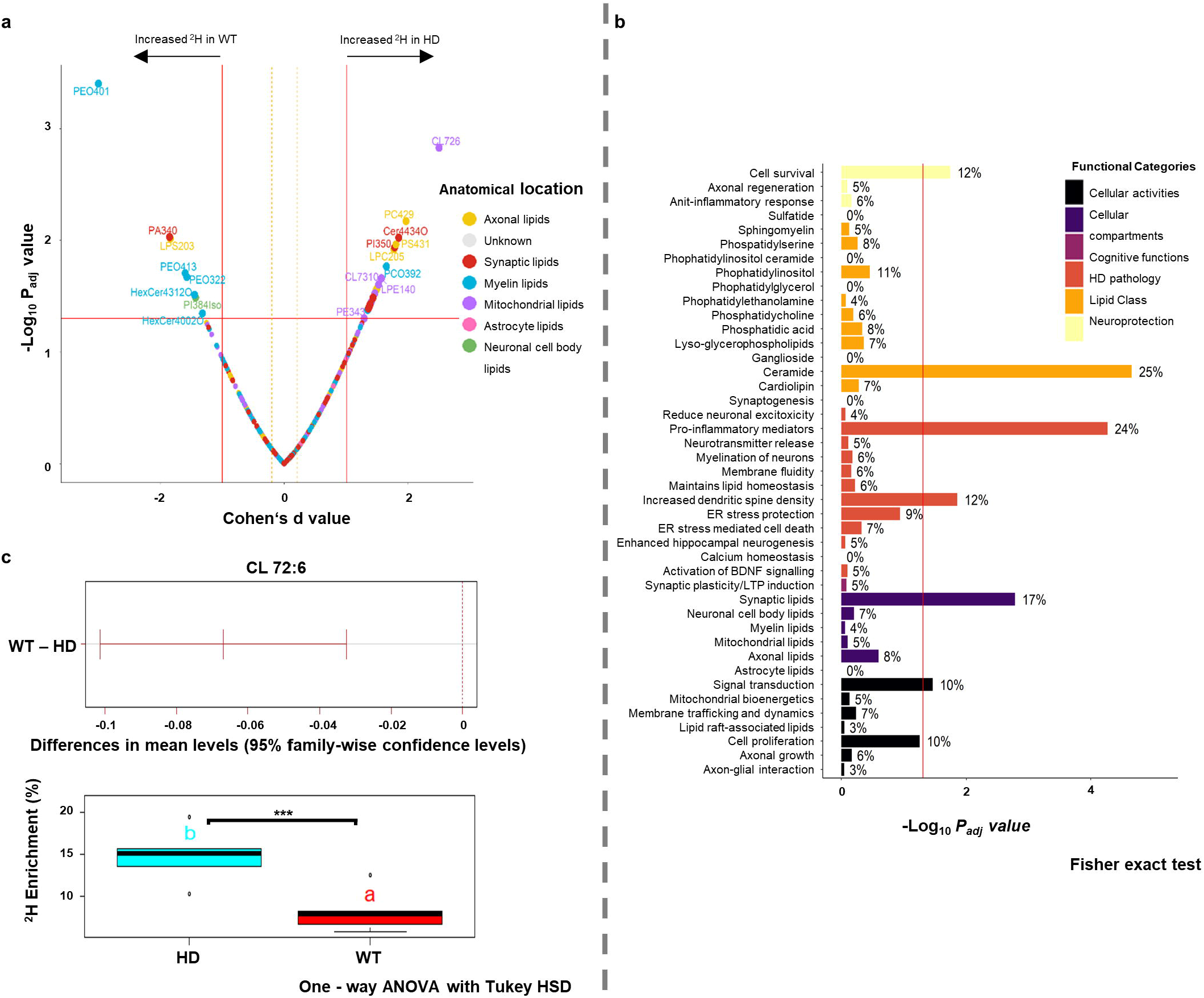
Differential analysis of ^2^H incorporation in lipids of WT and HD mice using homogenized hippocampal tissue by LC-MS. **a**, Volcano plot (Cohen’s d value (X-axis) versus -log_10_ FDR-adjusted P values obtained using one-way ANOVA followed by Tukey HSD post-hoc test (Y-axis)) showing significant differences in mean ^2^H incorporation in lipid features of WT and HD mice hippocampi (n = 6/group). Cohen’s d values ≥ 0.3 indicate greater differentiation of WT and HD mice. Each dot representing an individual lipid feature, is colored based on its known anatomical location in the mouse hippocampus. Grey dots represent features that have no defined category in the custom-built library provided for pathway enrichment analysis (Table S3). Significantly changed features are highlighted and labelled. **b**, Mean comparison of ^2^H incorporation in CL 72:6 (*m/z* 1451.9) showing significantly higher ^2^H incorporation in HD mice, compared to WT controls (n=6/group). **c**, Bar plot representation showing the results of pathway enrichment analysis (-log_10_ FDR-adjusted P value from the Fisher exact test (X-axis) versus functional categories associated with the detected lipid features in our study (Y-axis)). Functional categories are colored by lipid class, cellular activity, compartment, cognitive function, known HD pathology and neuroprotection to enhance interpretation of results (BDNF – Brain-derived neurotrophic factor; LTP – Long-term potentiation). Values denotes the proportion of lipids with significantly altered ^2^H incorporation in each functional category. Red line denotes significance P < 0.05.

