## Supplementary information for "KineticMSI, an R-based framework for relative quantification of spatial isotopic incorporation in mass spectrometry imaging experiments"

**Author Information**

Farheen Farzana^1,2^*^#^, Federico Martinez-Seidel^3,4^*^#^, Anthony J. Hannan^1^, Danny Hatters^2^, Berin A Boughton^4,5*^.

^1^ Florey Institute of Neuroscience & Mental Health, University of Melbourne, Melbourne, Australia; ^2^ Department of Biochemistry and Pharmacology, Bio21 Molecular Science and Biotechnology Institute, The University of Melbourne, Victoria 3010 Australia; ^3^ Molecular Physiology Department, Max Planck Institute of Molecular Plant Physiology, Potsdam, Germany; ^4^ School of Biosciences, University of Melbourne, Melbourne, Australia; ^5^Australian National Phenome Centre, Murdoch University, Perth, Australia.

^#^These authors contributed equally.

^*^For correspondence:

SUPPLEMENTARY INFORMATION

### **SUPPLEMENTARY RESULTS**

#### 1. **Calculation of deuterium incorporation**

Stable non-radioactive isotopic tracers (^2^H, ^13^C, ^15^N, ^18^O and ^34^S) serve as indispensable tools for determining *in vivo* kinetics of metabolism by monitoring the metabolism of a wide range of substrates such as DNA^2^, sugars^3^, proteins^4^ and lipids^5^. The choice of the tracer molecule depends on the pathway being investigated and has been extensively reviewed elsewhere^6^. Regardless of the analytical platform used for isotopic measurement, the general principle for calculating isotopic incorporation into metabolites remains the same. In principle, when a stable-isotope label (SIL) is incorporated into a biological system, the heavy isotope is substituted for light isotopes during metabolic synthesis and is observable by a specific mass shift when using mass spectrometric techniques. Using deuterated water (^2^H_2_O) as an example, the deuterium (^2^H) from the ^2^H_2_O replaces light ^1^H in metabolic reactions with a prime example being the growing hydrocarbon chain during fatty acid chain synthesis and elongation, thus providing an indicator of phospholipid synthesis and membrane activity in the tissue being interrogated. This slow process of labelling becomes visible as a shift in the molecular weight of the metabolic species for every ^1^H-^2^H substitution, thus modulating the isotope distribution pattern detected by MSI as shown in Fig. S2. For instance, for the substitution of a single ^1^H atom by ^2^H, we would expect a shift in the isotopologue by 1 Dalton as ^2^H is approximately one atomic mass unit heavier than its most abundant counterpart (i.e., ^1^H) (Fig. S2a and b). A complete ^2^H incorporation in a fully labelled state results in the complete disappearance of M_0_/A_0_ accompanied by an increase in the signal of its labelled isotopic envelope (Fig. S2b). However, SIL experiments usually yield partially labelled molecules (Fig. S2c), which is apparent by a decrease in the intensity of the monoisotopic peak (M_0_/A_0_) and an equivalent increase in the intensity of the isotopologues (M_1_, M_2_…. M_n_). Note that Fig. S2 shows the case where there is only a single site of substitution, however in the case of individual fatty acids the possible distribution is more complex because there are H_n_ possible substitutions.

#### 2. **Determination of the most suitable isotope ^2^H incorporation proxy**

To determine the most accurate proxy that matched the biology of the actual pool changes, we compared batch effect corrected steady-state lipid pools derived from unlabelled control animals (Fig. S3a) to equivalent labelled lipid species from the labelled mice (Fig. S3). We calculated two ratios between the ^2^H-labelled and steady state lipid pools to test if a single-labelled monoisotopic ratio (M_1_/M_0_) or the ratio of summed intensities of all the labelled isotopes ((∑ M_1_ + M_2_ + ...M_n_) / M_0_) represents the most suitable proxy for measuring ^2^H incorporation in the examined lipid features. These ratios include: 1. Ratio between the summed intensities of the M_0_ and M_1_ peaks from the labelled pools and the intensity of monoisotopic peak (M_0_) from unlabelled mice and, 2. Ratio between summed intensities of M_0_ and all the detected labelled isotopologues (M_1_, M_2_ …M_n_, where n is the last detected labelled isotopologue) from ^2^H-labelled mice and the intensity of monoisotopic peak (M_0_) from unlabelled mice. The calculation of these ratios allowed us to test two contrasting theoretical scenarios. In the first case, ^2^H would be expected to be incorporated exclusively into the first isotopic peak M_1_, and so the single-labelled monoisotopic ratio (M_1_/M_0_) can be considered the legitimate proxy for measuring ^2^H incorporation. On the other hand, if ^2^H was incorporated uniformly across all the detected isotopologue peaks (M_1_, M_2_ …M_n_), the summed intensities of all the labelled isotopes ((∑ M_1_ + M_2_ + ...M_n_) / M_0_) would be considered the most legitimate proxy method.

An ideal scenario would be to achieve a ratio of one between labelled and steady state pools for all examined lipid features, which would indicate that the biology in terms of lipid pool sizes between the labelled and unlabelled specimens is fully conserved across all treatments. However, to account for biological variations, we accepted a threshold between 0.7 - 1.4. Therefore, the aim of this step is to consider the heatmap that satisfies the criterion of the highest number of lipid features with a ratio close to one and select the associated proxy as the best measure of ^2^H incorporation.

As can be readily seen in the heatmap of the labelled to unlabelled steady state pool ratios (Fig. S3b) for some lipids the biology is completely preserved, with the ratio close to one (represented by grey). While for others, the abundances in the steady state pools are either higher than the labelled pools (ratio < 0.7 - white) or lower than the labelled pools (ratio > 1.4 - black). It is worth noting that the heatmap on the right, created by summing all isotopologues (M_0_ to M_n_) contained more lipid features with a ratio close to one, compared to the heatmap on the left that was based only on the sum of M_0_ and M_1_ labelled isotopes. This suggests that in our example dataset, the ^2^H incorporation calculations are closer to reality when all isotopic peaks (M1, M2…Mn) are considered, and thus provide a better proxy to measuring metabolite synthesis and were therefore selected for subsequent statistical analysis.

Upon further examination of the lipid features whose ratios deviated from a predefined range of 0.7-1.4, we found that peaks with a ratio smaller than 0.7 (white colors) correspond to metabolites present in very low amounts in the unlabelled pools. It is therefore likely that the corresponding labelled pools, particularly the heavier isotopologues (M_4_, M­_5_), which are much less abundant in nature, were not detected due to the lack of sensitivity of the instrument, explaining the observed decrease in the ratio of labelled to unlabelled pools for these lipids. On the other hand, lipids with ratios greater than 1.4 (black colors) indicate an "extra" abundance of labelled isotopologues due to the presence of interfering peaks, which is likely an artifact of peak coalescence, i.e., the merging of two closely spaced peaks with similar mass-to-charge ratios into a single average peak. This overlap between labelled isotope peaks of one metabolite and the monoisotopic peak of the adjacent metabolite resulted from the lack of sufficient resolving power at the higher molecular weight end of the labelled spectra in our study, adding “extra” biased abundances to the labelled isotopologues and hence a higher resolving power in kMSI studies can help to minimize peak coalescence in complex SIL spectra. Nevertheless, while continuing our analyses, we interpreted isotope ^2^H results derived from metabolites with labelled to steady-state pool ratios outside the range of 0.7 - 1.4 with caution.

#### 3. **Quality assessment of consolidated data matrices**

To evaluate the correctness of random sampling and ensure that the procedure preserved the mean and variance structures of the complete datasets, we confirmed that the mean and standard deviation ratios of ^2^H incorporation from randomly sampled sub-subsets and the entire datasets was close to 1 across the examined lipid features (Extended data fig. 1b). Finally, a consolidated matrix containing a common number of pixels from each kMSI replicate of WT and HD was built and the mean ^2^H incorporation in each lipid species across the selected pixels was computed. Next, taking advantage of the high number of spatial data points per lipid feature, we assessed the data distributions of ^2^H incorporation across MSI pixels and determined the appropriateness of using parametric or non-parametric class comparison statistical tests using the procedure detailed in the R package RandoDiStats (<https://github.com/MSeidelFed/RandodiStats_package>) (Extended data fig. 1a and c). For instance, based on the suggested link i.e., Gaussian distribution for the exemplary lipid SHexCer 40:2 (*m/z* 860.5) (Extended Fig. 1d), a parametric test (GLM) was used for mean comparison of ^2^H incorporation between WT and HD, whereas for other lipids that were not normally distributed, a parametrized GLM was used.

### **EXPERIMENTAL METHODOLOGY**

#### **Animal care**

R6/1 mouse model of Huntington’s disease (HD) and wild-type (WT) littermates were generated by crossing F1 hybrid (CBA x C57Bl/6) females and hemizygous R6/1 males. Animals were bred on site and randomly group housed (4-5 mice/cage) by genotype in a fully controlled SPF facility (12 h light/dark cycle, temperature (22 ºC) and humidity (45%)) at the Florey Institute of Neuroscience and Mental Health. All mice were provided *ad libitum* access to food and water. HD mice and age-matched WT controls (n = 6/group) at 16 weeks of age were randomly assigned to the non-labelled and deuterium (^2^H) labelled groups. The ^2^H labelled groups were injected with deuterated water (99% ^2^H_2_O and 0.9% NaCl) via an intraperitoneal injection bolus of 35 µl/gm, followed by a maintenance dose of 9% deuterated water in drinking water. For the sake of comparison, unlabelled control mice were maintained on regular drinking water, without access to deuterium water. The investigators were blinded to experimental group allocation during sample preparation. Post-sectioning, we randomly selected WT-HD (non-labelled and labelled) pairs to collect the MSI data across multiple batches (4 samples/batch).

#### **Tissue collection**

Mice were euthanized 8 days post-labelling using cervical dislocation, and the excised brain tissues were hemi-sectioned on a cold plate. While one brain hemisphere (left) was flash-frozen on liquid nitrogen and reserved for mass spectrometry imaging (MSI), the other matched hemisphere (right) was dissected to obtain the frontal cortex, hippocampus, and striatum and examined in detail using LC-MS/MS analysis. The brain regions were immediately snap-frozen in liquid nitrogen (within 3-5 minutes after brain excision) and stored at -80ºC until further use. The left hemisphere of the brain was mounted onto the chuck using Optimal Cutting Temperature (OCT) embedding medium (Thermo Fisher Scientific, Waltham, MA, USA) and cryosectioned using Leica 1950 cryostat at -17ºC, ensuring sections were not contaminated with OCT. Serial coronal sections from 1.42 to -2.48 mm Bregma were obtained at 20 μm thickness and collected in a one in 10 series spaced 200 μm apart. Sections were thaw mounted on microscopic glass slides (Menzel -Glaeser Superfrost Ultra Plus, Thermo Fisher Scientific, Waltham, MA, USA). While one section was dehydrated in a vacuum desiccator for 15 minutes and used for MSI measurement, the neighbouring section was stained with hematoxylin and eosin (H&E) to check the integrity of the tissue and obtain anatomical information. High-resolution histological images of the H&E-stained sections were recorded using a Pannoramic MIDI digital slide scanner (3D Histech, Budapest, Hungary).

#### **Sample preparation for MALDI-MSI**

Norharmane matrix was prepared by dissolving norharmane in 95% acetone in Milli-Q water (5 mg/ml), followed by the addition of sodium anthraquinone-2-sulfonate (500 ng/ml in 1:1 norharmane solution:95% acetone) as an internal standard. An automatic TM-Sprayer matrix application device (HTX Technologies LLC) attached to LC20-AD HPLC pump (Shimadzu Scientific Instruments) was used for matrix deposition by applying the following method parameters: flow rate - 0.15 ml/min, a gas flow rate of 10 litres/min, nitrogen pressure - 10 psi and a nozzle temperature of 30°C. The spray conditions used were as follow: 16 passes, nozel velocity - 1200 mm/min, track spacing – 2 mm, with alternate passes set at 90º offset, repeat passes set to 1mm and drying time as 0 seconds. The slides were assessed for an even application of matrix crystals using light microscopy prior to data acquisition.

#### **Data acquisition using MALDI-MSI**

MALDI–MSI experiments were performed using a Bruker SolariX 7T XR hybrid ESI–MALDI–FT–ICR–MS platform equipped with a SmartBeam II UV laser. Intact lipid spectra were acquired over *m/z* 100 – 2000, across 30 × 30 µm pixel/raster array in the negative ionization mode using age- and gender-matched WT and HD pairs. The laser power was set to 25%, and 300 shots per spot were collected at a frequency of 2 kHz. The laser was set to the minimum focus with smart walk enabled within a 25 µm area, a grid increment of 10 and offset of 1 providing an ablation spot of ~30 µm. Image acquisition was performed with a Time Domain for Acquisition of 2 megaword, providing an estimated resolving power of 130,000 at 400 m/z. With every batch-run, on-line calibration was performed using a single reference mass peak corresponding to the internal standard (287.00197 *m/z* ± 0.05 Da). Bruker Daltonics ftmsControl 2.1.0 and FlexImaging 5.1 was used to acquire the mass spectra.

#### **MALDI-MSI data processing**

Following data acquisition, the overall average spectra was visualized using SCiLS Lab software (Version 7.02.10980 - 2019c Bremen, Germany) across *m/z* 250 – 2000 Da. Data pre-processing steps including data reduction using binning method (bin width = 0.01 Da.) and peak alignment was performed using SCiLS Lab. To visualize spatial distribution of lipids across the tissue sections, MS images were total ion count (TIC) normalized and for quantitative comparison of lipid abundances and synthesis rates between WT and HD mice, the signal intensities of all the lipid features were normalized to a single internal standard peak (*m/z* 287.00197 ± 0.05 Da). Lipid assignment was performed by accurate precursor mass (<5ppm) search using online databases such as LIPID MAPS^8^, Human metabolome database^9^ and Swiss Lipids^10^ (Table S1). The presence of the species level lipids from MS1 only MSI data were verified using identified lipids at the molecular species level from LC-MS/MS (i.e., MS2) data from the unlabelled lipid extracts of homogenized hippocampal regions of WT and HD mice. For MSI lipid annotations, names were assigned to the species level using the following convention: lipid class, number of carbons contained in side chains and number of double bonds as per LipidMaps recommendations^13^*.*

*^13^*. Standard use of punctuation found in normal naming convention has been removed as special characters are not allowed in variable names in R.

Peak picking and integration were visually confirmed to screen for peak coalescence i.e., the merging of two closely spaced peaks with similar mass-to-charge ratios into a single average peak. Metabolites whose labelled isotopes (M_1,_ M_2_…M_n_) overlapped with the monoisotopic peak of the adjacent metabolite were filtered out as they would otherwise lead to inaccurate ^2^H incorporation calculations. Manual peak selection, while time-consuming, provided several advantages for downstream multivariate statistical analysis including removal of background noise and confirmation of true signal peaks for complex stable isotope labelled (SIL) spectra.

#### **Multivariate analysis of MALDI-MSI data**

Spatial segmentation was performed by applying Probabilistic latent semantic analysis with deterministic initialization (PLS-DA) in SCiLS Lab to segment data from mouse hippocampus into anatomically distinct layers based on lipid compositions in an unsupervised manner. The component that best matched the spatial expression of hippocampus proper was selected and further segmented using a k-mean clustering algorithm with edge-preserving image denoising in SCiLS Lab. The resulting k-mean clusters were assigned anatomical labels corresponding to the CA1, CA3 and DG sub-fields by visually mapping to the most closely aligned anatomical structures in the Allen Mouse Brain Atlas^1^, as confirmed by an expert in Anatomy. For validation of KineticMSI workflow, the CA1 hippocampal sub-field was selected as a region of interest (ROI) for comparison of metabolic activity between WT and HD mice.

#### **Monophasic lipid extraction from mouse brain hippocampus for LC-MS analysis**

All samples were prepared for homogenization by combining the total hippocampal tissue with ice-cold 60% methanol in MilliQ water containing 0.01% (v/v) butylated hydroxytoluene (BHT) (200µl), Mouse splash lipidomix (10 µl), (Avanti Polar Lipids, Albaster, AL, USA) and deuterated lipid standards : C18:0 (d5) GM1 Ganglioside (10 µM, 10 µl), 14:0 Cardiolipin (30 µM, 20 µl,), Ceramide-d7 (30 µM, 5 µl) (Avanti Polar Lipids, Albaster, AL, USA) and d18:1/18:0 (d3) C18 3-sulfo Galactosylceramide (15 µM, 5 µl) (Cayman Chemical, Ann Arbor, MI, USA). Homogenization was performed for 30 seconds 6 times at 15 s intervals using a Bullet blender (Next Advance, Averill Park, NY) and 0.5 mm Zirconium Oxide beads (100 µl), followed by sonication in a water bath sonicator (20 min). Samples were then subjected to monophasic lipid extraction method as described by Lydic et al.^11^ with modifications as stated below. Homogenates were combined with MilliQ water (120 µl), methanol with 0.01% (w/v) BHT (420 µl) and chloroform with 0.01% (w/v) BHT (260 µl) (the ratio of total H_2_O: CHCl_3_: MeOH used was 0.74:1:2). Samples were vortexed thoroughly and shaken (Eppendorf Thermomixer Compact 5350 Mixer) at 1,400 rpm at room temperature (30 min). Extracts were then centrifuged (Beckman Coulter Microfuge 20R) at 20,000 g at room temperature (15 min) and the supernatants collected. The remaining pellets were resuspended in MilliQ water (100 µl), vortexed and re-extracted by adding chloroform: methanol (1:2, v/v) containing 0.01% (w/v) BHT (420 µl), followed by incubation and centrifugation as described above. The monophasic supernatants from repeat extractions were pooled, dried by evaporation under vacuum using a GeneVac miVac sample concentrator (SP Scientific Warminster, PA, USA) and resuspended in chloroform: methanol (1:9, v/v) containing 0.01% (w/v) BHT (50 µl), vortexed and incubated with 1,400 rpm shaking at room temperature (3 min) using a Thermomixer (Eppendorf Thermomixer Compact 5350 Mixer). The reconstituted brain lipid extracts were pelleted by centrifugation 20,000 g at room temperature (15 min). The supernatants were then transferred to 2.0 ml glass vials and stored at -20°C until further use.

#### **Data acquisition by LC-MS/MS**

Samples were analyzed using a Vanquish ultra-high-performance liquid chromatography (UHPLC) coupled to an Orbitrap Fusion Lumos MS (Thermo Fisher Scientific), with separate injections for positive and negative ion polarity in a randomized manner. Solvent A was 6/4 (v/v) acetonitrile/water with 10 mM ammonium acetate and 5 µM medronic acid and solvent B was 9/1 (v/v) isopropanol/acetonitrile with 10 mM ammonium acetate. Each sample was injected into an RRHD Eclipse Plus C18 column (2.1 × 100 mm, 1.8 µm, Agilent Technologies) at 50°C at a flow rate of 350 μL/min for 3 min using 3% solvent B. During separation, the percentage of solvent B was increased from 3% to 70% in 5 min, from 70% to 99% in 16 min, from 99% to 99% in 3 min, from 99% to 3% in 0.1 min and maintained at 3% for 3.9 min.

LC MS/MS analysis was performed using a Heated Electrospray Ionization (HESI) source, with spray voltages of 3.5 kV for positive and 3.0 kV in negative ionization-mode respectively. Gas flow rate for sheath, auxiliary and sweep gases were 20 and 6 and 1 ‘arbitrary’ unit(s), respectively. The ion transfer tube and vaporizer temperatures were maintained at 350 °C and 400 °C, respectively, and the S-Lens RF level was set at 50%. In both positive and negative ionisation-mode from 3 to 24 min, top speed data-dependent scan with a cycle time of 1 s was used. Within each cycle, a full-scan MS spectrum was first acquired in the Orbitrap at a mass resolving power of 120,000 (at *m/z* 200) across *m/z* 300–2000 using quadrupole isolation, an automatic gain control (AGC) target of 4e5 and a maximum injection time of 50 milliseconds, followed by higher-energy collisional dissociation (HCD)-MS/MS at a mass resolving power of 15,000 (at *m/z* 200), a normalized collision energy (NCE) of 27% at positive mode and 30% at negative mode, an *m/z* isolation window of 1, a maximum injection time of 22 milliseconds and an AGC target of 5e4. For the improved structural characterization of glycerophosphocholine (PC) in positive mode, a data-dependent product ion (*m/z* 184.0733)-triggered linear ion trap collision-induced dissociation (CID)-MS/MS scan was performed in the cycle using a q-value of 0.25 and a NCE of 30%, with other settings being the same as that for HCD-MS/MS. For the improved structural characterization of triacylglycerol (TG) lipid in positive mode, the fatty acid + NH_3_ neutral loss product ions (Table S2) observed by HCD-MS/MS were used to trigger the acquisition of the top-3 data-dependent linear ion trap CID-MS^3^ scans in the cycle using a q-value of 0.25 and a NCE of 30%, with other settings being the same as that for HCD-MS/MS.

#### **LC data pre-processing and analysis**

LC-MS/MS data was processed using MS Dial 4.70^12^. The following parameters were used for peak selection and lipid annotation: mass accuracy 0.005 Da and 0.025 Da for MS1 and MS2, a minimum peak height of 50,000 and mass slice width of 0.05 Da. The identification score cut off is 80%. In positive mode, [M+H] ^+^, [M+NH_4_] ^+^ and [M+H-H2O] ^+^ adducts were selected. In negative mode, [M-H]^-^ and [M+CH3COO]^-^ were selected. All lipid classes available were searched. The retention time tolerance for alignment is 0.1 min. Lipids with maximum intensity less than 5-fold of average intensity in blank were removed. All other settings were default. All lipid LC-MS/MS features were manually inspected and re-integrated when needed. These four types of lipids, 1) lipids with only sum composition except SM, 2) lipid identification due to peak tailing, 3) retention time outlier within each lipid class, 4) LPA and PA artifacts generated by in-source fragmentation of LPS and PS were also removed. Shorthand notation for lipid classification and structural representation was used according to LipidMaps recommendations^13^.

The peak areas of the identified lipid ions were exported from MS Dial 4.70^12^ and normalized to the tissue weight of individual samples. This was followed by natural isotope abundance correction and calculation of ^2^H incorporation using the R package - IsoCorrectoR^14^. For the subsequent statistical analysis, the input tables were formatted using the provided guidelines for adapting kinetic LC/GC-MS data for usage with the KineticMSI R package (https://github.com/MSeidelFed/KineticMSI_2_kLCMS). The data analysis was fully automated so that subjective factors would not impinge on the quantification of deuterium incorporation.

### **FIGURE LEGENDS**

**Fig. S1: Selection of region of interest for validation of KineticMSI package.** **a**, H&E-stained image of mouse hippocampus showing the CA1 pyramidal sub-field within the mouse hippocampus that was the focus of our study. The nuclei (blue) within the CA1 pyramidal neuronal cell bodies and the cytoplasm (pink) are shown in the inset (black arrows). Scale bars, 100µm; inset - 20µm. **b**, k-mean clustering of the mouse hippocampal formation MSI dataset using SCiLS Lab software showing the top three k-mean clusters (CA1/CA3 and DG sub-fields). Scale bars, 1mm. **c**, Coronal view of hippocampal sub-fields from Allen Mouse Brain Atlas^1^.

**Fig. S2: Conceptual schematic diagrams of ^2^H incorporation calculation. a,** Unlabelled isotopologue showing the relative abundance of monoisotopic peak (M_0_) and naturally occurring isotope signals (M_1_ - M_3_). Shift in isotopologue patterns by ^2^H labelling experiments in **b,** Fully labelled state and **c,** Partially labelled state. This figure shows an exemplary case where there is only a single site of substitution and hence applies to a single addition of a stable-isotope molecule.

**Fig. S3: Data preparation and pre-processing steps.** **a**, Batch effect correction of MSI datasets acquired from non-labelled control animals (WT and HD mice - n = 6/group) at the steady state. **b**, Comparison of non-labelled and labelled steady state pools to determine the most legitimate proxy for measuring isotopic incorporation that best reflects the changes in steady state lipid pools. The grey bars represent lipid species with a ratio between equivalent labelled and non-labelled steady states ranging between 0.7 and 1. The black and white bars indicate a ratio > 1.4 and < 0.7 respectively.

**Fig. S4: Differential analysis of ^2^H incorporation in brain lipids of WT and HD mice using pixel cluster means.** **a**, Volcano plot (Cohen’s d value (X-axis) versus -log_10_ FDR-adjusted P values obtained using GLM (Y-axis)) showing significant differences in mean ^2^H incorporation of pixel clusters in examined lipid features of WT and HD mice hippocampi (n = 6/group). Cohen’s d values ≥ 0.3 indicate greater differentiation of WT and HD mice. Each dot represents an individual cluster (pixel subset sharing similar ^2^H incorporation) from the examined lipid feature (Cluster IDs removed for clarity) and is coloured by its known anatomical location in the mouse hippocampus. Grey dots represent features that have no defined category in the custom-built library provided for pathway enrichment analysis (Table S3). Significantly changed features are highlighted and labelled. **b**, Bar plot representation showing the results of pathway enrichment analysis (-log_10_ FDR-adjusted P value from the Fisher exact test (X-axis) versus functional categories associated with the detected lipid features in our study (Y-axis)). Functional categories are colored by lipid class, cellular activity, compartment, cognitive function, known HD pathology and neuroprotection to enhance interpretation of results (BDNF – Brain-derived neurotrophic factor; LTP – Long-term potentiation). Values denotes the proportion of lipids with significantly altered ^2^H incorporation in each functional category. Red line denotes significance P < 0.05.

**Fig. S5: Spatial segmentation of WT and HD kMSI datasets using Cardinal versus KineticMSI.** **a**, Spatially aware shrunken centroid clustering performed on CA1 hippocampal sub-field of a representative WT-HD replicate pair using Cardinal (k=3, r=6, s=4). **b**, Spatial segmentation of ^2^H incorporation in PI 38:4 (*m/z* 885.5) in the CA1 hippocampal sub-field of a representative WT-HD replicate pair using KineticMSI using bootstrapped Hierarchical cluster analysis (HCA) AU-P = 0.95), which is a dependency from the R package pvclust^7^.

**Table S1: List of tentatively assigned lipids from MALDI-MSI by accurate precursor mass <5ppm search.**

**Table S2: Inclusion list for improved structural characterization of triacylglycerol (TG) lipid.**

**Table S3: Lipids categorized based on known biological functions/processes, cell type and cellular compartment in brain tissue, based on previously published studies.**
