## Supplement Figure 1 for "KineticMSI, an R-based framework for relative quantification of spatial isotopic incorporation in mass spectrometry imaging experiments"

### Slide 1
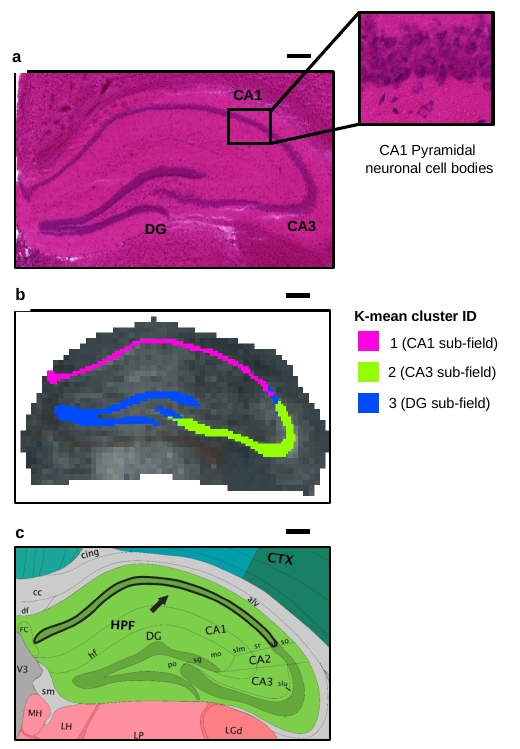

CA1
CA1 Pyramidal
neuronal cell bodies
CA3
DG
K-mean cluster ID
2 (CA3 sub-field)
3 (DG sub-field)
a
b
1 (CA1 sub-field)
c
