## Supplement Figure 2 for "KineticMSI, an R-based framework for relative quantification of spatial isotopic incorporation in mass spectrometry imaging experiments"

### Slide 1
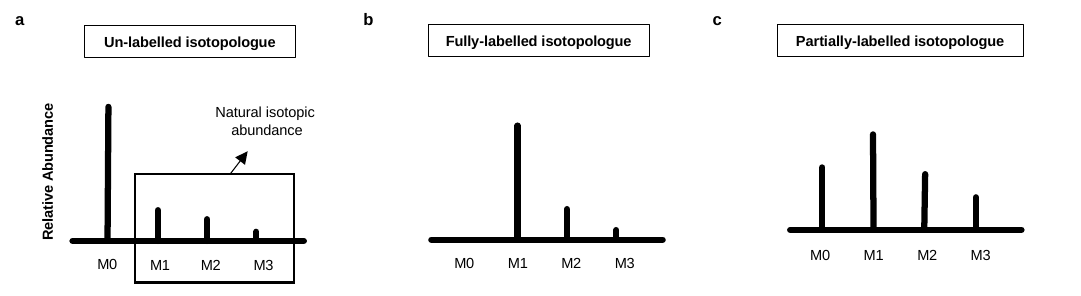

a
b
c
Fully-labelled isotopologue
Partially-labelled isotopologue
Un-labelled isotopologue
Natural isotopic
abundance
M0
M1
M2
M3
M0
M1
M2
M3
M0
M1
M2
M3
 Relative Abundance
