## Supplementary figures and images for "KineticMSI, an R-based framework for relative quantification of spatial isotopic incorporation in mass spectrometry imaging experiments"

### Supplement Figure 3

#### Slide 1
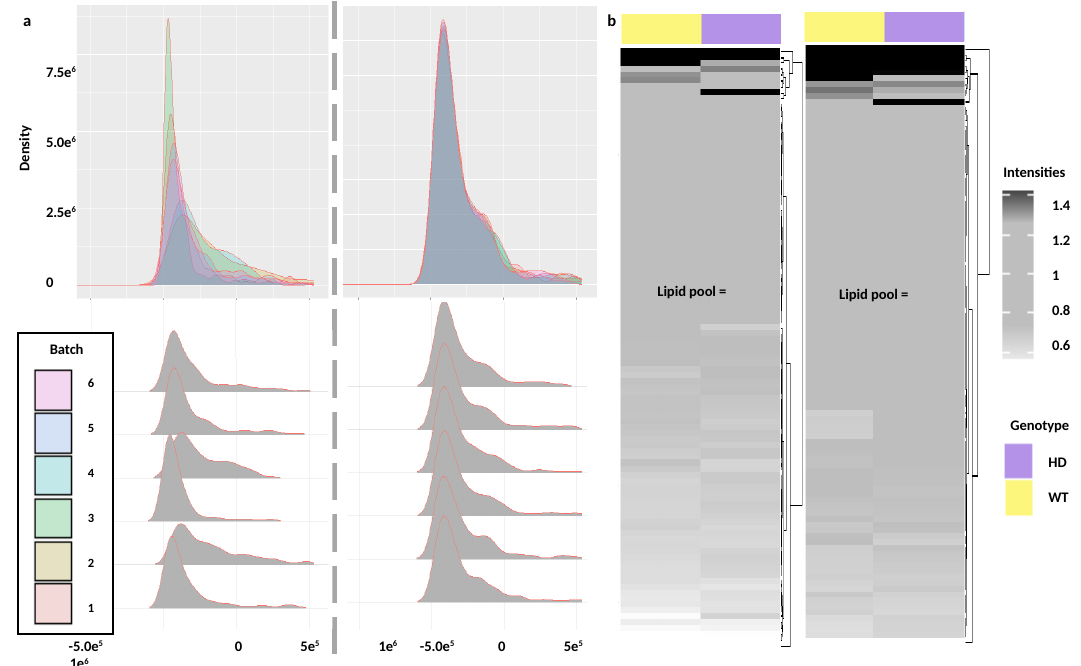

b
a
7.5e^6^
5.0e^6^
2.5e^6^
0
Density
Intensities
1.4
1.2
1
0.8
0.6
Batch
6
5
4
3
2
1
Genotype
HD
WT
 -5.0e^5^ 	 0 5e^5^	 1e^6^ -5.0e^5^ 0 5e^5^ 1e^6^
