## Supplement Figure 4 for "KineticMSI, an R-based framework for relative quantification of spatial isotopic incorporation in mass spectrometry imaging experiments"

### Slide 1
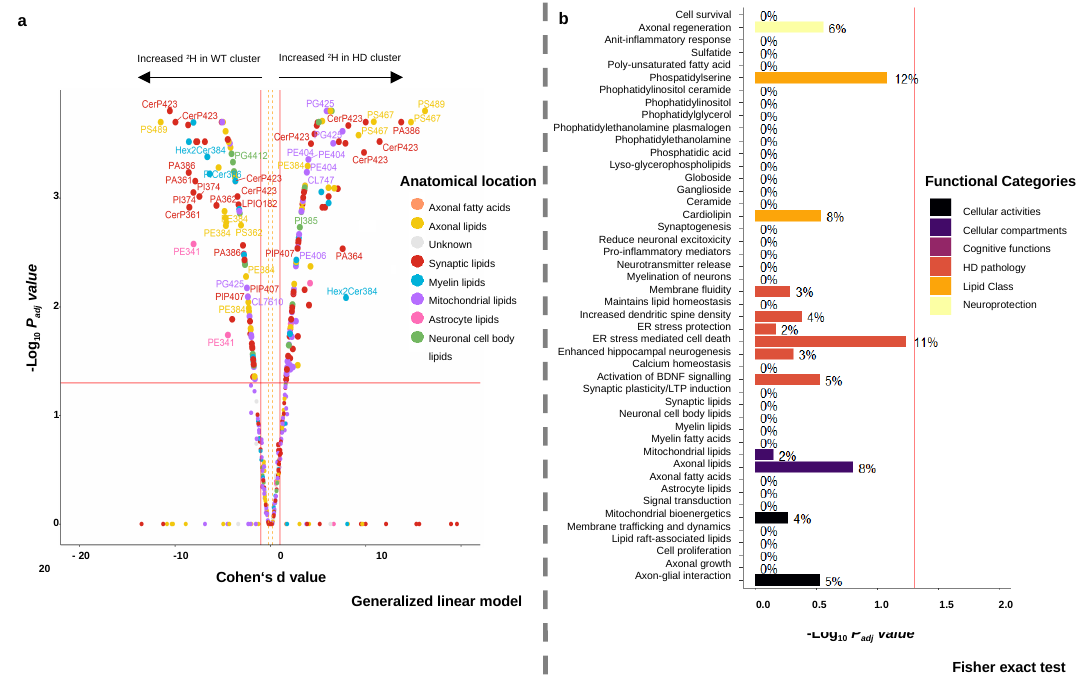

b
a
Cell survival
Axonal regeneration
Anit-inflammatory response
Sulfatide
Poly-unsaturated fatty acid
Phospatidylserine
Phophatidylinositol ceramide
Phophatidylinositol
Phophatidylglycerol
Phophatidylethanolamine plasmalogen
Phophatidylethanolamine
Phosphatidic acid
Lyso-glycerophospholipids
Globoside
Ganglioside
Ceramide
Cardiolipin
Synaptogenesis
Reduce neuronal excitoxicity
Pro-inflammatory mediators
Neurotransmitter release
Myelination of neurons
Membrane fluidity
Maintains lipid homeostasis
Increased dendritic spine density
ER stress protection
ER stress mediated cell death
Enhanced hippocampal neurogenesis
Calcium homeostasis
Activation of BDNF signalling
Synaptic plasticity/LTP induction
Synaptic lipids
Neuronal cell body lipids
Myelin lipids
Myelin fatty acids
Mitochondrial lipids
Axonal lipids
Axonal fatty acids
Astrocyte lipids
Signal transduction
Mitochondrial bioenergetics
Membrane trafficking and dynamics
Lipid raft-associated lipids
Cell proliferation
Axonal growth
Axon-glial interaction
Increased ^2^H in HD cluster
Increased ^2^H in WT cluster
Anatomical location
Axonal fatty acids
Axonal lipids
Unknown
Synaptic lipids
Myelin lipids
Mitochondrial lipids
Astrocyte lipids
Neuronal cell body lipids
Functional Categories
Cellular activities
Cellular compartments
Cognitive functions
HD pathology
Lipid Class
Neuroprotection
3
2
-Log_10_ P_adj_ value
1
0
 - 20 -10 0 10 20
Cohen‘s d value
Generalized linear model
 0.0 0.5 1.0 1.5 2.0
-Log_10_ P_adj_ value
Fisher exact test
