## Supplementary Figure 5 for "KineticMSI, an R-based framework for relative quantification of spatial isotopic incorporation in mass spectrometry imaging experiments"

### Slide 1
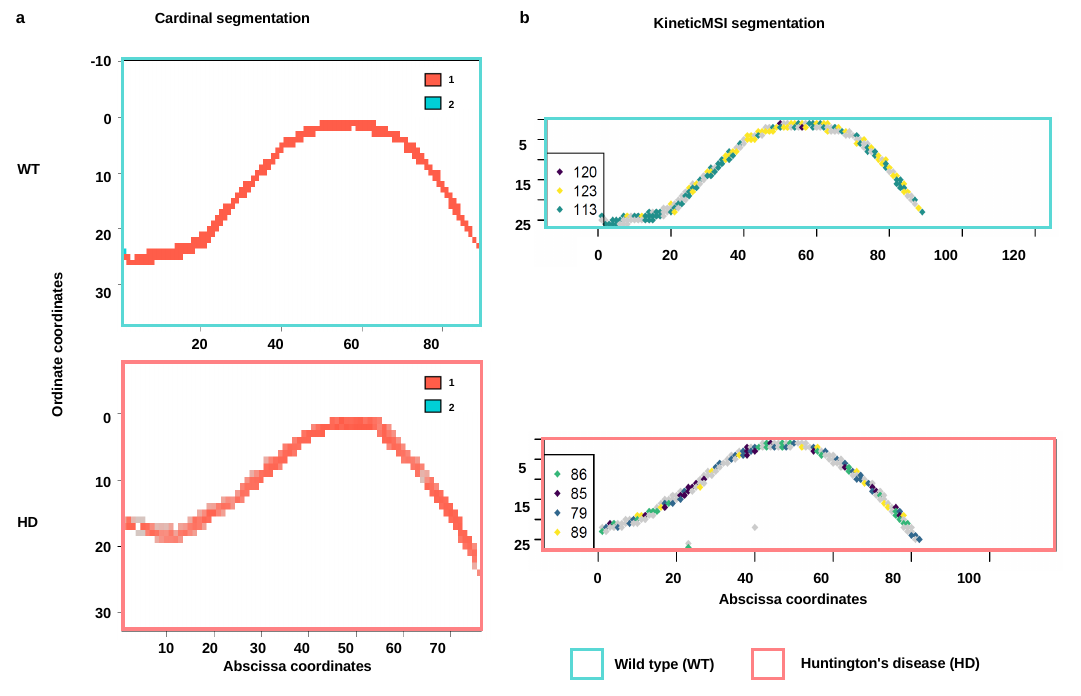

b
a
Cardinal segmentation
KineticMSI segmentation
-10
0
10
20
30
20 40 60 80
1
2
5
15
WT
25
0 20 40 60 80 100 120
Ordinate coordinates
0
10 20 30 40 50 60 70
1
2
5
0 20 40 60 80 100
10
15
HD
25
20
Abscissa coordinates
30
Huntington's disease (HD)
Wild type (WT)
Abscissa coordinates
