## Supplement Table 1 for "KineticMSI, an R-based framework for relative quantification of spatial isotopic incorporation in mass spectrometry imaging experiments"

| Annotation | Observed mass | Theoritical mass | Delta ppm | Molecular formula | Reference |
|---|---|---|---|---|---|
| AA | 303.23311 | 303.233 | -0.362757352 | C20H32O2 | https://hmdb.ca/metabolites/HMDB0001043 |
| DHA | 327.23317 | 327.233 | -0.519507507 | C22H32O2 | https://hmdb.ca/metabolites/HMDB0002183 |
| AA | 331.26479 | 331.264 | -2.384804869 | C22H36O2 | https://hmdb.ca/metabolites/HMDB0002226 |
| CPA 16:0 | 391.22681 | 391.2255 | -3.348452491 | C19H37O6P | https://hmdb.ca/metabolites/HMDB0007003 |
| CPA 18:1 | 417.2411 | 417.24291 | 4.338000615 | C21H39O6P | https://hmdb.ca/metabolites/HMDB0007005 |
| LPA 18:1 | 435.25286 | 435.2517 | -2.66512457 | C21H41O7P | https://hmdb.ca/metabolites/HMDB0007855 |
| LPA 18:0 | 437.26866 | 437.2674 | -2.881531987 | C21H43O7P | https://hmdb.ca/metabolites/HMDB0007854 |
| LPE 17:2 | 462.30015 | 462.299 | -2.48756757 | C23H46NO6P | https://www.lipidmaps.org/resources/tools/bulk_structure_searches.php?database=COMP_DB |
| LPG 15:3 | 463.21275 | 463.2103 | -5.289174269 | C21H37O9P | https://www.lipidmaps.org/resources/tools/bulk_structure_searches.php?database=COMP_DB |
| LPE 18:0 | 480.31202 | 480.3096 | -5.038416888 | C23H48NO7P | https://hmdb.ca/metabolites/HMDB0011130 |
| LPS O-18:2 | 506.2915 | 506.288818 | 5.297371588 | C24H45NO8P | http://www.swisslipids.org/#/entity/SLM:000115497/ |
| LPE 22:6 | 524.28031 | 524.278259 | 3.91204473 | C27H44NO7P | http://www.swisslipids.org/#/entity/SLM:000068676/ |
| PE 21:4 | 528.2756 | 528.273193 | 4.556354613 | C26H44NO8P | http://www.swisslipids.org/#/entity/SLM:000069251/ |
| PE 21:1 | 534.32255 | 534.320129 | 4.53099157 | C26H50NO8P | http://www.swisslipids.org/#/entity/SLM:000057082/ |
| LPI 14:3 | 537.19623 | 537.1978 | -2.922573399 | C23H39O12P | https://www.lipidmaps.org/resources/tools/bulk_structure_searches.php?database=COMP_DB |
| ST 19:1 | 540.28144 | 540.2814 | 0.074035493 | C27H43NO10 | https://www.lipidmaps.org/resources/tools/bulk_structure_searches.php?database=COMP_DB |
| LPS 20:3 | 546.28336 | 546.2838 | -0.805442153 | C26H46NO9P | https://www.lipidmaps.org/resources/tools/bulk_structure_searches.php?database=COMP_DB |
| PE 23:6 | 552.27507 | 552.2732 | 3.386005332 | C28H44NO8P | https://www.lipidmaps.org/resources/tools/bulk_structure_searches.php?database=COMP_DB |
| Unknown | 553.19066 | | | | |
| LPG 22:6 | 555.27498 | 555.2729 | 3.745905842 | C28H45O9P | https://www.lipidmaps.org/resources/tools/bulk_structure_searches.php?database=COMP_DB |
| CerP 28:5 | 556.30692 | 556.3045 | 4.350135582 | C28H48NO8P | https://www.lipidmaps.org/resources/tools/bulk_structure_searches.php?database=COMP_DB |
| ST 28:1 | 559.29659 | 559.2946 | 3.558053305 | C28H48O9S | https://www.lipidmaps.org/resources/tools/bulk_structure_searches.php?database=COMP_DB |
| LPS 21:3 | 560.29994 | 560.2994 | 0.963770441 | C27H48NO9P | https://www.lipidmaps.org/resources/tools/bulk_structure_searches.php?database=COMP_DB |
| LPI 16:0 | 571.2911 | 571.2889 | 3.850941266 | C25H49O12P | http://www.hmdb.ca/metabolites/HMDB0061695 |
| LPI O-17:0 | 571.324 | 571.3253 | 2 | C26H53O11P | https://www.lipidmaps.org/resources/tools/bulk_structure_searches.php?database=COMP_DB |
| LPS 22:4 | 572.29862 | 572.2994 | 1.362922973 | C28H48NO9P | Lipidmaps |
| LPI O-18:3 | 579.2968 | 579.294 | 4.833469706 | C27H49O11P | https://www.lipidmaps.org/resources/tools/bulk_structure_searches.php?database=COMP_DB |
| LPI O-18:2 | 581.3117 | 581.3096 | 3.612532805 | C27H51O11P | https://www.lipidmaps.org/resources/tools/bulk_structure_searches.php?database=COMP_DB |
| ST 30:2 | 585.31245 | 585.3103 | 3.673265275 | C30H50O9S | https://www.lipidmaps.org/resources/tools/bulk_structure_searches.php?database=COMP_DB |
| ST 30:1 | 587.32755 | 587.3259 | 2.80934316 | C30H52O9S | https://www.lipidmaps.org/resources/tools/bulk_structure_searches.php?database=COMP_DB |
| Unknown | 595.20124 | Unknown | Unknown | Unknown | Unknown |
| LPI 18:1 | 597.3066 | 597.3045 | 3.515794708 | C27H51O12P | http://www.hmdb.ca/metabolites/HMDB0061693 |
| LPI 18:0 | 599.323 | 599.3202 | 4.671959997 | C27H53O12P | http://www.hmdb.ca/metabolites/HMDB0240261 |
| PG 23:5 | 601.28035 | 601.2783 | 3.409402934 | C29H47O11P | https://www.lipidmaps.org/resources/tools/bulk_structure_searches.php?database=COMP_DB |
| Unknown | 603.16887 | Unknown | Unknown | Unknown | Unknown |
| LPI 20:4 | 619.28985 | 619.2889 | 1.534017484 | C29H49O12P | http://www.hmdb.ca/metabolites/HMDB0061690 |
| GlcCer d30:0 | 642.49599 | 642.495 | -1.540868022 | C36H69NO8 | https://hmdb.ca/metabolites/HMDB0012320 |
| CerP 36:1 | 644.5047 | 644.5025 | 3.413485595 | C36H72NO6P | https://hmdb.ca/metabolites/HMDB0010701 |
| PA 32:0 | 647.46489 | 647.4657 | -1.251031522 | C35H69O8P | http://www.hmdb.ca/metabolites/HMDB0000674 |
| ST 30:0 | 648.38288 | 648.3787 | 6.446849657 | C32H59NO10S | https://www.lipidmaps.org/resources/tools/bulk_structure_searches.php?database=COMP_DB |
| LPA 33:1 | 661.4829 | 661.4814 | 2.267637457 | C36H71O8P | https://www.lipidmaps.org/resources/tools/bulk_structure_searches.php?database=COMP_DB |
| CerP 38:3 | 668.50215 | 668.5025 | 0.523558252 | C38H72NO6P | Lipidmaps |
| GlcCer 32:1 | 670.52591 | 670.526 | 0.134222983 | C38H73NO8 | https://hmdb.ca/metabolites/HMDB0012321 |
| PA 34:1 | 673.48407 | 673.4814 | 3.964474743 | C37H71O8P | http://www.hmdb.ca/metabolites/HMDB0007858 |
| PA O-36:2 | 685.519 | 685.5178 | -1.750501592 | C39H75O7P | Lipimaps |
| PA 35:1 | 687.49767 | 687.497 | -0.974549707 | C38H73O8P | https://hmdb.ca/metabolites/HMDB0115093 |
| PA 36:4 | 695.4691 | 695.4657 | 4.888810476 | C39H69O8P | https://hmdb.ca/metabolites/HMDB0114880 |
| PA 36:2 | 699.49825 | 699.497 | 1.786998372 | C39H73O8P | https://hmdb.ca/metabolites/HMDB0007861 |
| PE O-34:1 | 700.52945 | 700.529 | -0.642371693 | C39H76NO7P | https://hmdb.ca/metabolites/HMDB0011342 |
| PA O-36:2 | 701.51243 | 701.5127 | -0.384882555 | C39H75O8P | http://www.hmdb.ca/metabolites/HMDB0114789 |
| SM d35:0 | 715.5783 | 715.5759 | 3.312017496 | C40H81N2O6P | https://hmdb.ca/metabolites/HMDB0240620 |
| PE 34:1 | 716.52563 | 716.5236 | 2.83312371 | C39H76NO8P | lipidmaps |
| PE 34:0 | 718.54237 | 718.5392 | 4.411728685 | C39H78NO8P | Lipidmaps |
| PA 38:6 | 719.46653 | 719.4657 | 1.15363387 | C41H69O8P | https://hmdb.ca/metabolites/HMDB0115179 |
| PA 38:5 | 721.48103 | 721.4814 | -0.512833733 | C41H71O8P | http://www.hmdb.ca/metabolites/HMDB0115126 |
| PA O-39:5 | 721.51776 | 721. 51776 | 0 | C42H73O7P | http://www.swisslipids.org/#/entity/SLM:000061661/ |
| PE P-36:4 | 722.5122 | 722.513 | 0 | C41H74NO7P | https://hmdb.ca/metabolites/HMDB0011444 |
| PA O-38:5 | 723.49971 | 723.497 | 3.74569625 | C41H73O8P | https://hmdb.ca/metabolites/HMDB0115100 |
| PA 38:3 | 725.51392 | 725.516 | 2.866925057 | C41H75O8P | https://hmdb.ca/metabolites/HMDB0115072 |
| PE P-36:2 | 726.54356 | 726.5443 | -1.018520137 | C41H78NO7P | https://hmdb.ca/metabolites/HMDB0011441 |
| PE P-36:1 | 728.56279 | 728.56 | 3.829471835 | C41H80NO7P | http://www.hmdb.ca/metabolites/HMDB0011348 |
| PA 38:1 | 729.54289 | 729.547 | 5.633632926 | C41H79O8P | https://hmdb.ca/metabolites/HMDB0114810 |
| PE-Cer 38:2 | 729.55322 | 729.5552 | 2.713982438 | C40H79N2O7P | Lipidmaps |

| | | | | | |
|---|---|---|---|---|---|
| PE 36:4 | 738.50851 | 738.5079 | 0.82598981 | C41H74NO8P | https://lipidmaps.org/data/chemdb_lm_text_ontology.php?ABBREV=PE%2036:4 |
| PE 36:2 | 742.5418 | 742.5392 | 3.501498641 | C41H78NO8P | https://lipidmaps.org/resources/tools/bulk-structure-search/7e0de5e2-d978-4b12-a130-0d272b294564 |
| PE 36:1 | 744.55769 | 744.5549 | 3.747205209 | C41H80NO8P | https://lipidmaps.org/data/chemdb_lm_text_ontology.php?ABBREV=PE%2036:1 |
| PE P-38:6 | 746.51294 | 746.513 | 0.080373684 | C43H74NO7P | https://hmdb.ca/metabolites/HMDB0005780 |
| PA 40:6 | 747.49733 | 747.497 | 0.441473344 | C43H73O8P | https://hmdb.ca/metabolites/HMDB0114942 |
| HexCer 36:6 | 748.50572 | 748.5005 | -6.973943237 | C42H71NO10 | Lipidmaps |
| CerP 43:6 | 748.52713 | 748.529 | 2.498233201 | C43H76NO7P | https://www.lipidmaps.org/resources/tools/bulk_structure_searches.php?database=COMP_DB |
| PE P-38:4 | 750.54634 | 750.5443 | 2.718027437 | C43H78NO7P | https://hmdb.ca/metabolites/HMDB0011386 |
| CerP 43:3 | 754.57611 | 754.5756 | 0.675876612 | C43H82NO7P | https://hmdb.ca/metabolites/HMDB0011357 |
| PA 40:2 | 755.56493 | 755.563 | -2.554386597 | C43H81O8P | https://hmdb.ca/metabolites/HMDB0115277 |
| PE P-38:1 | 756.59434 | 756.5913 | 4.018021354 | C43H84NO7P | https://hmdb.ca/metabolites/HMDB0011381 |
| PS 34:1 | 760.51672 | 760.5134 | 4.365472061 | C40H76NO10P | https://hmdb.ca/metabolites/HMDB0012357 |
| PE 38:6 | 762.51036 | 762.5079 | 3.226196083 | C43H74NO8P | https://lipidmaps.org/data/chemdb_lm_text_ontology.php?ABBREV=PE%2038:6 |
| PE 38:5 | 764.5226 | 764.5236 | 1.308004096 | C43H76NO8P | https://lipidmaps.org/data/chemdb_lm_text_ontology.php?ABBREV=PE%2038:6 |
| PE 38:4 | 766.54141 | 766.5392 | 2.88308804 | C43H78NO8P | https://lipidmaps.org/data/chemdb_lm_text_ontology.php?ABBREV=PE%2038:4 |
| PE O-40:8 | 772.52965 | 772.5287 | 1.229727776 | C45H76NO7P | https://lipidmaps.org/data/chemdb_lm_text_ontology.php?ABBREV=PE%20O-40:8 |
| PG 36:2 | 773.53403 | 773.5338 | 0.297336716 | C42H79O10P | https://hmdb.ca/metabolites/HMDB0010605 |
| PE P-40:6 | 774.54519 | 774.5443 | 1.149062746 | C45H78NO7P | https://hmdb.ca/metabolites/HMDB0009645 |
| PG 36:1 | 775.54946 | 775.551 | 1.985685016 | C42H81O10P | https://hmdb.ca/metabolites/HMDB0010632 |
| PE P-40:5 | 776.56289 | 776.56 | 3.721541156 | C45H80NO7P | https://hmdb.ca/metabolites/HMDB0011393 |
| PE P-40:4 | 778.5743 | 778.576 | 2.183473418 | C45H82NO7P | https://hmdb.ca/metabolites/HMDB0011391 |
| PS 36:2 | 786.53181 | 786.5291 | 3.445517782 | C42H78NO10P | https://hmdb.ca/metabolites/HMDB0112570 |
| PE 40:7 | 788.52184 | 788.5236 | -2.232019435 | C45H76NO8P | https://lipidmaps.org/data/chemdb_lm_text_ontology.php?ABBREV=PE%2040:7 |
| CerP 42:3 | 788.54662 | 788.5447 | 2.434865138 | C42H80NO10P | https://www.lipidmaps.org/resources/tools/bulk_structure_searches.php?database=COMP_DB |
| PE 40:6 | 790.5415 | 790.5392 | 2.909406643 | C45H78NO8P | https://lipidmaps.org/data/chemdb_lm_text_ontology.php?ABBREV=PE%2040:6 |
| PE 40:5 | 792.55435 | 792.5549 | 0.693958236 | C45H80NO8P | https://lipidmaps.org/data/chemdb_lm_text_ontology.php?ABBREV=PE%2040:5 |
| PE 40:4 | 794.57254 | 794.5705 | 2.567424791 | C45H82NO8P | https://lipidmaps.org/data/chemdb_lm_text_ontology.php?ABBREV=PE%2040:4 |
| ST d36:1 | 806.54438 | 806.5458 | -1.760594377 | C42H81NO11S | http://www.hmdb.ca/metabolites/HMDB0012314 |
| PI 32:0 | 809.51797 | 809.5186 | -0.778240302 | C41H79O13P | https://hmdb.ca/metabolites/HMDB0009778 |
| PS 38:4 | 810.53191 | 810.5291 | 3.466871208 | C44H78NO10P | https://hmdb.ca/metabolites/HMDB0112497 |
| PS 38:2 | 814.56335 | 814.56 | -4.112649774 | C44H82NO10P | https://hmdb.ca/metabolites/HMDB0112393 |
| PG 38:3 | 815.54064 | 815.5444 | -4.610417287 | C44H81O11P | https://www.lipidmaps.org/resources/tools/bulk_structure_searches.php?database=COMP_DB |
| HexCer 38:2 | 816.58363 | 816.5843 | -0.820490916 | C44H83NO12 | https://www.lipidmaps.org/resources/tools/bulk_structure_searches.php?database=COMP_DB |
| PI O-34:1 | 821.55281 | 821.5549 | -2.543956588 | C43H83O12P | https://www.lipidmaps.org/resources/tools/bulk_structure_searches.php?database=COMP_DB |
| PI-Cer 36:1 | 822.5478 | 822.5502 | -2.91775505 | C42H82NO12P | https://www.lipidmaps.org/resources/tools/bulk_structure_searches.php?database=COMP_DB |
| PS 39:4 | 824.54503 | 824.5447 | -0.400220873 | C45H80NO10P | Lipidmaps |
| PE 43:7 | 830.57379 | 830.5705 | 3.961132739 | C48H82NO8P | https://lipidmaps.org/resources/tools/bulk-structure-search/7e0de5e2-d978-4b12-a130-0d272b294564 |
| PS 40:7 | 832.51636 | 832.5134 | 3.555498326 | C46H76NO10P | Lipidmaps |
| PS 40:6 | 834.5261 | 834.5291 | -3.59484169 | C46H78NO10P | https://hmdb.ca/metabolites/HMDB0112421 |
| PI-Cer 38:1 | 834.58413 | 834.5866 | -2.959549075 | C44H86NO11P | https://www.lipidmaps.org/resources/tools/bulk_structure_searches.php?database=COMP_DB |
| PI 34:0 | 837.55039 | 837.5499 | -0.585039769 | C43H83O13P | https://hmdb.ca/metabolites/HMDB0009781 |
| PS 40:4 | 838.56256 | 838.5604 | 2.575843076 | C46H82NO10P | https://hmdb.ca/metabolites/HMDB0112665 |
| PG 40:4 | 841.55722 | 841.56 | -3.303388944 | C46H83O11P | https://www.lipidmaps.org/resources/tools/bulk_structure_searches.php?database=COMP_DB |
| PI O-36:1 | 849.58421 | 849.5862 | -2.342316765 | C45H87O12P | https://www.lipidmaps.org/resources/tools/bulk_structure_searches.php?database=COMP_DB |
| PS O-42:5 | 850.59795 | 850.596741 | 1.421355081 | C48H85NO9P | http://www.swisslipids.org/#/entity/SLM:000115714/ |
| PE O-46:9 | 854.60924 | 854.6069 | 2.738100991 | C51H86NO7P | https://www.lipidmaps.org/resources/tools/bulk_structure_searches.php?database=COMP_DB |
| PS 42:9 | 856.51313 | 856.51313 | 0 | C48H76NO10P | https://hmdb.ca/metabolites/HMDB0012428 |
| PI 36:4 | 857.51857 | 857.5186 | -0.034984664 | C45H79O13P | http://www.hmdb.ca/metabolites/HMDB0009899 |
| SHexCer 40:2 | 860.59329 | 860.5927 | -0.685574024 | C46H87NO11S | https://www.lipidmaps.org/resources/tools/bulk_structure_searches.php?database=COMP_DB |
| PI 36:2 | 861.54852 | 861.55 | 1.717834136 | C45H83O13P | https://hmdb.ca/metabolites/HMDB0009875 |
| PI-Cer 40:1 | 862.61698 | 862.6179 | -1.066520878 | C46H90NO11P | https://www.lipidmaps.org/resources/tools/bulk_structure_searches.php?database=COMP_DB |
| PI O-36:2 | 863.56894 | 863.5655 | 3.98348475 | C45H85O13P | https://www.lipidmaps.org/resources/tools/bulk_structure_searches.php?database=COMP_DB |
| PG O-44 6 | 863.61297 | 863.61713 | -4.816949381 | C50H88O9P | http://www.swisslipids.org/#/entity/SLM:000507720/ |
| PG 44:12 | 865.5032 | 865.5025 | 0.808778715 | C50H75O10P | http://www.hmdb.ca/metabolites/HMDB0116605 |
| PG 42:5 | 867.5756 | 867.5757 | -0.115263717 | C48H85O11P | https://www.lipidmaps.org/resources/tools/bulk_structure_searches.php?database=COMP_DB |
| CerP 48:5 | 868.60666 | 868.6073 | -0.736811675 | C48H88NO10P | https://www.lipidmaps.org/resources/tools/bulk_structure_searches.php?database=COMP_DB |

| | | | | | |
|---|---|---|---|---|---|
| PG 42:4 | 869.59028 | 869.5913 | -1.172964817 | C48H87O11P | https://www.lipidmaps.org/resources/tools/bulk_structure_searches.php?database=COMP_DB |
| PI 37:4 | 871.53783 | 871.5342 | 4.165068909 | C46H81O13P | https://www.lipidmaps.org/resources/tools/bulk_structure_searches.php?database=COMP_DB |
| PI-Cer 38:6 | 872.49374 | 872.4931 | 0.733530156 | C44H76NO14P | https://www.lipidmaps.org/resources/tools/bulk_structure_searches.php?database=COMP_DB |
| SHexCer 41:2 | 874.61103 | 874.6084 | 3.007060074 | C47H89NO11S | https://www.lipidmaps.org/resources/tools/bulk_structure_searches.php?database=COMP_DB |
| PE O-48:12 | 876.59718 | 876.5971 | 0.091261995 | C53H84NO7P | https://www.lipidmaps.org/resources/tools/bulk_structure_searches.php?database=COMP_DB |
| SHexCer 41:1 | 876.62653 | 876.624 | 2.886072022 | C47H91NO11S | https://www.lipidmaps.org/resources/tools/bulk_structure_searches.php?database=COMP_DB |
| PS 44:12 | 878.50123 | 878.4978 | 3.904392248 | C50H74NO10P | https://hmdb.ca/metabolites/HMDB0012450 |
| SHexCer 40:1;O3 | 878.60577 | 878.6033 | 2.811280131 | C46H89NO12S | https://www.lipidmaps.org/resources/tools/bulk_structure_searches.php?database=COMP_DB |
| PI 38:6 | 881.52127 | 881.5186 | 3.028864053 | C47H79O13P | https://www.lipidmaps.org/resources/tools/bulk_structure_searches.php?database=COMP_DB |
| PI 38:5 | 883.5389 | 883.5342 | 5 | C47H81O13P | http://www.hmdb.ca/metabolites/HMDB0009844 |
| PI 38:4 | 885.55229 | 885.5499 | 2.698888002 | C47H83O13P | http://www.hmdb.ca/metabolites/HMDB0009914 |
| SHexCer 42:3 | 886.61017 | 886.6084 | 1.996371792 | C48H89NO11S | https://www.lipidmaps.org/resources/tools/bulk_structure_searches.php?database=COMP_DB |
| ST d42:2 | 888.62383 | 88.624 | 0.19 | C48H91NO11S | http://www.hmdb.ca/metabolites/HMDB0012318 |
| PG 44:8 | 889.55989 | 889.56 | -0.123656639 | C50H83O11P | https://www.lipidmaps.org/resources/tools/bulk_structure_searches.php?database=COMP_DB |
| SHexCer 42:1;O2 | 890.63839 | 890.6397 | -1.470852916 | C48H93NO11S | https://www.lipidmaps.org/resources/tools/bulk_structure_searches.php?database=COMP_DB |
| PG 44:7 | 891.57594 | 891.5757 | 0.269186341 | C50H85O11P | https://www.lipidmaps.org/resources/tools/bulk_structure_searches.php?database=COMP_DB |
| PI-Cer 41:1 | 892.6289 | 892.6284 | 0.560143504 | C47H92NO12P | https://www.lipidmaps.org/resources/tools/bulk_structure_searches.php?database=COMP_DB |
| PS 44:4 | 894.62122 | 894.623 | 1.98966492 | C50H90NO10P | https://hmdb.ca/metabolites/HMDB0112620 |
| PG 44:5 | 895.60216 | 895.607 | -5.404156064 | C50H89O11P | https://www.lipidmaps.org/resources/tools/bulk_structure_searches.php?database=COMP_DB |
| CerP 50:5 | 896.63953 | 896.6386 | 1.037207187 | C50H92NO10P | https://www.lipidmaps.org/resources/tools/bulk_structure_searches.php?database=COMP_DB |
| PI-Cer 42:3 | 902.60891 | 902.6128 | -4.309710653 | C48H90NO12P | https://www.lipidmaps.org/resources/tools/bulk_structure_searches.php?database=COMP_DB |
| SHexCer 43:2;O2 | 902.64286 | 902.6397 | 3.500843138 | C49H93NO11S | https://www.lipidmaps.org/resources/tools/bulk_structure_searches.php?database=COMP_DB |
| SHexCer 42:2;O3 | 904.62209 | 904.6189 | 3.526346841 | C48H91NO12S | https://www.lipidmaps.org/resources/tools/bulk_structure_searches.php?database=COMP_DB |
| PE O-50:11 | 906.63856 | 906.6382 | 0.39707129 | C55H90NO7P | Lipidmaps |
| PI 40:6 | 909.55376 | 909.5499 | 4.243857319 | C49H83O13P | https://www.lipidmaps.org/resources/tools/bulk_structure_searches.php?database=COMP_DB |
| PI 40:5 | 911.56067 | 911.5655 | -5.298577009 | C49H85O13P | https://www.lipidmaps.org/resources/tools/bulk_structure_searches.php?database=COMP_DB |
| PS 44:3 | 912.63525 | 912.6335 | 1.917527682 | C50H92NO11P | https://www.lipidmaps.org/resources/tools/bulk_structure_searches.php?database=COMP_DB |
| PG 46:10 | 913.55595 | 913.56 | -4.433206357 | C52H83O11P | https://www.lipidmaps.org/resources/tools/bulk_structure_searches.php?database=COMP_DB |
| PG 46:9 | 915.5775 | 915.5757 | 1.965976161 | C52H85O11P | https://www.lipidmaps.org/resources/tools/bulk_structure_searches.php?database=COMP_DB |
| PI-Cer 44:2 | 916.66758 | 916.6648 | 3.032733449 | C50H96NO11P | https://www.lipidmaps.org/resources/tools/bulk_structure_searches.php?database=COMP_DB |
| TG 58:14 | 917.66369 | 917.667 | 3.6069729 | C61H90O6 | https://hmdb.ca/metabolites/HMDB0055525 |
| PIP 35:1 | 929.51498 | 929.516183 | -1.294221684 | C44H81O16P2 | http://www.swisslipids.org/#/entity/SLM:000417578/ |
| PS 46:8 | 930.59034 | 930.5866 | 4.0189704 | C52H86NO11P | https://www.lipidmaps.org/resources/tools/bulk_structure_searches.php?database=COMP_DB |
| PS 46:7 | 932.60657 | 932.6022 | 4.685813523 | C52H88NO11P | https://www.lipidmaps.org/resources/tools/bulk_structure_searches.php?database=COMP_DB |
| Hex2Cer 38:1 | 932.66396 | 932.668 | -4.331659283 | C50H95NO14 | https://www.lipidmaps.org/resources/tools/bulk_structure_searches.php?database=COMP_DB |
| PS 46:6 | 934.61943 | 934.6179 | 1.637032631 | C52H90NO11P | https://www.lipidmaps.org/resources/tools/bulk_structure_searches.php?database=COMP_DB |
| PE 50:9 | 940.64599 | 940.6437 | 2.434503096 | C55H92NO9P | Liidmaps |
| Hex2Cer 38:4 | 942.61653 | 942.616 | -0.562265016 | C50H89NO15 | https://www.lipidmaps.org/resources/tools/bulk_structure_searches.php?database=COMP_DB |
| PE 52:12 | 946.64795 | 946.6331 | -15.68717595 | C57H90NO8P | Lipidmaps |
| PGP 44:8 | 953.53204 | 953.531439 | 0.630288605 | C50H81O13P2 | http://www.swisslipids.org/#/entity/SLM:000400076/ |
| PS 48:9 | 956.60722 | 956.6022 | 5.247740388 | C54H88NO11P | https://www.lipidmaps.org/resources/tools/bulk_structure_searches.php?database=COMP_DB |
| PS 48:8 | 958.61981 | 958.6179 | 1.992451841 | C54H90NO11P | https://www.lipidmaps.org/resources/tools/bulk_structure_searches.php?database=COMP_DB |
| PS 48:6 | 962.6444 | 962.6492 | -4.986240055 | C54H94NO11P | https://www.lipidmaps.org/resources/tools/bulk_structure_searches.php?database=COMP_DB |
| PIP 38:4 | 965.5116 | 965.5162 | -4.764290853 | C47H84O16P2 | https://hmdb.ca/metabolites/HMDB0009922 |
| SHexCer 46:6 | 968.61559 | 968.6138 | 1.848001753 | C52H91NO13S | https://www.lipidmaps.org/resources/tools/bulk_structure_searches.php?database=COMP_DB |
| SHexCer 48:0 | 976.75032 | 976.7492 | 1.14666078 | C54H107NO11S | https://www.lipidmaps.org/resources/tools/bulk_structure_searches.php?database=COMP_DB |
| PIP 39:3 | 981.54214 | 981.5475 | -5.460764762 | C48H88O16P2 | https://www.lipidmaps.org/resources/tools/bulk_structure_searches.php?database=COMP_DB |
| PIP 40:7 | 987.50557 | 987.5005 | 5.134174616 | C49H82O16P2 | https://www.lipidmaps.org/resources/tools/bulk_structure_searches.php?database=COMP_DB |

| | | | | | | |
|---|---|---|---|---|---|---|
| PIP2 34:0 | 997.4842 | 997.4825 | 1.704290551 | C43H85O19P3 | https://hmdb.ca/metabolites/HMDB0010035 | |
| PI-Cer 46:6 | 1000.60632 | 1000.6037 | 2.61841926 | C52H92NO15P | https://www.lipidmaps.org/resources/tools/bulk_structure_searches.php?database=COMP_DB | |
| PIP 41:6 | 1003.53595 | 1003.53186 | 4.075605532 | C50H83O16P2 | http://www.swisslipids.org/#/entity/SLM:000400805/ | |
| MIPC 39:6 | 1048.55632 | 1048.5616 | -5.035469542 | C51H88NO19P | https://www.lipidmaps.org/resources/tools/bulk_structure_searches.php?database=COMP_DB | |
| PIP 44:3 | 1051.62114 | 1051.6257 | -4.336143554 | C53H98O16P2 | https://www.lipidmaps.org/resources/tools/bulk_structure_searches.php?database=COMP_DB | |
| PIP 45:3 | 1081.63727 | 1081.6363 | 0.896789429 | C54H100O17P2 | https://www.lipidmaps.org/resources/tools/bulk_structure_searches.php?database=COMP_DB | |
| GM3 d36:1 | 1179.73087 | 1179.7372 | 5.365601763 | C59H108N2O21 | https://www.lipidmaps.org/data/chemdb_lm_text_ontology.php?ABBREV=Hex(2)-NeuAc-Cer%2036:1 | O2 |
| PIP2 49:2 | 1219.67457 | 1219.6809 | -5.189882042 | C58H111O20P3 | https://www.lipidmaps.org/resources/tools/bulk_structure_searches.php?database=COMP_DB | |
| PIP2 49:1 | 1221.69742 | 1221.6965 | 0.753051187 | C58H113O20P3 | https://www.lipidmaps.org/resources/tools/bulk_structure_searches.php?database=COMP_DB | |
| PIP 59:7 | 1253.79845 | 1253.7979 | 0.438667189 | C68H120O16P2 | https://www.lipidmaps.org/resources/tools/bulk_structure_searches.php?database=COMP_DB | |
| GM2 d36:1 | 1382.80466 | 1382.8166 | 8.63455067 | C67H121N3O26 | https://www.lipidmaps.org/data/chemdb_lm_text_ontology.php?ABBREV=Hex(2)-HexNAc-NeuAc-Cer%2036:1 | O2 |
| CL 70:4 | 1427.99209 | 1427.9963 | -2.948186911 | C79H146O17P2 | https://www.lipidmaps.org/resources/tools/bulk_structure_searches.php?database=COMP_DB | |
| CL 72:7 | 1449.98123 | 1449.9806 | 0.434488572 | C81H144O17P2 | https://www.lipidmaps.org/resources/tools/bulk_structure_searches.php?database=COMP_DB | |
| CL 72:5 | 1454.00793 | 1454.0119 | -2.730376553 | C81H148O17P2 | https://www.lipidmaps.org/resources/tools/bulk_structure_searches.php?database=COMP_DB | |
| CL 72:4 | 1456.02231 | 1456.0276 | -3.633172888 | C81H150O17P2 | https://www.lipidmaps.org/resources/tools/bulk_structure_searches.php?database=COMP_DB | |
| CL 74:10 | 1471.96429 | 1471.965 | -0.482348425 | C83H142O17P2 | https://www.lipidmaps.org/resources/tools/bulk_structure_searches.php?database=COMP_DB | |
| CL 74:9 | 1473.97595 | 1473.9806 | -3.154722661 | C83H144O17P2 | https://www.lipidmaps.org/resources/tools/bulk_structure_searches.php?database=COMP_DB | |
| CL 74:8 | 1475.99313 | 1475.9963 | -2.147701861 | C83H146O17P2 | https://www.lipidmaps.org/resources/tools/bulk_structure_searches.php?database=COMP_DB | |
| CL 74:7 | 1478.00824 | 1478.0119 | -2.476299413 | C83H148O17P2 | https://www.lipidmaps.org/resources/tools/bulk_structure_searches.php?database=COMP_DB | |
| CL 76:11 | 1497.97515 | 1497.9806 | -3.638231363 | C85H144O17P2 | http://www.hmdb.ca/metabolites/HMDB0010272 | |
| CL 76:10 | 1499.99227 | 1499.9963 | -2.686673294 | C85H146O17P2 | https://www.lipidmaps.org/resources/tools/bulk_structure_searches.php?database=COMP_DB | |
| CL 78:12 | 1523.99373 | 1523.9963 | -1.6863558 | C87H146O17P2 | https://www.lipidmaps.org/resources/tools/bulk_structure_searches.php?database=COMP_DB | |
| CL 84:19 | 1537.92787 | 1537.91797 | -6.437274415 | C89H134O17P2 | http://swisslipids.org/#/entity/SLM:000570098/ | |
| GM1 36:0 | 1544.86661 | 1544.8694 | -1.805977903 | C73H131N3O31 | http://www.hmdb.ca/metabolites/HMDB0004856 | |
| CL 82:18 | 1569.97406 | 1569.9806 | -4.165656569 | C91H144O17P2 | https://hmdb.ca/metabolites/HMDB0059523 | |
| GM1 38:1 | 1572.89945 | 1572.9007 | -0.794710054 | C75H135N3O31 | http://www.hmdb.ca/metabolites/HMDB0004857 | |
| GA1 d50:6 | 1585.97658 | 1585.97278 | 2.396005813 | C82H142N2O27 | http://www.swisslipids.org/#/entity/SLM:000759248/ | |
| Globoside d46:1 | 1677.0087 | 1677.00952 | -0.48896562 | C82H151NO33 | http://www.swisslipids.org/#/entity/SLM:000752672/ | |
| GM1b d35:0 | 1751.94911 | 1751.94373 | 3.070874885 | C80H143N4O37 | http://www.swisslipids.org/#/entity/SLM:000774598/ | |
| GA1 d35:0 | 1752.96889 | 1752.96411 | 2.726809963 | C81H147N3O37 | http://www.swisslipids.org/#/entity/SLM:000759894/ | |
| GD2 d42:0 | 1759.02289 | 1759.0263 | -1.938572493 | C85H153N3O34 | http://www.hmdb.ca/metabolites/HMDB0011847 | |
| GD2 d44:7 | 1773.94591 | 1773.943271 | -1.487646219 | C86H140N4O34 | http://swisslipids.org/#/entity/SLM:000487699/ | |
| GT3 d38:0 | 1791.97952 | 1791.975 | 2.522356618 | C83H148N4O37 | http://www.hmdb.ca/metabolites/HMDB0012058 | |
| GA1 t52::5 | 1794.06007 | 1794.06738 | -4.074540389 | C90H158N2O33 | http://www.swisslipids.org/#/entity/SLM:000759505/ | |
| GD1 d36:1 | 1835.96146 | 1835.96484 | -1.840993861 | C84H146N4O39 | http://www.swisslipids.org/#/entity/SLM:000489463/ | |
| GM1b t42:4 | 1857.98946 | 1857.98547 | 2.147487192 | C87H149N4O38 | http://www.swisslipids.org/#/entity/SLM:000774485/ | |
| GD1 d38:1 | 1863.99995 | 1863.99609 | 2.070819794 | C86H150N4O39 | https://www.swisslipids.org/#/entity/SLM:000489464/ | |
| GT3 d41:1 | 1874.01661 | 1874.01683 | -0.11739489 | C88H151N4O38 | swisslipids.org/#/entity/SLM:000781996/ | |
| GD1a d40:4 | 1885.99023 | 1885.980445 | 5.188282851 | C88H148N4O39 | http://www.swisslipids.org/#/entity/SLM:000761962/ | |
| GD1a d38:3 | 1901.98377 | 1901.97534 | 4.432234121 | C88H148N4O40 | http://www.swisslipids.org/#/entity/SLM:000762492/ | |
