## Supplement Table 2 for "KineticMSI, an R-based framework for relative quantification of spatial isotopic incorporation in mass spectrometry imaging experiments"

| Fatty acid | *m/z* |
|---|---|
| 2:00 | 77.0476 |
| 3:00 | 91.0633 |
| 4:00 | 105.0789 |
| 5:00 | 119.0946 |
| 6:00 | 133.1102 |
| 7:00 | 147.1259 |
| 8:00 | 161.1415 |
| 9:00 | 175.1572 |
| 10:00 | 189.1728 |
| 11:00 | 203.1885 |
| 12:00 | 217.2041 |
| 13:00 | 231.2198 |
| 14:00 | 245.2354 |
| 14:01 | 243.2198 |
| 15:00 | 259.2511 |
| 15:01 | 257.2354 |
| 16:00 | 273.2667 |
| 16:01 | 271.2511 |
| 17:00 | 287.2824 |
| 17:01 | 285.2668 |
| 17:02 | 283.2511 |
| 18:00 | 301.2981 |
| 18:01 | 299.2825 |
| 18:02 | 297.2668 |
| 18:03 | 295.2512 |
| 18:04 | 293.2355 |
| 19:00 | 315.3137 |
| 20:00 | 329.3294 |
| 20:01 | 327.3138 |
| 20:02 | 325.2981 |
| 20:03 | 323.2825 |
