## Supplement Table 3 for "KineticMSI, an R-based framework for relative quantification of spatial isotopic incorporation in mass spectrometry imaging experiments"

| Lipid assignment | Anatomical_location | References | Assocaited neuronal function | References for neuronal functions |
|---|---|---|---|---|
| AA | Axonal_fatty_acids | (Piomelli et al., 2007) | Neurotransmitter signalling; pro-inflammatory mediators; enhanced hippocampal neurogenesis; Synaptic plasticity through induction of Long-term potentiation (LTP); Reduce neuronal excitability by regulation of glutamate N methyl d-aspartate (NMDA) receptor | (Piomelli et al., 2007; Rapoport 2008; Yang et al. 2012; Tokuda et al. 2014; Jang, Kim, and Hwang 2020) |
| ADA | Myelin_fatty_acids | (Martínez and Mougan 1998; VanRollins et al. 2008) | Myelination of neurons; Oligodendrocyte differentiation | (Martínez and Mougan 1998) |
| CerP361 | Synaptic_lipids | (Bajjalieh, Martin, and Floor 1989; Hoeferlin, Wijesinghe, and Chalfant 2013) | Neuronal proliferation ; Promotes cell survival; Increase in dendritic spine density ; Pro-inflammatory response | (Buccoliero, Bodennec, and Futerman 2002; Gómez-Muñoz 2006; Marcu and Chalfant 2007; Lamour and Chalfant 2005 ) |
| CerP423 | Synaptic_lipids | (Bajjalieh, Martin, and Floor 1989; Hoeferlin, Wijesinghe, and Chalfant 2013) | Neuronal proliferation ; Promotes cell survival; Increase in dendritic spine density ; Pro-inflammatory response | (Buccoliero, Bodennec, and Futerman 2002; Gómez-Muñoz 2006; Marcu and Chalfant 2007; Lamour and Chalfant 2005 ) |
| CerP436 | Synaptic_lipids | (Bajjalieh, Martin, and Floor 1989; Hoeferlin, Wijesinghe, and Chalfant 2013) | Neuronal proliferation ; Promotes cell survival; Increase in dendritic spine density ; Pro-inflammatory response | (Buccoliero, Bodennec, and Futerman 2002; Gómez-Muñoz 2006; Marcu and Chalfant 2007; Lamour and Chalfant 2005 ) |
| CerP485 | Synaptic_lipids | (Bajjalieh, Martin, and Floor 1989; Hoeferlin, Wijesinghe, and Chalfant 2013) | Neuronal proliferation ; Promotes cell survival; Increase in dendritic spine density ; Pro-inflammatory response | (Buccoliero, Bodennec, and Futerman 2002; Gómez-Muñoz 2006; Marcu and Chalfant 2007; Lamour and Chalfant 2005 ) |
| CerPEd381 | Myelin_lipids | (Panevska et al. 2019) | Myelination of neurons; Oligodendrocyte differentiation | (Panevska et al. 2019) |
| CL704 | Mitochondrial_lipids | (Mejia, Nguyen, and Hatch 2014) | Mitochondrial bioenergentics - ATP synthesis; Induces ER-stress mediated cell death | (Claypool and Koehler 2012; Paradies et al. 2014; Fan and Simmen 2019; Falabella et al. 2021) |
| CL724 | Mitochondrial_lipids | (Mejia, Nguyen, and Hatch 2014) | Mitochondrial bioenergentics - ATP synthesis; Induces ER-stress mediated cell death | (Claypool and Koehler 2012; Paradies et al. 2014; Fan and Simmen 2019; Falabella et al. 2021) |
| CL725 | Mitochondrial_lipids | (Mejia, Nguyen, and Hatch 2014) | Mitochondrial bioenergentics - ATP synthesis; Induces ER-stress mediated cell death | (Claypool and Koehler 2012; Paradies et al. 2014; Fan and Simmen 2019; Falabella et al. 2021) |
| CL727 | Mitochondrial_lipids | (Mejia, Nguyen, and Hatch 2014) | Mitochondrial bioenergentics - ATP synthesis; Induces ER-stress mediated cell death | (Claypool and Koehler 2012; Paradies et al. 2014; Fan and Simmen 2019; Falabella et al. 2021) |
| CL7410 | Mitochondrial_lipids | (Mejia, Nguyen, and Hatch 2014) | Mitochondrial bioenergentics - ATP synthesis; Induces ER-stress mediated cell death | (Claypool and Koehler 2012; Paradies et al. 2014; Fan and Simmen 2019; Falabella et al. 2021) |
| CL747 | Mitochondrial_lipids | (Mejia, Nguyen, and Hatch 2014) | Mitochondrial bioenergentics - ATP synthesis; Induces ER-stress mediated cell death | (Claypool and Koehler 2012; Paradies et al. 2014; Fan and Simmen 2019; Falabella et al. 2021) |
| CL748 | Mitochondrial_lipids | (Mejia, Nguyen, and Hatch 2014) | Mitochondrial bioenergentics - ATP synthesis; Induces ER-stress mediated cell death | (Claypool and Koehler 2012; Paradies et al. 2014; Fan and Simmen 2019; Falabella et al. 2021) |
| CL749 | Mitochondrial_lipids | (Mejia, Nguyen, and Hatch 2014) | Mitochondrial bioenergentics - ATP synthesis; Induces ER-stress mediated cell death | (Claypool and Koehler 2012; Paradies et al. 2014; Fan and Simmen 2019; Falabella et al. 2021) |
| CL7610 | Neuronal_cell_body | (Tian et al. 2019) | Mitochondrial bioenergentics - ATP synthesis; Induces ER-stress mediated cell death | (Claypool and Koehler 2012; Paradies et al. 2014; Fan and Simmen 2019; Falabella et al. 2021) |
| CL7611 | Mitochondrial_lipids | (Mejia, Nguyen, and Hatch 2014) | Mitochondrial bioenergentics - ATP synthesis; Induces ER-stress mediated cell death | (Claypool and Koehler 2012; Paradies et al. 2014; Fan and Simmen 2019; Falabella et al. 2021) |
| CL7812 | Mitochondrial_lipids | (Mejia, Nguyen, and Hatch 2014) | Mitochondrial bioenergentics - ATP synthesis; Induces ER-stress mediated cell death | (Claypool and Koehler 2012; Paradies et al. 2014; Fan and Simmen 2019; Falabella et al. 2021) |
| CL8218 | Neuronal_cell_body | (Tian et al. 2019) | Mitochondrial bioenergentics - ATP synthesis; Induces ER-stress mediated cell death | (Claypool and Koehler 2012; Paradies et al. 2014; Fan and Simmen 2019; Falabella et al. 2021) |
| CL8419 | Mitochondrial_lipids | (Mejia, Nguyen, and Hatch 2014) | Mitochondrial bioenergentics - ATP synthesis; Induces ER-stress mediated cell death | (Claypool and Koehler 2012; Paradies et al. 2014; Fan and Simmen 2019; Falabella et al. 2021) |
| CPA160 | Synaptic_lipids | (Schwarz et al. 2011) | Axonal growth; Promotes cell survival | (Fujiwara et al. 2003; Gotoh, Hotta, and Murakami-Murofushi 2010) |
| CPA181 | Synaptic_lipids | (Schwarz et al. 2011) | Axonal growth; Promotes cell survival | (Fujiwara et al. 2003; Gotoh, Hotta, and Murakami-Murofushi 2010) |
| DHA | Synaptic_fatty_acids | (Tanaka et al. 2012) | Promotes cell survival; Synaptic plasticity through induction of Long-term potentiation (LTP); ER stress protection; Neurotransmitter release; Alteration of membrane fluidity; Enhanced hippocampal neurogenesis - improve cognitive function | (Akbar and Kim 2002; Cao et al. 2009; Tanaka et al. 2012; Chaung et al. 2013; Chaung et al. 2013; Gharami et.al, 2015) |
| GA1350 | Synaptic_lipids | (Egawa et al. 2016; Sipione et al. 2020) | Membrane lipid raft-associated lipids that aid in signal transduction; Neurotransmitter release; Synaptogenesis; Synaptic plasticity through induction of Long-term potentiation (LTP) | (Egawa et al. 2016; Sipione et al. 2020) |
| GA1525 | Synaptic_lipids | (Egawa et al. 2016; Sipione et al. 2020) | Membrane lipid raft-associated lipids that aids in signal transduction, protein and lipid sorting and trafficking, neurotransmission signal transduction and synapse development; Enhanced synaptic plasticity and Neurotransmitter signalling in the hippocampus | (Egawa et al. 2016; Sipione et al. 2020) |
| GA1d506 | Synaptic_lipids | (Egawa et al. 2016; Sipione et al. 2020) | Membrane lipid raft-associated lipids that aids in signal transduction, protein and lipid sorting and trafficking, neurotransmission signal transduction and synapse development; Enhanced synaptic plasticity and Neurotransmitter signalling in the hippocampus | (Egawa et al. 2016; Sipione et al. 2020) |
| GA1t565 | Synaptic_lipids | (Egawa et al. 2016; Sipione et al. 2020) | Membrane lipid raft-associated lipids that aids in signal transduction, protein and lipid sorting and trafficking, neurotransmission signal transduction and synapse development; Enhanced synaptic plasticity and Neurotransmitter signalling in the hippocampus | (Egawa et al. 2016; Sipione et al. 2020) |
| GD1a383 | Synaptic_lipids | (Calderon, Attema, and DeVries 2002; van Echten-Deckert and Herget 2006; Egawa et al. 2016; Sipione et al. 2020) | Improves spatial learning and memory via enhanced synaptic plasticity in the CA3 hippocampal region; Synaptogenesis; Increase in dendritic spine density ; Axon-glial Interaction; Promotes axonal regeneration | (Kesner 2007;Schnaar 2010; Palmano et al. 2015) |
| GD1a404 | Synaptic_lipids | (Calderon, Attema, and DeVries 2002; van Echten-Deckert and Herget 2006; Egawa et al. 2016; Sipione et al. 2020) | Improves spatial learning and memory via enhanced synaptic plasticity in the CA3 hippocampal region; Synaptogenesis; Increase in dendritic spine density ; Axon-glial Interaction; Promotes axonal regeneration | (Kesner 2007;Schnaar 2010; Palmano et al. 2015) |
| GD1d361 | Synaptic_lipids | (Calderon, Attema, and DeVries 2002; van Echten-Deckert and Herget 2006; Egawa et al. 2016; Sipione et al. 2020) | Improves spatial learning and memory via enhanced synaptic plasticity in the CA3 hippocampal region; Synaptogenesis; Increase in dendritic spine density ; Axon-glial Interaction; Promotes axonal regeneration | (Kesner 2007;Schnaar 2010; Palmano et al. 2015) |
| GD2d420 | Synaptic_lipids | (Egawa et al. 2016; Sipione et al. 2020) | Improves spatial learning and memory via enhanced synaptic plasticity in the CA3 hippocampal region; Synaptogenesis; Increase in dendritic spine density ; Axon-glial Interaction; Promotes axonal regeneration | (Kesner 2007;Schnaar 2010; Palmano et al. 2015) |

| | | | | |
|---|---|---|---|---|
| GlcCer321 | Axonal_lipids | (Radin et al. 1972; van Echten-Deckert and Herget 2006) | Axonal growth; Enhanced hippocampal neurogenesis - improve cognitive function | (Buccoliero, Bodennec, and Futerman 2002) |
| Globosided461 | Synaptic_lipids | (Egawa et al. 2016; Sipione et al. 2020) | Membrane lipid raft-associated lipids that aid in signal transduction | (Kraft 2017) |
| Globosided406 | Synaptic_lipids | (Egawa et al. 2016; Sipione et al. 2020) | Membrane lipid raft-associated lipids that aid in signal transduction | (Kraft 2017) |
| Globosided561 | Synaptic_lipids | (Egawa et al. 2016; Sipione et al. 2020) | Membrane lipid raft-associated lipids that aid in signal transduction | (Kraft 2017) |
| GM1361 | Synaptic_lipids | (Calderon, Attema, and DeVries 2002; van Echten-Deckert and Herget 2006; Egawa et al. 2016; Sipione et al. 2020) | Membrane lipid raft-associated lipids that aid in signal transduction; Enhanced hippocampal neurogenesis - improve cognitive function; Neurotransmitter signalling; Anti-inflammatory response; Increase in dendritic spine density ; Axonal growth; Regulation of intracellular Ca2+ homeostasis; Reduce neuronal excitability by regulation of glutamate N methyl d-aspartate (NMDA) receptor; Synaptic plasticity through induction of Long-term potentiation (LTP); Improves spatial learning and memory via enhanced synaptic plasticity in the CA3 hippocampal region; Activation of brain derived neurotrophic factor signaling cascade. | (Lipartiti et al. 1991; Buccoliero, Bodennec, and Futerman 2002; Fujii et al. 2002; Zhang et al. 2005; Kesner 2007; Ohmi et al. 2011; Lim et al. 2011; Bisel, Pavone, and Calamai 2014; Palmano et al. 2015; Jiang et al. 2014; Schnaar, Gerardy-Schahn, and Hildebrandt 2014; Jiang et al. 2016; Ledeen and Wu 2015); Di Biase et al. 2020) |
| GM1381 | Synaptic_lipids | (Calderon, Attema, and DeVries 2002; van Echten-Deckert and Herget 2006; Egawa et al. 2016; Sipione et al. 2020) | Membrane lipid raft-associated lipids that aid in signal transduction; Enhanced hippocampal neurogenesis - improve cognitive function; Neurotransmitter signalling; Anti-inflammatory response; Increase in dendritic spine density ; Axonal growth; Regulation of intracellular Ca2+ homeostasis; Reduce neuronal excitability by regulation of glutamate N methyl d-aspartate (NMDA) receptor; Synaptic plasticity through induction of Long-term potentiation (LTP); Improves spatial learning and memory via enhanced synaptic plasticity in the CA3 hippocampal region; Activation of brain derived neurotrophic factor signaling cascade. | (Lipartiti et al. 1991; Buccoliero, Bodennec, and Futerman 2002; Fujii et al. 2002; Zhang et al. 2005; Kesner 2007; Ohmi et al. 2011; Lim et al. 2011; Bisel, Pavone, and Calamai 2014; Palmano et al. 2015; Jiang et al. 2014; Schnaar, Gerardy-Schahn, and Hildebrandt 2014; Jiang et al. 2016; Ledeen and Wu 2015); Di Biase et al. 2020) |
| GM1b424 | Synaptic_lipids | (Calderon, Attema, and DeVries 2002; van Echten-Deckert and Herget 2006; Egawa et al. 2016; Sipione et al. 2020) | Membrane lipid raft-associated lipids that aid in signal transduction; Enhanced hippocampal neurogenesis - improve cognitive function; Neurotransmitter signalling; Anti-inflammatory response; Increase in dendritic spine density ; Axonal growth; Regulation of intracellular Ca2+ homeostasis; Reduce neuronal excitability by regulation of glutamate N methyl d-aspartate (NMDA) receptor; Synaptic plasticity through induction of Long-term potentiation (LTP); Improves spatial learning and memory via enhanced synaptic plasticity in the CA3 hippocampal region; Activation of brain derived neurotrophic factor signaling cascade. | (Lipartiti et al. 1991; Buccoliero, Bodennec, and Futerman 2002; Fujii et al. 2002; Zhang et al. 2005; Kesner 2007; Ohmi et al. 2011; Lim et al. 2011; Bisel, Pavone, and Calamai 2014; Palmano et al. 2015; Jiang et al. 2014; Schnaar, Gerardy-Schahn, and Hildebrandt 2014; Jiang et al. 2016; Ledeen and Wu 2015); Di Biase et al. 2020) |
| GM1d382 | Synaptic_lipids | (Calderon, Attema, and DeVries 2002; van Echten-Deckert and Herget 2006; Egawa et al. 2016; Sipione et al. 2020) | Membrane lipid raft-associated lipids that aid in signal transduction; Enhanced hippocampal neurogenesis - improve cognitive function; Neurotransmitter signalling; Anti-inflammatory response; Increase in dendritic spine density ; Axonal growth; Regulation of intracellular Ca2+ homeostasis; Reduce neuronal excitability by regulation of glutamate N methyl d-aspartate (NMDA) receptor; Synaptic plasticity through induction of Long-term potentiation (LTP); Improves spatial learning and memory via enhanced synaptic plasticity in the CA3 hippocampal region; Activation of brain derived neurotrophic factor signaling cascade. | (Lipartiti et al. 1991; Buccoliero, Bodennec, and Futerman 2002; Fujii et al. 2002; Zhang et al. 2005; Kesner 2007; Ohmi et al. 2011; Lim et al. 2011; Bisel, Pavone, and Calamai 2014; Palmano et al. 2015; Jiang et al. 2014; Schnaar, Gerardy-Schahn, and Hildebrandt 2014; Jiang et al. 2016; Ledeen and Wu 2015); Di Biase et al. 2020) |
| GM2d361 | Synaptic_lipids | (Egawa et al. 2016; Sipione et al. 2020) | Membrane lipid raft-associated lipids that aid in signal transduction; Anti-inflammatory response; Myelination of neurons; Axonal growth, Enhanced hippocampal neurogenesis - improve cognitive function; Neurotransmitter signalling | (Lipartiti et al. 1991; Buccoliero, Bodennec, and Futerman 2002; Fujii et al. 2002; Zhang et al. 2005; Kesner 2007; Ohmi et al. 2011; Lim et al. 2011; Bisel, Pavone, and Calamai 2014; Palmano et al. 2015; Jiang et al. 2014; Schnaar, Gerardy-Schahn, and Hildebrandt 2014; Jiang et al. 2016; Ledeen and Wu 2015); Di Biase et al. 2020) |
| GM3d361 | Astrocyte_lipids | (Sipione et al. 2020) | Axonal growth; Neurotransmitter signalling; Enhanced hippocampal neurogenesis - improve cognitive function | |
| GT3411 | Synaptic_lipids | (Egawa et al. 2016; Sipione et al. 2020) | Axon-glial Interaction; Reduce neuronal excitability by regulation of glutamate N methyl d-aspartate (NMDA) receptor; Regulation of intracellular Ca2+ homeostasis | |
| GT3d380 | Synaptic_lipids | (Egawa et al. 2016; Sipione et al. 2020) | Axon-glial Interaction; Reduce neuronal excitability by regulation of glutamate N methyl d-aspartate (NMDA) receptor; Regulation of intracellular Ca2+ homeostasis | |
| GT3d480 | Synaptic_lipids | (Egawa et al. 2016; Sipione et al. 2020) | Axon-glial Interaction; Reduce neuronal excitability by regulation of glutamate N methyl d-aspartate (NMDA) receptor; Regulation of intracellular Ca2+ homeostasis | |
| GT3d41 | Synaptic_lipids | (Egawa et al. 2016; Sipione et al. 2020) | Axon-glial Interaction; Reduce neuronal excitability by regulation of glutamate N methyl d-aspartate (NMDA) receptor; Regulation of intracellular Ca2+ homeostasis | |
| Hex2Cer325 | Myelin_lipids | (Fitzner et al. 2020) | Myelination of neurons | (Fitzner et al. 2020) |
| Hex2Cer384 | Myelin_lipids | (Fitzner et al. 2020) | Myelination of neurons | (Fitzner et al. 2020) |
| HexCer366 | Myelin_lipids | (Fitzner et al. 2020) | Myelination of neurons | (Fitzner et al. 2020) |
| IPC411 | Myelin_lipids | (Björkbom et al. 2010) | Alteration of membrane fluidity; Neurotransmitter signalling | (Björkbom et al. 2010) |
| LPA180 | Synaptic_lipids | (Schwarz et al. 2011) | Axonal growth, Enhanced hippocampal neurogenesis - improve cognitive function; Neurotransmitter signalling; Signal transduction; Neuronal proliferation ; axon growth; Axon-glial Interaction | (Moolenaar 1995; Sengupta et al. 2004; Matas-Rico et al. 2008; Yung et al. 2015; Ladrón de Guevara-Miranda et al. 2019; Geraldo et al. 2021) |
| LPA181 | Synaptic_lipids | (Schwarz et al. 2011) | Axonal growth, Enhanced hippocampal neurogenesis - improve cognitive function; Neurotransmitter signalling; signal transduction; Neuronal proliferation ; axon growth; Axon-glial Interaction | (Matas-Rico et al. 2008; Yung et al. 2015; Ladrón de Guevara-Miranda et al. 2019; Geraldo et al. 2021) |
| LPA331 | Synaptic_lipids | (Schwarz et al. 2011) | Axonal growth, Enhanced hippocampal neurogenesis - improve cognitive function; Neurotransmitter signalling; signal transduction; Neuronal proliferation ; axon growth; Axon-glial Interaction | (Matas-Rico et al. 2008; Yung et al. 2015; Ladrón de Guevara-Miranda et al. 2019; Geraldo et al. 2021) |
| LPE172 | Mitochondrial_lipids | (Osman, Voelker, and Langer 2011; Horvath and Daum 2013; Tasseva et al. 2013) | Alteration of membrane fluidity; Signal transduction | (Calzada, Onguka, and Claypool 2016) |
| LPE180 | Mitochondrial_lipids | (Osman, Voelker, and Langer 2011; Horvath and Daum 2013; Tasseva et al. 2013) | Alteration of membrane fluidity; Signal transduction | |
| LPE226 | Mitochondrial_lipids | (Osman, Voelker, and Langer 2011; Horvath and Daum 2013; Tasseva et al. 2013) | Alteration of membrane fluidity; Signal transduction | |

| | | | | |
|---|---|---|---|---|
| LPG153 | Mitochondrial_lipids | (Horvath and Daum 2013; Morita and Terada 2015) | Neuronal proliferation ; Signal transduction | (Farooqui and Horrocks 2006) |
| LPG224 | Mitochondrial_lipids | (Horvath and Daum 2013; Morita and Terada 2015) | Neuronal proliferation ; Signal transduction | (Farooqui and Horrocks 2006) |
| LPG226 | Mitochondrial_lipids | (Horvath and Daum 2013; Morita and Terada 2015) | Neuronal proliferation ; Signal transduction | (Farooqui and Horrocks 2006) |
| LPG245 | Mitochondrial_lipids | (Horvath and Daum 2013; Morita and Terada 2015) | Neuronal proliferation ; Signal transduction | (Farooqui and Horrocks 2006) |
| LPI143 | Synaptic_lipids | (Frere, Chang-Ileto, and Di Paolo 2012; Lauwers, Goodchild, and Verstreken 2016) | ER stress protection; Anti-inflammatory agent | (Gangadharan et al. 2013) |
| LPI160 | Synaptic_lipids | (Frere, Chang-Ileto, and Di Paolo 2012; Lauwers, Goodchild, and Verstreken 2016) | ER stress protection; Anti-inflammatory agent | (Kallendrusch et al. 2013; Masquelier et al. 2018; Minamihata et al. 2020) |
| LPI180 | Synaptic_lipids | (Frere, Chang-Ileto, and Di Paolo 2012; Lauwers, Goodchild, and Verstreken 2016) | ER stress protection; Anti-inflammatory agent | (Kallendrusch et al. 2013; Masquelier et al. 2018; Minamihata et al. 2020) |
| LPI181 | Synaptic_lipids | (Frere, Chang-Ileto, and Di Paolo 2012; Lauwers, Goodchild, and Verstreken 2016) | ER stress protection; Anti-inflammatory agent | (Kallendrusch et al. 2013; Masquelier et al. 2018; Minamihata et al. 2020) |
| LPI204 | Synaptic_lipids | (Frere, Chang-Ileto, and Di Paolo 2012; Lauwers, Goodchild, and Verstreken 2016) | ER stress protection; Anti-inflammatory agent | (Kallendrusch et al. 2013; Masquelier et al. 2018; Minamihata et al. 2020) |
| LPIO170 | Synaptic_lipids | (Frere, Chang-Ileto, and Di Paolo 2012; Lauwers, Goodchild, and Verstreken 2016) | ER stress protection; Anti-inflammatory agent | (Kallendrusch et al. 2013; Masquelier et al. 2018; Minamihata et al. 2020) |
| LPIO182 | Synaptic_lipids | (Frere, Chang-Ileto, and Di Paolo 2012; Lauwers, Goodchild, and Verstreken 2016) | ER stress protection; Anti-inflammatory agent | (Kallendrusch et al. 2013; Masquelier et al. 2018; Minamihata et al. 2020) |
| LPIO183 | Synaptic_lipids | (Frere, Chang-Ileto, and Di Paolo 2012; Lauwers, Goodchild, and Verstreken 2016) | ER stress protection; Anti-inflammatory agent | (Kallendrusch et al. 2013; Masquelier et al. 2018; Minamihata et al. 2020) |
| LPS182 | Axonal_lipids | (Calderon, Attema, and DeVries 2002; Kim, Huang, and Spector 2014) | Signal transduction; Anti-inflammatory agent; Axonal growth | (Frasch and Bratton 2012) |
| LPS203 | Axonal_lipids | (Calderon, Attema, and DeVries 2002; Kim, Huang, and Spector 2014) | Signal transduction; Anti-inflammatory agent; Axonal growth | (Frasch and Bratton 2012) |
| LPSO202 | Axonal_lipids | (Calderon, Attema, and DeVries 2002; Kim, Huang, and Spector 2014) | Signal transduction; Anti-inflammatory agent; Axonal growth | (Frasch and Bratton 2012) |
| PA320 | Synaptic_lipids | (Schwarz et al. 2011) | Alteration of membrane fluidity; Neurotransmitter signalling; Signal transduction; ER stress protection; Axonal growth; Maintains lipid homeostasis | (Hite, Butterwick, and MacKinnon 2014; Raben and Barber 2017; Zhukovsky et al. 2019; Tanguy et al. 2019; Tanguy, Wang, and Vitale 2020) |
| PA341 | Synaptic_lipids | (Schwarz et al. 2011) | Alteration of membrane fluidity; Neurotransmitter signalling; Signal transduction; ER stress protection; Axonal growth; Maintains lipid homeostasis | (Hite, Butterwick, and MacKinnon 2014; Raben and Barber 2017; Zhukovsky et al. 2019; Tanguy et al. 2019; Tanguy, Wang, and Vitale 2020) |
| PA351 | Synaptic_lipids | (Schwarz et al. 2011) | Alteration of membrane fluidity; Neurotransmitter signalling; Signal transduction; ER stress protection; Axonal growth; Maintains lipid homeostasis | (Hite, Butterwick, and MacKinnon 2014; Raben and Barber 2017; Zhukovsky et al. 2019; Tanguy et al. 2019; Tanguy, Wang, and Vitale 2020) |
| PA362 | Synaptic_lipids | (Schwarz et al. 2011) | Alteration of membrane fluidity; Neurotransmitter signalling; Signal transduction; ER stress protection; Axonal growth; Maintains lipid homeostasis | (Hite, Butterwick, and MacKinnon 2014; Raben and Barber 2017; Zhukovsky et al. 2019; Tanguy et al. 2019; Tanguy, Wang, and Vitale 2020) |
| PA364 | Synaptic_lipids | (Schwarz et al. 2011) | Alteration of membrane fluidity; Neurotransmitter signalling; Signal transduction; ER stress protection; Axonal growth; Maintains lipid homeostasis | (Hite, Butterwick, and MacKinnon 2014; Raben and Barber 2017; Zhukovsky et al. 2019; Tanguy et al. 2019; Tanguy, Wang, and Vitale 2020) |
| PA384 | Synaptic_lipids | (Schwarz et al. 2011) | Alteration of membrane fluidity; Neurotransmitter signalling; Signal transduction; ER stress protection; Axonal growth; Maintains lipid homeostasis | (Hite, Butterwick, and MacKinnon 2014; Raben and Barber 2017; Zhukovsky et al. 2019; Tanguy et al. 2019; Tanguy, Wang, and Vitale 2020) |
| PA385 | Synaptic_lipids | (Schwarz et al. 2011) | Alteration of membrane fluidity; Neurotransmitter signalling; Signal transduction; ER stress protection; Axonal growth; Maintains lipid homeostasis | (Hite, Butterwick, and MacKinnon 2014; Raben and Barber 2017; Zhukovsky et al. 2019; Tanguy et al. 2019; Tanguy, Wang, and Vitale 2020) |
| PA386 | Synaptic_lipids | (Schwarz et al. 2011) | Alteration of membrane fluidity; Neurotransmitter signalling; Signal transduction; ER stress protection; Axonal growth; Maintains lipid homeostasis | (Hite, Butterwick, and MacKinnon 2014; Raben and Barber 2017; Zhukovsky et al. 2019; Tanguy et al. 2019; Tanguy, Wang, and Vitale 2020) |
| PA406 | Synaptic_lipids | (Schwarz et al. 2011) | Alteration of membrane fluidity; Neurotransmitter signalling; Signal transduction; ER stress protection; Axonal growth; Maintains lipid homeostasis | (Hite, Butterwick, and MacKinnon 2014; Raben and Barber 2017; Zhukovsky et al. 2019; Tanguy et al. 2019; Tanguy, Wang, and Vitale 2020) |
| PAO362 | Synaptic_lipids | (Schwarz et al. 2011) | Alteration of membrane fluidity; Neurotransmitter signalling; Signal transduction; ER stress protection; Axonal growth; Maintains lipid homeostasis | (Hite, Butterwick, and MacKinnon 2014; Raben and Barber 2017; Zhukovsky et al. 2019; Tanguy et al. 2019; Tanguy, Wang, and Vitale 2020) |
| PAO385 | Synaptic_lipids | (Schwarz et al. 2011) | Alteration of membrane fluidity; Neurotransmitter signalling; Signal transduction; ER stress protection; Axonal growth; Maintains lipid homeostasis | (Hite, Butterwick, and MacKinnon 2014; Raben and Barber 2017; Zhukovsky et al. 2019; Tanguy et al. 2019; Tanguy, Wang, and Vitale 2020) |
| PE214 | Mitochondrial_lipids | (Osman, Voelker, and Langer 2011;Tamura et al. 2012; Horvath and Daum 2013; Calzada et al. 2019) | Alteration of membrane fluidity; Signal transduction | |
| PE236 | Mitochondrial_lipids | (Osman, Voelker, and Langer 2011;Tamura et al. 2012; Horvath and Daum 2013; Calzada et al. 2019) | Alteration of membrane fluidity; Signal transduction | |
| PE340 | Axonal_lipids | (Merrill et al. 2017; Xie et al. 2020) | Promotes axonal regeneration; Neuronal proliferation ; Alteration of membrane fluidity; ER stress protection; Maintains lipid homeostasis; enhanced hippocampal neurogenesis; Synaptic plasticity through induction of Long-term potentiation (LTP); Reduce neuronal excitability by regulation of glutamate N methyl d-aspartate (NMDA) receptor; Activation of brain derived neurotrophic factor signaling cascade; Axonal growth | (Kreutzberger et al. 2017; Neumann et al. 2019; Che et al. 2020) |
| PE341 | Astrocyte_lipids | (Fitzner et al. 2020) | Promotes axonal regeneration; Neuronal proliferation ; Alteration of membrane fluidity; ER stress protection; Maintains lipid homeostasis; enhanced hippocampal neurogenesis; Synaptic plasticity through induction of Long-term potentiation (LTP); Reduce neuronal excitability by regulation of glutamate N methyl d-aspartate (NMDA) receptor; Activation of brain derived neurotrophic factor signaling cascade | (Kreutzberger et al. 2017; Neumann et al. 2019; Che et al. 2020) |
| PE361 | Axonal_lipids | (Neumann et al. 2019) | Promotes axonal regeneration; Neuronal proliferation ; Alteration of membrane fluidity; ER stress protection; Maintains lipid homeostasis; enhanced hippocampal neurogenesis; Synaptic plasticity through induction of Long-term potentiation (LTP); Reduce neuronal excitability by regulation of glutamate N methyl d-aspartate (NMDA) receptor; Activation of brain derived neurotrophic factor signaling cascade; Axonal growth | (Kreutzberger et al. 2017; Neumann et al. 2019; Che et al. 2020) |

| | | | | |
|---|---|---|---|---|
| PE362 | Mitochondrial_lipids | (Osman, Voelker, and Langer 2011;Tamura et al. 2012; Horvath and Daum 2013; Calzada et al. 2019) | Promotes axonal regeneration; Neuronal proliferation ; Alteration of membrane fluidity; ER stress protection; Maintains lipid homeostasis; enhanced hippocampal neurogenesis; Synaptic plasticity through induction of Long-term potentiation (LTP); Reduce neuronal excitability by regulation of glutamate N methyl d-aspartate (NMDA) receptor; Activation of brain derived neurotrophic factor signaling cascade | (Fu et al. 2011)(Kreutzberger et al. 2017)(Trentzsch et al. 2020)(Patel and Witt 2017)(Donoso et al. 2020)(Tasseva et al. 2013)(Emoto et al. 1996)(Yang et al. 2020); Che et al. 2020 |
| PE361 | Mitochondrial_lipids | (Osman, Voelker, and Langer 2011;Tamura et al. 2012; Horvath and Daum 2013; Calzada et al. 2019) | Promotes axonal regeneration; Neuronal proliferation ; Alteration of membrane fluidity; ER stress protection; Maintains lipid homeostasis; enhanced hippocampal neurogenesis; Synaptic plasticity through induction of Long-term potentiation (LTP); Reduce neuronal excitability by regulation of glutamate N methyl d-aspartate (NMDA) receptor; Activation of brain derived neurotrophic factor signaling cascade | (Fu et al. 2011)(Kreutzberger et al. 2017)(Trentzsch et al. 2020)(Patel and Witt 2017)(Donoso et al. 2020)(Tasseva et al. 2013)(Emoto et al. 1996)(Yang et al. 2020); Che et al. 2022 |
| PE384 | Mitochondrial_lipids | (Osman, Voelker, and Langer 2011;Tamura et al. 2012; Horvath and Daum 2013; Calzada et al. 2019) | Promotes axonal regeneration; Neuronal proliferation ; Alteration of membrane fluidity; ER stress protection; Maintains lipid homeostasis; enhanced hippocampal neurogenesis; Synaptic plasticity through induction of Long-term potentiation (LTP); Reduce neuronal excitability by regulation of glutamate N methyl d-aspartate (NMDA) receptor; Activation of brain derived neurotrophic factor signaling cascade | (Fu et al. 2011)(Kreutzberger et al. 2017)(Trentzsch et al. 2020)(Patel and Witt 2017)(Donoso et al. 2020)(Tasseva et al. 2013)(Emoto et al. 1996)(Yang et al. 2020); Che et al. 2023 |
| PE385 | Axonal_lipids | (Merrill et al. 2017) | Promotes axonal regeneration; Neuronal proliferation ; Alteration of membrane fluidity; ER stress protection; Maintains lipid homeostasis; enhanced hippocampal neurogenesis; Synaptic plasticity through induction of Long-term potentiation (LTP); Reduce neuronal excitability by regulation of glutamate N methyl d-aspartate (NMDA) receptor; Activation of brain derived neurotrophic factor signaling cascade; Axonal growth | (Fu et al. 2011)(Kreutzberger et al. 2017)(Trentzsch et al. 2020)(Patel and Witt 2017)(Donoso et al. 2020)(Tasseva et al. 2013)(Emoto et al. 1996)(Yang et al. 2020); Che et al. 2024 |
| PE404 | Mitochondrial_lipids | (Osman, Voelker, and Langer 2011;Tamura et al. 2012; Horvath and Daum 2013; Calzada et al. 2019) | Promotes axonal regeneration; Neuronal proliferation ; Alteration of membrane fluidity; ER stress protection; Maintains lipid homeostasis; enhanced hippocampal neurogenesis; Synaptic plasticity through induction of Long-term potentiation (LTP); Reduce neuronal excitability by regulation of glutamate N methyl d-aspartate (NMDA) receptor; Activation of brain derived neurotrophic factor signaling cascade | (Fu et al. 2011)(Kreutzberger et al. 2017)(Trentzsch et al. 2020)(Patel and Witt 2017)(Donoso et al. 2020)(Tasseva et al. 2013)(Emoto et al. 1996)(Yang et al. 2020); Che et al. 2025 |
| PE405 | Mitochondrial_lipids | (Osman, Voelker, and Langer 2011;Tamura et al. 2012; Horvath and Daum 2013; Calzada et al. 2019) | Promotes axonal regeneration; Neuronal proliferation ; Alteration of membrane fluidity; ER stress protection; Maintains lipid homeostasis; enhanced hippocampal neurogenesis; Synaptic plasticity through induction of Long-term potentiation (LTP); Reduce neuronal excitability by regulation of glutamate N methyl d-aspartate (NMDA) receptor; Activation of brain derived neurotrophic factor signaling cascade | (Fu et al. 2011)(Kreutzberger et al. 2017)(Trentzsch et al. 2020)(Patel and Witt 2017)(Donoso et al. 2020)(Tasseva et al. 2013)(Emoto et al. 1996)(Yang et al. 2020); Che et al. 2026 |
| PE407 | Mitochondrial_lipids | (Osman, Voelker, and Langer 2011;Tamura et al. 2012; Horvath and Daum 2013; Calzada et al. 2019) | Promotes axonal regeneration; Neuronal proliferation ; Alteration of membrane fluidity; ER stress protection; Maintains lipid homeostasis; enhanced hippocampal neurogenesis; Synaptic plasticity through induction of Long-term potentiation (LTP); Reduce neuronal excitability by regulation of glutamate N methyl d-aspartate (NMDA) receptor; Activation of brain derived neurotrophic factor signaling cascade | (Fu et al. 2011)(Kreutzberger et al. 2017)(Trentzsch et al. 2020)(Patel and Witt 2017)(Donoso et al. 2020)(Tasseva et al. 2013)(Emoto et al. 1996)(Yang et al. 2020); Che et al. 2028 |
| PE437 | Mitochondrial_lipids | (Osman, Voelker, and Langer 2011;Tamura et al. 2012; Horvath and Daum 2013; Calzada et al. 2019) | Promotes axonal regeneration; Neuronal proliferation ; Alteration of membrane fluidity; ER stress protection; Maintains lipid homeostasis; enhanced hippocampal neurogenesis; Synaptic plasticity through induction of Long-term potentiation (LTP); Reduce neuronal excitability by regulation of glutamate N methyl d-aspartate (NMDA) receptor; Activation of brain derived neurotrophic factor signaling cascade | (Fu et al. 2011)(Kreutzberger et al. 2017)(Trentzsch et al. 2020)(Patel and Witt 2017)(Donoso et al. 2020)(Tasseva et al. 2013)(Emoto et al. 1996)(Yang et al. 2020); Che et al. 2029 |
| PE384 | Axonal_lipids | (Kaya et al. 2017) | Promotes axonal regeneration; Neuronal proliferation ; Alteration of membrane fluidity; ER stress protection; Maintains lipid homeostasis; enhanced hippocampal neurogenesis; Synaptic plasticity through induction of Long-term potentiation (LTP); Reduce neuronal excitability by regulation of glutamate N methyl d-aspartate (NMDA) receptor; Activation of brain derived neurotrophic factor signaling cascade; Axonal growth | (Fu et al. 2011)(Kreutzberger et al. 2017)(Trentzsch et al. 2020)(Patel and Witt 2017)(Donoso et al. 2020)(Tasseva et al. 2013)(Emoto et al. 1996)(Yang et al. 2020); Che et al. 2030 |
| PE386 | Mitochondrial_lipids | (Osman, Voelker, and Langer 2011;Tamura et al. 2012; Horvath and Daum 2013; Calzada et al. 2019) | Promotes axonal regeneration; Neuronal proliferation ; Alteration of membrane fluidity; ER stress protection; Maintains lipid homeostasis; enhanced hippocampal neurogenesis; Synaptic plasticity through induction of Long-term potentiation (LTP); Reduce neuronal excitability by regulation of glutamate N methyl d-aspartate (NMDA) receptor; Activation of brain derived neurotrophic factor signaling cascade | (Fu et al. 2011)(Kreutzberger et al. 2017)(Trentzsch et al. 2020)(Patel and Witt 2017)(Donoso et al. 2020)(Tasseva et al. 2013)(Emoto et al. 1996)(Yang et al. 2020); Che et al. 2031 |
| PEO365 | Myelin_lipids | (Fitzner et al. 2020) | Alteration of membrane fluidity; Neurotransmitter signalling; Signal transduction; ER stress protection; Regulators of membrane trafficking and dynamics | (Hahnel et al. 1999; Messias et al. 2018; Wanders and Brites 2010) |
| PEO407_0 | Myelin_lipids | (Fitzner et al. 2020) | Alteration of membrane fluidity; Neurotransmitter signalling; Signal transduction; ER stress protection; Regulators of membrane trafficking and dynamics | (Hahnel et al. 1999; Messias et al. 2018; Wanders and Brites 2010) |
| PEO408 | Myelin_lipids | (Fitzner et al. 2020) | Alteration of membrane fluidity; Neurotransmitter signalling; Signal transduction; ER stress protection; Regulators of membrane trafficking and dynamics | (Hahnel et al. 1999; Messias et al. 2018; Wanders and Brites 2010) |
| PEO469 | Myelin_lipids | (Fitzner et al. 2020) | Alteration of membrane fluidity; Neurotransmitter signalling; Signal transduction; ER stress protection; Regulators of membrane trafficking and dynamics | (Hahnel et al. 1999; Messias et al. 2018; Wanders and Brites 2010) |
| PEO4812 | Myelin_lipids | (Fitzner et al. 2020) | Alteration of membrane fluidity; Neurotransmitter signalling; Signal transduction; ER stress protection; Regulators of membrane trafficking and dynamics | (Hahnel et al. 1999; Messias et al. 2018; Wanders and Brites 2010) |
| PEO5011 | Myelin_lipids | (Fitzner et al. 2020) | Alteration of membrane fluidity; Neurotransmitter signalling; Signal transduction; ER stress protection; Regulators of membrane trafficking and dynamics | (Hahnel et al. 1999; Messias et al. 2018; Wanders and Brites 2010) |

| | | | | |
|---|---|---|---|---|
| PEP362 | Myelin_lipids | (Fitzner et al. 2020) | Alteration of membrane fluidity; Neurotransmitter signalling; Signal transduction; ER stress protection; Regulators of membrane trafficking and dynamics | (Hahnel et al. 1999; Messias et al. 2018; Wanders and Brites 2010) |
| PEP363 | Myelin_lipids | (Fitzner et al. 2020) | Alteration of membrane fluidity; Neurotransmitter signalling; Signal transduction; ER stress protection; Regulators of membrane trafficking and dynamics | (Hahnel et al. 1999; Messias et al. 2018; Wanders and Brites 2010) |
| PEP364 | Myelin_lipids | (Fitzner et al. 2020) | Alteration of membrane fluidity; Neurotransmitter signalling; Signal transduction; ER stress protection; Regulators of membrane trafficking and dynamics | (Hahnel et al. 1999; Messias et al. 2018; Wanders and Brites 2010) |
| PEP384 | Myelin_lipids | (Fitzner et al. 2020) | Alteration of membrane fluidity; Neurotransmitter signalling; Signal transduction; ER stress protection; Regulators of membrane trafficking and dynamics | (Hahnel et al. 1999; Messias et al. 2018; Wanders and Brites 2010) |
| PEP386 | Myelin_lipids | (Fitzner et al. 2020) | Alteration of membrane fluidity; Neurotransmitter signalling; Signal transduction; ER stress protection; Regulators of membrane trafficking and dynamics | (Hahnel et al. 1999; Messias et al. 2018; Wanders and Brites 2010) |
| PEP405 | Myelin_lipids | (Fitzner et al. 2020) | Alteration of membrane fluidity; Neurotransmitter signalling; Signal transduction; ER stress protection; Regulators of membrane trafficking and dynamics | (Hahnel et al. 1999; Messias et al. 2018; Wanders and Brites 2010) |
| PEP424 | Myelin_lipids | (Fitzner et al. 2020) | Alteration of membrane fluidity; Neurotransmitter signalling; Signal transduction; ER stress protection; Regulators of membrane trafficking and dynamics | (Hahnel et al. 1999; Messias et al. 2018; Wanders and Brites 2010) |
| PEP406 | Myelin_lipids | (Fitzner et al. 2020) | Alteration of membrane fluidity; Neurotransmitter signalling; Signal transduction; ER stress protection; Regulators of membrane trafficking and dynamics | (Hahnel et al. 1999; Messias et al. 2018; Wanders and Brites 2010) |
| PG235 | Mitochondrial_lipids | (Horvath and Daum 2013; Morita and Terada 2015) | Neuroprotectant; Mitochondrial bioenergentics - ATP synthesis; Enhanced hippocampal neurogenesis - improve cognitive function | (Chen et al. 2018; Crotty et al. 2008; Donoso et al. 2020; Geribaldi-Doldán et al. 2019; Morita and Terada 2015; Murray and Fields 1998; Nie et al. 2010) |
| PG361 | Mitochondrial_lipids | (Horvath and Daum 2013; Morita and Terada 2015) | Neuroprotectant; Mitochondrial bioenergentics - ATP synthesis; Enhanced hippocampal neurogenesis - improve cognitive function | (Chen et al. 2018; Crotty et al. 2008; Donoso et al. 2020; Geribaldi-Doldán et al. 2019; Morita and Terada 2015; Murray and Fields 1998; Nie et al. 2010) |
| PG362 | Mitochondrial_lipids | (Horvath and Daum 2013; Morita and Terada 2015) | Neuroprotectant; Mitochondrial bioenergentics - ATP synthesis; Enhanced hippocampal neurogenesis - improve cognitive function | (Chen et al. 2018; Crotty et al. 2008; Donoso et al. 2020; Geribaldi-Doldán et al. 2019; Morita and Terada 2015; Murray and Fields 1998; Nie et al. 2010) |
| PG383 | Mitochondrial_lipids | (Horvath and Daum 2013; Morita and Terada 2015) | Neuroprotectant; Mitochondrial bioenergentics - ATP synthesis; Enhanced hippocampal neurogenesis - improve cognitive function | (Chen et al. 2018; Crotty et al. 2008; Donoso et al. 2020; Geribaldi-Doldán et al. 2019; Morita and Terada 2015; Murray and Fields 1998; Nie et al. 2010) |
| PG404 | Mitochondrial_lipids | (Horvath and Daum 2013; Morita and Terada 2015) | Neuroprotectant; Mitochondrial bioenergentics - ATP synthesis; Enhanced hippocampal neurogenesis - improve cognitive function | (Chen et al. 2018; Crotty et al. 2008; Donoso et al. 2020; Geribaldi-Doldán et al. 2019; Morita and Terada 2015; Murray and Fields 1998; Nie et al. 2010) |
| PG424 | Mitochondrial_lipids | (Horvath and Daum 2013; Morita and Terada 2015) | Neuroprotectant; Mitochondrial bioenergentics - ATP synthesis; Enhanced hippocampal neurogenesis - improve cognitive function | (Chen et al. 2018; Crotty et al. 2008; Donoso et al. 2020; Geribaldi-Doldán et al. 2019; Morita and Terada 2015; Murray and Fields 1998; Nie et al. 2010) |
| PG425 | Mitochondrial_lipids | (Horvath and Daum 2013; Morita and Terada 2015) | Neuroprotectant; Mitochondrial bioenergentics - ATP synthesis; Enhanced hippocampal neurogenesis - improve cognitive function | (Chen et al. 2018; Crotty et al. 2008; Donoso et al. 2020; Geribaldi-Doldán et al. 2019; Morita and Terada 2015; Murray and Fields 1998; Nie et al. 2010) |
| PG4412 | euronal_cell_body_lipi | (Horvath and Daum 2013; Morita and Terada 2015) | Neuroprotectant; Mitochondrial bioenergentics - ATP synthesis; Enhanced hippocampal neurogenesis - improve cognitive function | (Chen et al. 2018; Crotty et al. 2008; Donoso et al. 2020; Geribaldi-Doldán et al. 2019; Morita and Terada 2015; Murray and Fields 1998; Nie et al. 2010) |
| PG445 | Mitochondrial_lipids | (Horvath and Daum 2013; Morita and Terada 2015) | Neuroprotectant; Mitochondrial bioenergentics - ATP synthesis; Enhanced hippocampal neurogenesis - improve cognitive function | (Chen et al. 2018; Crotty et al. 2008; Donoso et al. 2020; Geribaldi-Doldán et al. 2019; Morita and Terada 2015; Murray and Fields 1998; Nie et al. 2010) |
| PG447 | Mitochondrial_lipids | (Horvath and Daum 2013; Morita and Terada 2015) | Neuroprotectant; Mitochondrial bioenergentics - ATP synthesis; Enhanced hippocampal neurogenesis - improve cognitive function | (Chen et al. 2018; Crotty et al. 2008; Donoso et al. 2020; Geribaldi-Doldán et al. 2019; Morita and Terada 2015; Murray and Fields 1998; Nie et al. 2010) |
| PG448 | Mitochondrial_lipids | (Horvath and Daum 2013; Morita and Terada 2015) | Neuroprotectant; Mitochondrial bioenergentics - ATP synthesis; Enhanced hippocampal neurogenesis - improve cognitive function | (Chen et al. 2018; Crotty et al. 2008; Donoso et al. 2020; Geribaldi-Doldán et al. 2019; Morita and Terada 2015; Murray and Fields 1998; Nie et al. 2010) |
| PG4610 | Mitochondrial_lipids | (Horvath and Daum 2013; Morita and Terada 2015) | Neuroprotectant; Mitochondrial bioenergentics - ATP synthesis; Enhanced hippocampal neurogenesis - improve cognitive function | (Chen et al. 2018; Crotty et al. 2008; Donoso et al. 2020; Geribaldi-Doldán et al. 2019; Morita and Terada 2015; Murray and Fields 1998; Nie et al. 2010) |
| PG469 | Mitochondrial_lipids | (Horvath and Daum 2013; Morita and Terada 2015) | Neuroprotectant; Mitochondrial bioenergentics - ATP synthesis; Enhanced hippocampal neurogenesis - improve cognitive function | (Chen et al. 2018; Crotty et al. 2008; Donoso et al. 2020; Geribaldi-Doldán et al. 2019; Morita and Terada 2015; Murray and Fields 1998; Nie et al. 2010) |
| PGO446 | Myelin_lipids | (Fitzner et al. 2020) | Alteration of membrane fluidity; Neurotransmitter signalling; Signal transduction; ER stress protection; Regulators of membrane trafficking and dynamics | (Hahnel et al. 1999; Messias et al. 2018; Wanders and Brites 2010) |
| PGO447 | Myelin_lipids | (Fitzner et al. 2020) | Alteration of membrane fluidity; Neurotransmitter signalling; Signal transduction; ER stress protection; Regulators of membrane trafficking and dynamics | (Hahnel et al. 1999; Messias et al. 2018; Wanders and Brites 2010) |
| PGP448 | Myelin_lipids | (Fitzner et al. 2020) | Alteration of membrane fluidity; Neurotransmitter signalling; Signal transduction; ER stress protection; Regulators of membrane trafficking and dynamics | (Hahnel et al. 1999; Messias et al. 2018; Wanders and Brites 2010) |
| PI340 | Synaptic_lipids | (Frere, Chang-Ileto, and Di Paolo 2012; Lauwers, Goodchild, and Verstreken 2016) | Neurotransmitter signalling; Promotes cell survival; Regulators of membrane trafficking and dynamics; Increase in dendritic spine density; Synaptic plasticity through induction of Long-term potentiation (LTP) | (Cosker et al. 2008; Cremona et al. 1999; Di Paolo and De Camilli 2006; Dickson and Hille 2019; Frere, Chang-Ileto, and Di Paolo 2012; Hossain et al. 2013; Dotti, Esteban, and Ledesma 2014; Lauwers, Goodchild, and Verstreken 2016; Puchkov and Haucke 2013; Raghu et al. 2019; Roy et al. 2014) |
| PI361 | Neuronal_cell_body | (Kaya et al. 2017) | Neurotransmitter signalling; Promotes cell survival; Regulators of membrane trafficking and dynamics; Increase in dendritic spine density; Synaptic plasticity through induction of Long-term potentiation (LTP) | (Cosker et al. 2008; Cremona et al. 1999; Di Paolo and De Camilli 2006; Dickson and Hille 2019; Frere, Chang-Ileto, and Di Paolo 2012; Hossain et al. 2013; Dotti, Esteban, and Ledesma 2014; Lauwers, Goodchild, and Verstreken 2016; Puchkov and Haucke 2013; Raghu et al. 2019; Roy et al. 2014) |
| PI364 | Neuronal_cell_body | (Kaya et al. 2017) | Neurotransmitter signalling; Promotes cell survival; Regulators of membrane trafficking and dynamics; Increase in dendritic spine density; Synaptic plasticity through induction of Long-term potentiation (LTP) | (Cosker et al. 2008; Cremona et al. 1999; Di Paolo and De Camilli 2006; Dickson and Hille 2019; Frere, Chang-Ileto, and Di Paolo 2012; Hossain et al. 2013; Dotti, Esteban, and Ledesma 2014; Lauwers, Goodchild, and Verstreken 2016; Puchkov and Haucke 2013; Raghu et al. 2019; Roy et al. 2014) |
| PI374 | Synaptic_lipids | (Frere, Chang-Ileto, and Di Paolo 2012; Lauwers, Goodchild, and Verstreken 2016) | Neurotransmitter signalling; Promotes cell survival; Regulators of membrane trafficking and dynamics; Increase in dendritic spine density; Synaptic plasticity through induction of Long-term potentiation (LTP) | (Cosker et al. 2008; Cremona et al. 1999; Di Paolo and De Camilli 2006; Dickson and Hille 2019; Frere, Chang-Ileto, and Di Paolo 2012; Hossain et al. 2013; Dotti, Esteban, and Ledesma 2014; Lauwers, Goodchild, and Verstreken 2016; Puchkov and Haucke 2013; Raghu et al. 2019; Roy et al. 2014) |

| | | | | |
|---|---|---|---|---|
| PI384 | Neuronal_cell_body | (Takamori et al. 2006; Merrill et al. 2017) | Neurotransmitter signalling; Promotes cell survival; Regulators of membrane trafficking and dynamics; Increase in dendritic spine density; Synaptic plasticity through induction of Long-term potentiation (LTP) | (Cosker et al. 2008; Cremona et al. 1999; Di Paolo and De Camilli 2006; Dickson and Hille 2019; Frere, Chang-Ileto, and Di Paolo 2012; Hossain et al. 2013; Dotti, Esteban, and Ledesma 2014; Lauwers, Goodchild, and Verstreken 2016; Puchkov and Haucke 2013; Raghu et al. 2019; Roy et al. 2014) |
| PI385 | Neuronal_cell_body | (Merrill et al. 2017; Tian et al. 2019) | Neurotransmitter signalling; Promotes cell survival; Regulators of membrane trafficking and dynamics; Increase in dendritic spine density; Synaptic plasticity through induction of Long-term potentiation (LTP) | (Cosker et al. 2008; Cremona et al. 1999; Di Paolo and De Camilli 2006; Dickson and Hille 2019; Frere, Chang-Ileto, and Di Paolo 2012; Hossain et al. 2013; Dotti, Esteban, and Ledesma 2014; Lauwers, Goodchild, and Verstreken 2016; Puchkov and Haucke 2013; Raghu et al. 2019; Roy et al. 2014) |
| PI386 | Synaptic_lipids | (Takamori et al. 2006; Frere, Chang-Ileto, and Di Paolo 2012; Lauwers, Goodchild, and Verstreken 2016) | Neurotransmitter signalling; Promotes cell survival; Regulators of membrane trafficking and dynamics; Increase in dendritic spine density; Synaptic plasticity through induction of Long-term potentiation (LTP) | (Cosker et al. 2008; Cremona et al. 1999; Di Paolo and De Camilli 2006; Dickson and Hille 2019; Frere, Chang-Ileto, and Di Paolo 2012; Hossain et al. 2013; Dotti, Esteban, and Ledesma 2014; Lauwers, Goodchild, and Verstreken 2016; Puchkov and Haucke 2013; Raghu et al. 2019; Roy et al. 2014) |
| PI405 | Synaptic_lipids | (Frere, Chang-Ileto, and Di Paolo 2012; Lauwers, Goodchild, and Verstreken 2016) | Neurotransmitter signalling; Promotes cell survival; Regulators of membrane trafficking and dynamics; Increase in dendritic spine density; Synaptic plasticity through induction of Long-term potentiation (LTP) | (Cosker et al. 2008; Cremona et al. 1999; Di Paolo and De Camilli 2006; Dickson and Hille 2019; Frere, Chang-Ileto, and Di Paolo 2012; Hossain et al. 2013; Dotti, Esteban, and Ledesma 2014; Lauwers, Goodchild, and Verstreken 2016; Puchkov and Haucke 2013; Raghu et al. 2019; Roy et al. 2014) |
| PI406 | Neuronal_cell_body | (Takamori et al. 2006) | Neurotransmitter signalling; Promotes cell survival; Regulators of membrane trafficking and dynamics; Increase in dendritic spine density; Synaptic plasticity through induction of Long-term potentiation (LTP) | (Cosker et al. 2008; Cremona et al. 1999; Di Paolo and De Camilli 2006; Dickson and Hille 2019; Frere, Chang-Ileto, and Di Paolo 2012; Hossain et al. 2013; Dotti, Esteban, and Ledesma 2014; Lauwers, Goodchild, and Verstreken 2016; Puchkov and Haucke 2013; Raghu et al. 2019; Roy et al. 2014) |
| PICer361 | Myelin_lipids | (Björkbom et al. 2010) | Neurotransmitter signalling; Promotes cell survival; Regulators of membrane trafficking and dynamics; Increase in dendritic spine density; Axonal growth; Synaptic plasticity through induction of Long-term potentiation (LTP) | (Björkbom et al. 2010; Dotti, Esteban, and Ledesma 2014; Gronnier et al. 2016; Henry, Gaspar, and Jesch 2014; Levine, Wiggins, and Munro 2000) |
| PICer401 | Myelin_lipids | (Björkbom et al. 2010) | Neurotransmitter signalling; Promotes cell survival; Regulators of membrane trafficking and dynamics; Increase in dendritic spine density; Axonal growth; Synaptic plasticity through induction of Long-term potentiation (LTP) | (Björkbom et al. 2010; Dotti, Esteban, and Ledesma 2014; Gronnier et al. 2016; Henry, Gaspar, and Jesch 2014; Levine, Wiggins, and Munro 2000) |
| PICer423 | Myelin_lipids | (Björkbom et al. 2010) | Neurotransmitter signalling; Promotes cell survival; Regulators of membrane trafficking and dynamics; Increase in dendritic spine density; Axonal growth; Synaptic plasticity through induction of Long-term potentiation (LTP) | (Björkbom et al. 2010; Dotti, Esteban, and Ledesma 2014; Gronnier et al. 2016; Henry, Gaspar, and Jesch 2014; Levine, Wiggins, and Munro 2000) |
| PIO341 | Myelin_lipids | (Björkbom et al. 2010) | Neurotransmitter signalling; Promotes cell survival; Regulators of membrane trafficking and dynamics; Increase in dendritic spine density; Axonal growth; Synaptic plasticity through induction of Long-term potentiation (LTP) | (Björkbom et al. 2010; Dotti, Esteban, and Ledesma 2014; Gronnier et al. 2016; Henry, Gaspar, and Jesch 2014; Levine, Wiggins, and Munro 2000) |
| PIP2340 | Synaptic_lipids | (Waugh 2015) | Neurotransmitter signalling; Promotes cell survival; Regulators of membrane trafficking and dynamics; Increase in dendritic spine density; Synaptic plasticity through induction of Long-term potentiation (LTP) | (Dotti, Esteban, and Ledesma 2014; Waugh 2015) |
| PIP2491 | Synaptic_lipids | (Waugh 2015) | Neurotransmitter signalling; Promotes cell survival; Regulators of membrane trafficking and dynamics; Increase in dendritic spine density; Synaptic plasticity through induction of Long-term potentiation (LTP) | (Dotti, Esteban, and Ledesma 2014; Waugh 2015) |
| PIP2492 | Synaptic_lipids | (Waugh 2015) | Neurotransmitter signalling; Promotes cell survival; Regulators of membrane trafficking and dynamics; Increase in dendritic spine density; Synaptic plasticity through induction of Long-term potentiation (LTP) | (Dotti, Esteban, and Ledesma 2014; Waugh 2015) |
| PI4210 | Synaptic_lipids | (Waugh 2015) | Neurotransmitter signalling; Promotes cell survival; Regulators of membrane trafficking and dynamics; Increase in dendritic spine density; Synaptic plasticity through induction of Long-term potentiation (LTP) | (Dotti, Esteban, and Ledesma 2014; Waugh 2015) |
| PIP384 | Synaptic_lipids | (Waugh 2015) | Neurotransmitter signalling; Promotes cell survival; Regulators of membrane trafficking and dynamics; Increase in dendritic spine density; Synaptic plasticity through induction of Long-term potentiation (LTP) | (Dotti, Esteban, and Ledesma 2014; Waugh 2015) |
| PIP393 | Synaptic_lipids | (Waugh 2015) | Neurotransmitter signalling; Promotes cell survival; Regulators of membrane trafficking and dynamics; Increase in dendritic spine density; Synaptic plasticity through induction of Long-term potentiation (LTP) | (Dotti, Esteban, and Ledesma 2014; Waugh 2015) |
| PIP407 | Synaptic_lipids | (Waugh 2015) | Neurotransmitter signalling; Promotes cell survival; Regulators of membrane trafficking and dynamics; Increase in dendritic spine density; Synaptic plasticity through induction of Long-term potentiation (LTP) | (Dotti, Esteban, and Ledesma 2014; Waugh 2015) |
| PIP416 | Synaptic_lipids | (Waugh 2015) | Neurotransmitter signalling; Promotes cell survival; Regulators of membrane trafficking and dynamics; Increase in dendritic spine density; Synaptic plasticity through induction of Long-term potentiation (LTP) | (Dotti, Esteban, and Ledesma 2014; Waugh 2015) |
| PIP443 | Synaptic_lipids | (Waugh 2015) | Neurotransmitter signalling; Promotes cell survival; Regulators of membrane trafficking and dynamics; Increase in dendritic spine density; Synaptic plasticity through induction of Long-term potentiation (LTP) | (Dotti, Esteban, and Ledesma 2014; Waugh 2015) |
| PIP597 | Synaptic_lipids | (Waugh 2015) | Neurotransmitter signalling; Promotes cell survival; Regulators of membrane trafficking and dynamics; Increase in dendritic spine density; Synaptic plasticity through induction of Long-term potentiation (LTP) | (Dotti, Esteban, and Ledesma 2014; Waugh 2015) |

| | | | | |
|---|---|---|---|---|
| PIP453 | Synaptic_lipids | (Waugh 2015) | Neurotransmitter signalling; Promotes cell survival; Regulators of membrane trafficking and dynamics; Increase in dendritic spine density; Synaptic plasticity through induction of Long-term potentiation (LTP) | (Dotti, Esteban, and Ledesma 2014; Waugh 2015) |
| PS341 | Axonal_lipids | (Calderon, Attema, and DeVries 2002; Kim, Huang, and Spector 2014) | Neuroprotectant via promotion of axonal regeneration; Induces ER-stress mediated apoptotic death; Neuronal proliferation; Axon-glial Interaction; Alteration of membrane fluidity; Promotes axonal regeneration; Increase in dendritic spine density ; Enhanced hippocampal neurogenesis - improve cognitive function; Synaptic plasticity through induction of Long-term potentiation (LTP); Increase in dendritic spine density; Activation of brain derived neurotrophic factor signaling cascade.; Axonal growth | (Nunzi et al. 1987; Arai et al. 2015; Calderon, Attema, and DeVries 2002; Nolan et al. 2004; Xu et al. 2005; Jiang et al. 2009; Kim, Akbar, and Kim 2010; Donoso et al. 2020; Fadok et al. 1998; Hisamoto et al. 2018; Kim, Huang, and Spector 2014; Vakhapova et al. 2014) |
| PS362 | Axonal_lipids | (Calderon, Attema, and DeVries 2002; Kim, Huang, and Spector 2014) | Neuroprotectant via promotion of axonal regeneration; Induces ER-stress mediated apoptotic death; Neuronal proliferation; Axon-glial Interaction; Alteration of membrane fluidity; Promotes axonal regeneration; Increase in dendritic spine density ; Enhanced hippocampal neurogenesis - improve cognitive function; Synaptic plasticity through induction of Long-term potentiation (LTP); Increase in dendritic spine density; Activation of brain derived neurotrophic factor signaling cascade.; Axonal growth | (Nunzi et al. 1987; Arai et al. 2015; Calderon, Attema, and DeVries 2002; Nolan et al. 2004; Xu et al. 2005; Jiang et al. 2009; Kim, Akbar, and Kim 2010; Donoso et al. 2020; Fadok et al. 1998; Hisamoto et al. 2018; Kim, Huang, and Spector 2014; Vakhapova et al. 2014) |
| PS384 | Axonal_lipids | (Calderon, Attema, and DeVries 2002; Kim, Huang, and Spector 2014; Arai et al. 2015; Donoso et al. 2020; Fadok et al. 1998; Hisamoto et al. 2018; Vakhapova et al. 2014; Merrill et al. 2017) | Neuroprotectant via promotion of axonal regeneration; Induces ER-stress mediated apoptotic death; Neuronal proliferation; Axon-glial Interaction; Alteration of membrane fluidity; Promotes axonal regeneration; Increase in dendritic spine density ; Enhanced hippocampal neurogenesis - improve cognitive function; Synaptic plasticity through induction of Long-term potentiation (LTP); Increase in dendritic spine density; Activation of brain derived neurotrophic factor signaling cascade.; Axonal growth | (Nunzi et al. 1987; Arai et al. 2015; Calderon, Attema, and DeVries 2002; Nolan et al. 2004; Xu et al. 2005; Jiang et al. 2009; Kim, Akbar, and Kim 2010; Donoso et al. 2020; Fadok et al. 1998; Hisamoto et al. 2018; Kim, Huang, and Spector 2014; Vakhapova et al. 2014) |
| PS404 | Axonal_lipids | (Calderon, Attema, and DeVries 2002; Kim, Huang, and Spector 2014; Arai et al. 2015; Donoso et al. 2020; Fadok et al. 1998; Hisamoto et al. 2018; Vakhapova et al. 2014; Merrill et al. 2017) | Neuroprotectant via promotion of axonal regeneration; Induces ER-stress mediated apoptotic death; Neuronal proliferation; Axon-glial Interaction; Alteration of membrane fluidity; Promotes axonal regeneration; Increase in dendritic spine density ; Enhanced hippocampal neurogenesis - improve cognitive function; Synaptic plasticity through induction of Long-term potentiation (LTP); Increase in dendritic spine density; Activation of brain derived neurotrophic factor signaling cascade.; Axonal growth | (Nunzi et al. 1987; Arai et al. 2015; Calderon, Attema, and DeVries 2002; Nolan et al. 2004; Xu et al. 2005; Jiang et al. 2009; Kim, Akbar, and Kim 2010; Donoso et al. 2020; Fadok et al. 1998; Hisamoto et al. 2018; Kim, Huang, and Spector 2014; Vakhapova et al. 2014) |
| PS406 | Axonal_lipids | (Calderon, Attema, and DeVries 2002; Kim, Huang, and Spector 2014; Arai et al. 2015; Donoso et al. 2020; Fadok et al. 1998; Hisamoto et al. 2018; Vakhapova et al. 2014; Takamori et al. 2006) | Neuroprotectant via promotion of axonal regeneration; Induces ER-stress mediated apoptotic death; Neuronal proliferation; Axon-glial Interaction; Alteration of membrane fluidity; Promotes axonal regeneration; Increase in dendritic spine density ; Enhanced hippocampal neurogenesis - improve cognitive function; Synaptic plasticity through induction of Long-term potentiation (LTP); Increase in dendritic spine density; Activation of brain derived neurotrophic factor signaling cascade.; Axonal growth | (Nunzi et al. 1987; Arai et al. 2015; Calderon, Attema, and DeVries 2002; Nolan et al. 2004; Xu et al. 2005; Jiang et al. 2009; Kim, Akbar, and Kim 2010; Donoso et al. 2020; Fadok et al. 1998; Hisamoto et al. 2018; Kim, Huang, and Spector 2014; Vakhapova et al. 2014) |
| PS407 | Axonal_lipids | (Calderon, Attema, and DeVries 2002; Kim, Huang, and Spector 2014; Merrill et al. 2017) | Neuroprotectant via promotion of axonal regeneration; Induces ER-stress mediated apoptotic death; Neuronal proliferation; Axon-glial Interaction; Alteration of membrane fluidity; Promotes axonal regeneration; Increase in dendritic spine density ; Enhanced hippocampal neurogenesis - improve cognitive function; Synaptic plasticity through induction of Long-term potentiation (LTP); Increase in dendritic spine density; Activation of brain derived neurotrophic factor signaling cascade.; Axonal growth | (Nunzi et al. 1987; Arai et al. 2015; Calderon, Attema, and DeVries 2002; Nolan et al. 2004; Xu et al. 2005; Jiang et al. 2009; Kim, Akbar, and Kim 2010; Donoso et al. 2020; Fadok et al. 1998; Hisamoto et al. 2018; Kim, Huang, and Spector 2014; Vakhapova et al. 2014) |
| PS429 | Axonal_lipids | (Calderon, Attema, and DeVries 2002;Kim, Huang, and Spector 2014;Arai et al. 2015; Donoso et al. 2020; Fadok et al. 1998; Hisamoto et al. 2018; Vakhapova et al. 2014) | Neuroprotectant via promotion of axonal regeneration; Induces ER-stress mediated apoptotic death; Neuronal proliferation; Axon-glial Interaction; Alteration of membrane fluidity; Promotes axonal regeneration; Increase in dendritic spine density ; Enhanced hippocampal neurogenesis - improve cognitive function; Synaptic plasticity through induction of Long-term potentiation (LTP); Increase in dendritic spine density; Activation of brain derived neurotrophic factor signaling cascade.; Axonal growth | (Nunzi et al. 1987; Arai et al. 2015; Calderon, Attema, and DeVries 2002; Nolan et al. 2004; Xu et al. 2005; Jiang et al. 2009; Kim, Akbar, and Kim 2010; Donoso et al. 2020; Fadok et al. 1998; Hisamoto et al. 2018; Kim, Huang, and Spector 2014; Vakhapova et al. 2014) |

| | | | | |
|---|---|---|---|---|
| PS4412 | Axonal_lipids | (Calderon, Attema, and DeVries 2002; Guo et al. 2007;Kim, Huang, and Spector 2014;) | Neuroprotectant via promotion of axonal regeneration; Induces ER-stress mediated apoptotic death; Neuronal proliferation; Axon-glial Interaction; Alteration of membrane fluidity; Promotes axonal regeneration; Increase in dendritic spine density ; Enhanced hippocampal neurogenesis - improve cognitive function; Synaptic plasticity through induction of Long-term potentiation (LTP); Increase in dendritic spine density; Activation of brain derived neurotrophic factor signaling cascade.; Axonal growth | (Nunzi et al. 1987; Arai et al. 2015; Calderon, Attema, and DeVries 2002; Nolan et al. 2004; Xu et al. 2005; Jiang et al. 2009; Kim, Akbar, and Kim 2010; Donoso et al. 2020; Fadok et al. 1998; Hisamoto et al. 2018; Kim, Huang, and Spector 2014; Vakhapova et al. 2014) |
| PS443 | Axonal_lipids | (Calderon, Attema, and DeVries 2002;Kim, Huang, and Spector 2014; Arai et al. 2015; Donoso et al. 2020; Fadok et al. 1998; Hisamoto et al. 2018; Vakhapova et al. 2014) | Neuroprotectant via promotion of axonal regeneration; Induces ER-stress mediated apoptotic death; Neuronal proliferation; Axon-glial Interaction; Alteration of membrane fluidity; Promotes axonal regeneration; Increase in dendritic spine density ; Enhanced hippocampal neurogenesis - improve cognitive function; Synaptic plasticity through induction of Long-term potentiation (LTP); Increase in dendritic spine density; Activation of brain derived neurotrophic factor signaling cascade.; Axonal growth | (Nunzi et al. 1987; Arai et al. 2015; Calderon, Attema, and DeVries 2002; Nolan et al. 2004; Xu et al. 2005; Jiang et al. 2009; Kim, Akbar, and Kim 2010; Donoso et al. 2020; Fadok et al. 1998; Hisamoto et al. 2018; Kim, Huang, and Spector 2014; Vakhapova et al. 2014) |
| PS444 | Axonal_lipids | (Calderon, Attema, and DeVries 2002;Kim, Huang, and Spector 2014; Arai et al. 2015; Donoso et al. 2020; Fadok et al. 1998; Hisamoto et al. 2018; Vakhapova et al. 2014) | Neuroprotectant via promotion of axonal regeneration; Induces ER-stress mediated apoptotic death; Neuronal proliferation; Axon-glial Interaction; Alteration of membrane fluidity; Promotes axonal regeneration; Increase in dendritic spine density ; Enhanced hippocampal neurogenesis - improve cognitive function; Synaptic plasticity through induction of Long-term potentiation (LTP); Increase in dendritic spine density; Activation of brain derived neurotrophic factor signaling cascade.; Axonal growth | (Nunzi et al. 1987; Arai et al. 2015; Calderon, Attema, and DeVries 2002; Nolan et al. 2004; Xu et al. 2005; Jiang et al. 2009; Kim, Akbar, and Kim 2010; Donoso et al. 2020; Fadok et al. 1998; Hisamoto et al. 2018; Kim, Huang, and Spector 2014; Vakhapova et al. 2014) |
| PS466 | Axonal_lipids | (Calderon, Attema, and DeVries 2002;Kim, Huang, and Spector 2014; Arai et al. 2015; Donoso et al. 2020; Fadok et al. 1998; Hisamoto et al. 2018; Vakhapova et al. 2014) | Neuroprotectant via promotion of axonal regeneration; Induces ER-stress mediated apoptotic death; Neuronal proliferation; Axon-glial Interaction; Alteration of membrane fluidity; Promotes axonal regeneration; Increase in dendritic spine density ; Enhanced hippocampal neurogenesis - improve cognitive function; Synaptic plasticity through induction of Long-term potentiation (LTP); Increase in dendritic spine density; Activation of brain derived neurotrophic factor signaling cascade.; Axonal growth | (Nunzi et al. 1987; Arai et al. 2015; Calderon, Attema, and DeVries 2002; Nolan et al. 2004; Xu et al. 2005; Jiang et al. 2009; Kim, Akbar, and Kim 2010; Donoso et al. 2020; Fadok et al. 1998; Hisamoto et al. 2018; Kim, Huang, and Spector 2014; Vakhapova et al. 2014) |
| PS468 | Axonal_lipids | (Calderon, Attema, and DeVries 2002;Kim, Huang, and Spector 2014; Arai et al. 2015; Donoso et al. 2020; Fadok et al. 1998; Hisamoto et al. 2018; Vakhapova et al. 2014) | Neuroprotectant via promotion of axonal regeneration; Induces ER-stress mediated apoptotic death; Neuronal proliferation; Axon-glial Interaction; Alteration of membrane fluidity; Promotes axonal regeneration; Increase in dendritic spine density ; Enhanced hippocampal neurogenesis - improve cognitive function; Synaptic plasticity through induction of Long-term potentiation (LTP); Increase in dendritic spine density; Activation of brain derived neurotrophic factor signaling cascade.; Axonal growth | (Nunzi et al. 1987; Arai et al. 2015; Calderon, Attema, and DeVries 2002; Nolan et al. 2004; Xu et al. 2005; Jiang et al. 2009; Kim, Akbar, and Kim 2010; Donoso et al. 2020; Fadok et al. 1998; Hisamoto et al. 2018; Kim, Huang, and Spector 2014; Vakhapova et al. 2014) |
| PS486 | Axonal_lipids | (Calderon, Attema, and DeVries 2002;Kim, Huang, and Spector 2014; Arai et al. 2015; Donoso et al. 2020; Fadok et al. 1998; Hisamoto et al. 2018; Vakhapova et al. 2014) | Neuroprotectant via promotion of axonal regeneration; Induces ER-stress mediated apoptotic death; Neuronal proliferation; Axon-glial Interaction; Alteration of membrane fluidity; Promotes axonal regeneration; Increase in dendritic spine density ; Enhanced hippocampal neurogenesis - improve cognitive function; Synaptic plasticity through induction of Long-term potentiation (LTP); Increase in dendritic spine density; Activation of brain derived neurotrophic factor signaling cascade.; Axonal growth | (Nunzi et al. 1987; Arai et al. 2015; Calderon, Attema, and DeVries 2002; Nolan et al. 2004; Xu et al. 2005; Jiang et al. 2009; Kim, Akbar, and Kim 2010; Donoso et al. 2020; Fadok et al. 1998; Hisamoto et al. 2018; Kim, Huang, and Spector 2014; Vakhapova et al. 2014) |
| PS488 | Axonal_lipids | (Calderon, Attema, and DeVries 2002;Kim, Huang, and Spector 2014; Arai et al. 2015; Donoso et al. 2020; Fadok et al. 1998; Hisamoto et al. 2018; Vakhapova et al. 2014) | Neuroprotectant via promotion of axonal regeneration; Induces ER-stress mediated apoptotic death; Neuronal proliferation; Axon-glial Interaction; Alteration of membrane fluidity; Promotes axonal regeneration; Increase in dendritic spine density ; Enhanced hippocampal neurogenesis - improve cognitive function; Synaptic plasticity through induction of Long-term potentiation (LTP); Increase in dendritic spine density; Activation of brain derived neurotrophic factor signaling cascade.; Axonal growth | (Nunzi et al. 1987; Arai et al. 2015; Calderon, Attema, and DeVries 2002; Nolan et al. 2004; Xu et al. 2005; Jiang et al. 2009; Kim, Akbar, and Kim 2010; Donoso et al. 2020; Fadok et al. 1998; Hisamoto et al. 2018; Kim, Huang, and Spector 2014; Vakhapova et al. 2014) |
| PS489 | Axonal_lipids | (Calderon, Attema, and DeVries 2002;Kim, Huang, and Spector 2014; Arai et al. 2015; Donoso et al. 2020; Fadok et al. 1998; Hisamoto et al. 2018; Vakhapova et al. 2014) | Neuroprotectant via promotion of axonal regeneration; Induces ER-stress mediated apoptotic death; Neuronal proliferation; Axon-glial Interaction; Alteration of membrane fluidity; Promotes axonal regeneration; Increase in dendritic spine density ; Enhanced hippocampal neurogenesis - improve cognitive function; Synaptic plasticity through induction of Long-term potentiation (LTP); Increase in dendritic spine density; Activation of brain derived neurotrophic factor signaling cascade.; Axonal growth | (Nunzi et al. 1987; Arai et al. 2015; Calderon, Attema, and DeVries 2002; Nolan et al. 2004; Xu et al. 2005; Jiang et al. 2009; Kim, Akbar, and Kim 2010; Donoso et al. 2020; Fadok et al. 1998; Hisamoto et al. 2018; Kim, Huang, and Spector 2014; Vakhapova et al. 2014) |

| | | | | |
|---|---|---|---|---|
| PSO425 | Axonal_lipids | (Calderon, Attema, and DeVries 2002;Kim, Huang, and Spector 2014; Arai et al. 2015; Donoso et al. 2020; Fadok et al. 1998; Hisamoto et al. 2018; Vakhapova et al. 2014) | Neuroprotectant via promotion of axonal regeneration; Induces ER-stress mediated apoptotic death; Neuronal proliferation; Axon-glial Interaction; Alteration of membrane fluidity; Promotes axonal regeneration; Increase in dendritic spine density ; Enhanced hippocampal neurogenesis - improve cognitive function; Synaptic plasticity through induction of Long-term potentiation (LTP); Increase in dendritic spine density; Activation of brain derived neurotrophic factor signaling cascade. | (Nunzi et al. 1987; Arai et al. 2015; Calderon, Attema, and DeVries 2002; Nolan et al. 2004; Xu et al. 2005; Jiang et al. 2009; Kim, Akbar, and Kim 2010; Donoso et al. 2020; Fadok et al. 1998; Hisamoto et al. 2018; Kim, Huang, and Spector 2014; Vakhapova et al. 2014) |
| SHexCer386 | Myelin_lipids | (Fitzner et al. 2020; Ishizuka 1997; Jackman, Ishii, and Bansal 2009; Kaya et al. 2017) | Myelination of neurons; Oligodendrocyte differentiation; Neurotransmitter signalling; Axon-glial Interaction | (Fledrich et al. 2018; Baba and Ishibashi 2019; Jeon et al. 2008; Palavicini et al. 2016) |
| SHexCer401 | Myelin_lipids | (Fitzner et al. 2020; Ishizuka 1997; Jackman, Ishii, and Bansal 2009; Kaya et al. 2017) | Myelination of neurons; Oligodendrocyte differentiation; Neurotransmitter signalling; Axon-glial Interaction | (Fledrich et al. 2018; Baba and Ishibashi 2019; Jeon et al. 2008; Palavicini et al. 2016) |
| SHexCer402 | Myelin_lipids | (Fitzner et al. 2020; Ishizuka 1997; Jackman, Ishii, and Bansal 2009; Kaya et al. 2017) | Myelination of neurons; Oligodendrocyte differentiation; Neurotransmitter signalling; Axon-glial Interaction | (Fledrich et al. 2018; Baba and Ishibashi 2019; Jeon et al. 2008; Palavicini et al. 2016) |
| SHexCer411 | Myelin_lipids | (Fitzner et al. 2020; Ishizuka 1997; Jackman, Ishii, and Bansal 2009; Kaya et al. 2017) | Myelination of neurons; Oligodendrocyte differentiation; Neurotransmitter signalling; Axon-glial Interaction | (Fledrich et al. 2018; Baba and Ishibashi 2019; Jeon et al. 2008; Palavicini et al. 2016) |
| SHexCer412 | Myelin_lipids | (Fitzner et al. 2020; Ishizuka 1997; Jackman, Ishii, and Bansal 2009; Kaya et al. 2017) | Myelination of neurons; Oligodendrocyte differentiation; Neurotransmitter signalling; Axon-glial Interaction | (Fledrich et al. 2018; Baba and Ishibashi 2019; Jeon et al. 2008; Palavicini et al. 2016) |
| SHexCer421 | Myelin_lipids | (Fitzner et al. 2020; Ishizuka 1997; Jackman, Ishii, and Bansal 2009; Kaya et al. 2017) | Myelination of neurons; Oligodendrocyte differentiation; Neurotransmitter signalling; Axon-glial Interaction | (Fledrich et al. 2018; Baba and Ishibashi 2019; Jeon et al. 2008; Palavicini et al. 2016) |
| SHexCer422 | Myelin_lipids | (Fitzner et al. 2020; Ishizuka 1997; Jackman, Ishii, and Bansal 2009; Kaya et al. 2017) | Myelination of neurons; Oligodendrocyte differentiation; Neurotransmitter signalling; Axon-glial Interaction | (Fledrich et al. 2018; Baba and Ishibashi 2019; Jeon et al. 2008; Palavicini et al. 2016) |
| SHexCerd422 | Myelin_lipids | (Fitzner et al. 2020; Ishizuka 1997; Jackman, Ishii, and Bansal 2009; Kaya et al. 2017) | Myelination of neurons; Oligodendrocyte differentiation; Neurotransmitter signalling; Axon-glial Interaction | (Fledrich et al. 2018; Baba and Ishibashi 2019; Jeon et al. 2008; Palavicini et al. 2016) |
| SHexCer423 | Myelin_lipids | (Fitzner et al. 2020; Ishizuka 1997; Jackman, Ishii, and Bansal 2009; Kaya et al. 2017) | Myelination of neurons; Oligodendrocyte differentiation; Neurotransmitter signalling; Axon-glial Interaction | (Fledrich et al. 2018; Baba and Ishibashi 2019; Jeon et al. 2008; Palavicini et al. 2016) |
| SHexCer432 | Myelin_lipids | (Fitzner et al. 2020; Ishizuka 1997; Jackman, Ishii, and Bansal 2009; Kaya et al. 2017) | Myelination of neurons; Oligodendrocyte differentiation; Neurotransmitter signalling; Axon-glial Interaction | (Fledrich et al. 2018; Baba and Ishibashi 2019; Jeon et al. 2008; Palavicini et al. 2016) |
| SHexCer466 | Myelin_lipids | (Fitzner et al. 2020; Ishizuka 1997; Jackman, Ishii, and Bansal 2009; Kaya et al. 2017) | Myelination of neurons; Oligodendrocyte differentiation; Neurotransmitter signalling; Axon-glial Interaction | (Fledrich et al. 2018; Baba and Ishibashi 2019; Jeon et al. 2008; Palavicini et al. 2016) |
| ST361 | Myelin_lipids | (Fitzner et al. 2020; Ishizuka 1997; Jackman, Ishii, and Bansal 2009; Kaya et al. 2017) | Myelination of neurons; Oligodendrocyte differentiation; Neurotransmitter signalling; Axon-glial Interaction | (Fledrich et al. 2018; Baba and Ishibashi 2019; Jeon et al. 2008; Palavicini et al. 2016) |
| STd422 | Myelin_lipids | (Fitzner et al. 2020; Ishizuka 1997; Jackman, Ishii, and Bansal 2009; Kaya et al. 2017) | Myelination of neurons; Oligodendrocyte differentiation; Neurotransmitter signalling; Axon-glial Interaction | (Fledrich et al. 2018; Baba and Ishibashi 2019; Jeon et al. 2008; Palavicini et al. 2016) |
